# PCMT1 generates the C-terminal cyclic imide degron on CRBN substrates

**DOI:** 10.1101/2025.03.24.645050

**Authors:** Zhenguang Zhao, Wenqing Xu, Ethan Yang Feng, Shiyun Cao, Alba Hermoso-López, Pablo Peña-Vega, Hannah C. Lloyd, Abigail K. D. Porter, Manuel Guzmán, Ning Zheng, Christina M. Woo

## Abstract

The E3 ligase substrate adapter cereblon (CRBN), the primary target of clinical agents thalidomide and lenalidomide, recognizes endogenous substrates bearing the C-terminal cyclic imide modification. Although C-terminal cyclic imides can form spontaneously, an enzyme that regulates the formation of these modifications and thereby promotes a biological pathway connecting substrates to CRBN is unknown. Here, we report that protein carboxymethyltransferase (PCMT1) promotes formation of the C-terminal cyclic imide on C-terminal asparagine residues of CRBN substrates. PCMT1 and CRBN co-regulate the levels of metabolic enzymes glutamine synthetase (GLUL) and inorganic pyrophosphatase 1 (PPA1) in vitro, in cells, and in vivo, and this regulation is associated with the proepileptic phenotype of CRBN knockout mouse models. The discovery of an enzyme that regulates CRBN substrates through the C-terminal cyclic imide modification reveals a previously unknown biological pathway that is perturbed by thalidomide derivatives and provides a biochemical basis for the connection between multiple biological processes and CRBN.

## Introduction

Cereblon (CRBN) is a substrate adapter of the CRL4 E3 ligase complex and the primary target of thalidomide and lenalidomide,^1^ clinical therapeutics that engage the thalidomide-binding domain of CRBN and are used to treat hematopoietic malignancies.^2,3^ Thalidomide, lenalidomide, and their derivatives modulate the selection of protein substrates by CRBN to promote the proteasomal degradation of induced substrates, which partially underlie these compounds’ therapeutic efficacy^4–7^ or their off-target effects (**Figure 1a–b**).^8,9^ These ligands mimic the C-terminal cyclic imide degron, a post-translational modification that arises from the intramolecular cyclization of asparagine or glutamine residues (cN or cQ, respectively) in protein substrates that are then recognized by CRBN for ubiquitination and proteasomal degradation (**Figure 1a–b**).^10^ However, the native substrates of CRBN and the mechanisms by which C-terminal cyclic imides form on substrates remain poorly understood. Identification of cellular processes that generate the C-terminal cyclic imide degron may reveal regulatory pathways that are mediated by CRBN and are modulated by thalidomide and its derivatives.

**Figure 1.**
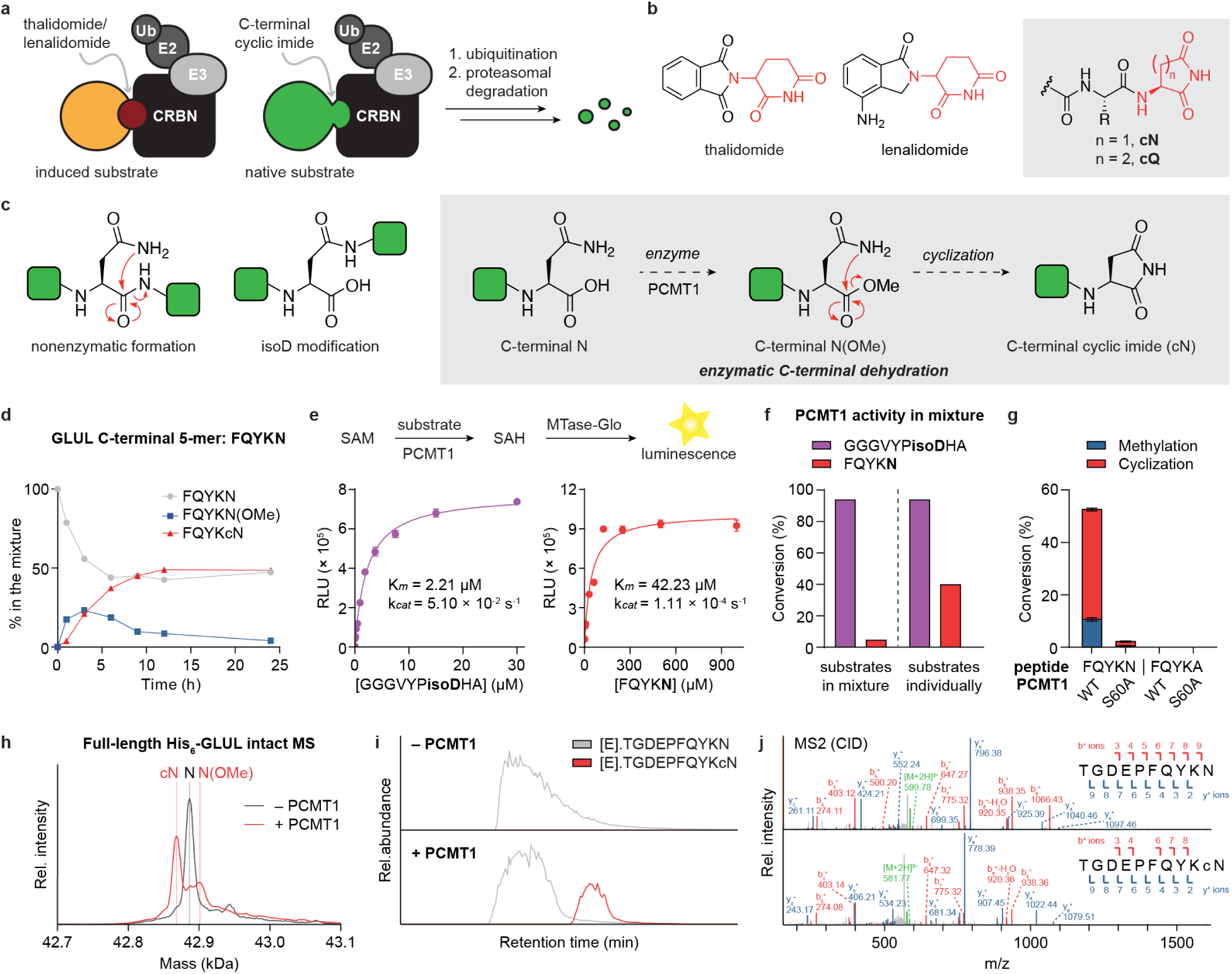
PCMT1 converts C-terminal Asn to C-terminal cyclic imide. **(a)** Scheme of the recognition of small molecule-induced or degron-mediated substrates by CRBN. **(b)** Ligands recognized by the thalidomide-binding domain of CRBN. **(c)** Structure of internal Asn, isoaspartate (isoD), and C-terminal Asn residues and proposed mechanism of enzyme-mediated C-terminal cyclic imide formation. **(d)** Time-course incubation of the peptide representing the C-terminus of GLUL with PCMT1 and SAM in 50 mM Tris-Cl, pH 7.4 buffer at 37 °C. **(e)** Comparison of the methyltransferase kinetics of PCMT1 for an isoD model substrate and the GLUL peptide by MTase-Glo assay. **(f)** Comparison of the catalytic activity of PCMT1 in a mixture of isoD and C-terminal N substrates and in each substrate individually (1 h incubation). **(g)** Conversion efficiency of the indicated peptides after 14 h incubation with wild-type or catalytically inactive PCMT1. **(h)** Intact mass spectra of full-length His_6_-GLUL after 14 h incubation with or without PCMT1. **(i)** Extracted ion chromatograms of the unmodified and cN-modified GLUL C-terminal peptides. Full-length His_6_-GLUL was treated with or without PCMT1 for 14 h and digested with GluC. **(j)** MS2 spectra of the uncyclized and cyclized GLUL C-terminal peptides.

The C-terminal cyclic imide modification can be generated spontaneously in eukaryotes as a form of protein damage from stress or aging, which occurs via intramolecular cleavage of the protein backbone primarily at asparagine residues (**Figure 1c**).^11,12^ Intramolecular cleavage from protein damage is mechanistically similar to deamidation, a form of protein damage that occurs primarily on asparagine and gives rise to an internal cyclic imide that, upon hydrolysis, affords an aspartate or isoaspartate (isoD) modification (**Figure 1c**, **EDF 1a**).^13,14^ The isoD modification requires correction to aspartate by the action of protein carboxymethyltransferase (PCMT1).^15–17^ PCMT1 is an S-adenosylmethionine (SAM)-dependent enzyme that methylates the isoD carboxylate to promote regeneration of the cyclic imide for the eventual disproportionation to aspartate, a less deleterious end product compared to isoD. Loss of PCMT1,^18^ like other enzymes that repair protein damage,^19^ remarkably impacts the brain, resulting in neurological disorders like seizures, which are reminiscent of the phenotypes that are genetically associated with loss-of-function mutations in CRBN in humans.^20,21^

The C-terminal cyclic imide degron could also conceivably be derived from an enzymatic source. Intriguingly, several native substrates previously connected to CRBN bear a C-terminal N or Q residue, pointing to a possible mechanism involving cyclization at the protein C-terminus to form a C-terminal cyclic imide (**Figure 1c**). These substrates, including glutamine synthetase (GLUL),^22^ APP,^23,24^ MEIS2,^25^ AMPK,^26^ and ClC-2,^27^ have been previously associated with CRBN, but whether that regulation occurs via the C-terminal cyclic imide degron is unknown. Further, an enzyme that tailors C-terminal N to a cyclic imide has not been previously annotated in eukaryotes. Thus, the identification of an enzyme that promotes C-terminal cyclic imide formation would reveal a pathway that unites the regulation of native CRBN substrates with the C-terminal cyclic imide degron.

## Results

### PCMT1 promotes C-terminal cyclic imide formation on GLUL

We first examined potential reactive intermediates that could be generated by an enzyme for their ability to promote formation of the C-terminal cyclic imide. Peptides with C-terminal N residues bearing methyl ester or thioester functional groups were monitored for their conversion to the C-terminal cyclic imide under physiological conditions (20 mM NH_4_OAc, pH 7.4 at 37 °C) by mass spectrometry (MS). While no conversion was observed on the free carboxylic acid, the methyl ester and thioester converted to the C-terminal cyclic imide at similar rates with a maximum conversion of 39% and 34%, respectively, within 24 h (**EDF 1b, Supplementary Table 1**). Therefore, activation of the C-terminal N carboxylate is required for the formation of the C-terminal cyclic imide, motivating us to explore biological mechanisms that generate one of these functional groups on the protein C-terminus.

Since PCMT1 is a carboxymethyltransferase with broad substrate recognition and the isoD modification is reminiscent of a C-terminal N, we investigated whether PCMT1 could activate the carboxylate on a C-terminal N to promote cyclic imide formation, which would be mechanistically analogous to PCMT1 activity on isoD (**Figure 1c**, **EDF 1c**). Incubation of recombinant PCMT1 with 5-mer peptides representing the C-termini of substrates previously connected to CRBN (GLUL, APP, MEIS2, AMPK α1, CIC-2) revealed the robust formation of the methyl ester and C-terminal cyclic imide products on peptides bearing C-terminal N, but not those bearing C-terminal Q, by mass spectrometry (**Figure 1d, EDF 2a**). In particular, the peptide representing the C-terminus of GLUL, FQYKN, showed an appreciable 49% formation of the C-terminal cyclic imide product, FQYKcN, after a 12 h incubation with PCMT1, despite contributions from background hydrolysis (**Figure 1d, EDF 1b**). Methylation was observed on the peptides bearing C-terminal Q, but subsequent formation of the C-terminal cQ was not detected. Further examination of PCMT1 activity on additional model peptides (Fmoc-FN, GGGFN) showed that PCMT1 minimally requires the acyclic C-terminal N to catalyze cN formation (**EDF 2b**). PCMT1 primarily promotes the generation of the methyl ester intermediate, after which cyclization is spontaneous, as confirmed by the comparable conversion of synthetic PFQYKN(OMe) peptide into PFQYKcN independent of PCMT1 or SAM (**EDF 2c**).

We selected the most reactive peptide, FQYKN, for deeper investigation. Using a methyltransferase glo (MTase-Glo) assay,^28^ the K_m_ and k_cat_ of PCMT1 for FQYKN were measured to be 42.23 µM and 1.11×10^−4^ s^-1^, respectively, which falls within the range of PCMT1 activity on various isoD-modified peptides, although it is approximately 20-fold less active than an optimal isoD peptide, GGGVYPisoDHA (**Figure 1e**).^29^ Both peptides engage the same active site on PCMT1, as the isoD peptide competitively inhibited the conversion of FQYKN in an equimolar mixture (**Figure 1f**). Methylation and formation of the C-terminal cyclic imide on PFQKYN were ablated with the inactive mutant PCMT1 S60A,^30^ or if the C-terminal N was mutated to alanine (**Figure 1g, EDF 2d**). These data demonstrate that PCMT1 selectively methylates C-terminal N to promote conversion to the C-terminal cyclic imide on the GLUL C-terminal 5-mer FQYKN.

We then asked if PCMT1 could enzymatically modify the full-length GLUL protein. Although GLUL was previously identified as a substrate of PCMT1, the specific molecular recognition event was not defined.^31^ We therefore incubated recombinant full-length GLUL with PCMT1 and SAM for 14 h and measured the formation of the cN-modified form of GLUL (GLUL-cN) by both intact MS and bottom-up proteomics. Intact MS revealed robust formation of GLUL-cN, as well as a smaller population of the C-terminal methylated product, upon incubation with PCMT1 (**Figure 1h**). Digestion of PCMT1-treated GLUL with the Glu-C protease confirmed the generation of the cyclic imide at the C-terminus of GLUL in a PCMT1-dependent manner, which is readily observed, despite likely hydrolysis of the cyclic imide during sample preparation (**Figure 1i–j**). Altogether, these data show that PCMT1 catalyzes the formation of the C-terminal cyclic imide on full-length GLUL in vitro.

### Sequence preference of PCMT1 activity and CRBN recognition

We next examined whether CRBN recognizes the C-terminal cyclic imide degron on the GLUL C-terminal peptide installed by PCMT1. To quantitatively measure binding to CRBN, we employed a time-resolved Förster resonance energy transfer (TR-FRET) assay using the His_6_-CRBN/DDB1 complex labeled with CoraFluor-1 as the FRET donor and assessed the ability of different species to competitively displace the thalidomide-fluorescein isothiocyanate (thal-FITC) tracer as the FRET acceptor (**Figure 2a, EDF 3a**).^32,33^ A tracer equilibrium dissociation constant (K_D_) of 93.7 nM for thal-FITC to the His_6_-CRBN/DDB1 complex was used to convert ligand IC_50_ to K_D_ values in this study.^12^ Using this assay, we first showed that a synthetic GLUL C-terminal 6-mer, PFQYKcN, binds CRBN with an affinity equivalent to lenalidomide. Moreover, when we incubated the acyclic PFQYKN peptide with PCMT1 for 14 h, the product engaged CRBN with equivalent affinity, but this binding was ablated if PCMT1 was omitted from the mixture (**Figure 2b, Supplementary Table 2**). These data validate CRBN’s recognition of the cyclic imide degron installed by PCMT1.

**Figure 2.**
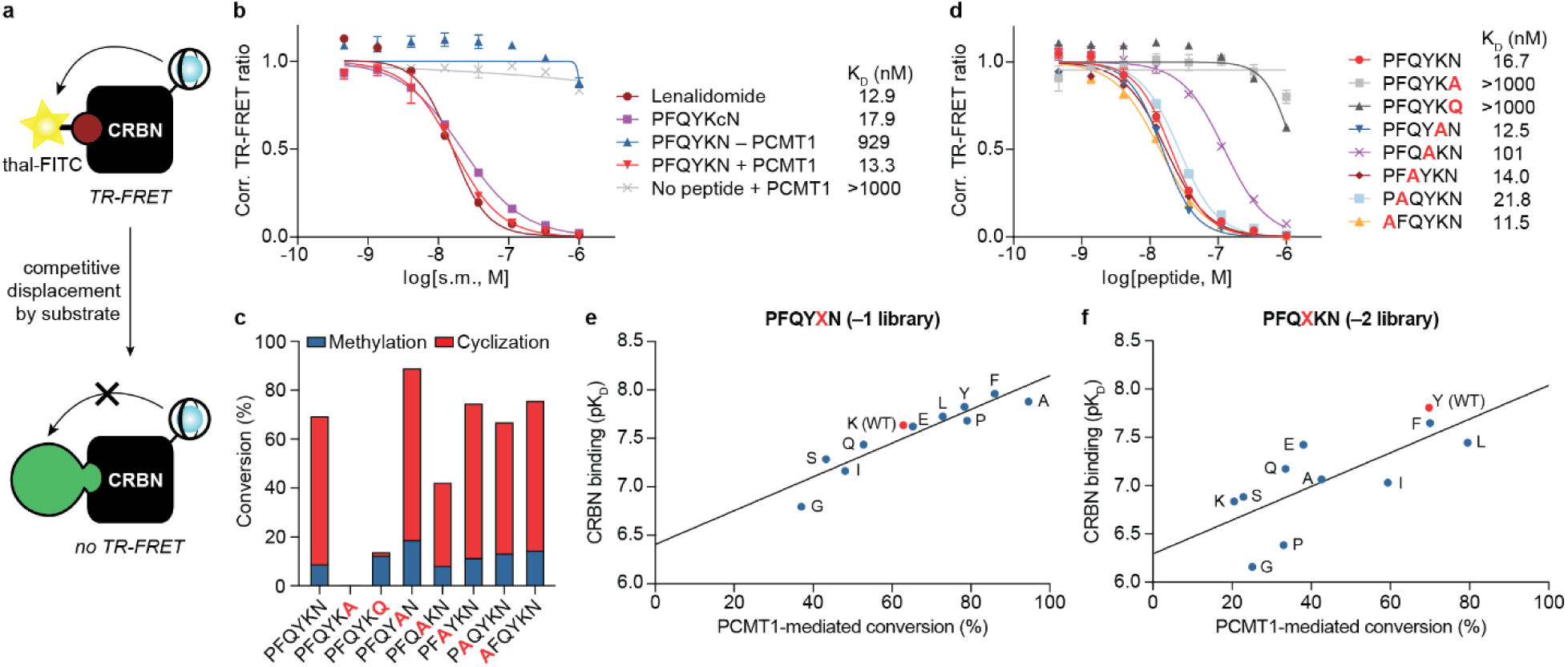
CRBN recognizes the C-terminal cyclic imide degron installed by PCMT1. **(a)** Schematic of the TR-FRET assay used to measure the binding of ligands to CRBN based on their ability to displace the tracer molecule. **(b)** TR-FRET assay of lenalidomide, PFQYKcN, and PFQYKN incubated with or without PCMT1 to measure their affinity to His_6_-CRBN/DDB1. **(c)** Conversion efficiency of GLUL peptides with single residue substitutions after 14 h incubation with PCMT1 by mass spectrometry. **(d)** TR-FRET assay of GLUL peptides with single residue substitutions after 14 h incubation with PCMT1 to measure their affinity to His_6_-CRBN/DDB1. **(e)** Correlation of % PCMT1-mediated conversion measured by mass spectrometry and pK_D_ values measured by TR-FRET assay in GLUL peptides with variable –1 residue. **(f)** Correlation of % PCMT1-mediated conversion measured by mass spectrometry and pK_D_ values measured by TR-FRET assay in GLUL peptides with variable –2 residue. All TR-FRET experiments were performed with 3 technical replicates, and data are presented as mean ± s.d. (n = 3).

We sought to evaluate the sensitivity of PCMT1 and CRBN to systematic variations in peptide length and sequence. GLUL C-terminal peptides of varying lengths were incubated with PCMT1, and their conversion and binding to CRBN were evaluated by MS and TR-FRET, respectively. The scope of PCMT1 activity in vitro is relatively flexible with different peptide lengths, as shown by the comparable MS conversion, whereas CRBN requires at least five amino acids for appreciable binding (**EDF 3b**). An alanine scan across the GLUL 6-mer validated the ablation of PCMT1 activity and CRBN recognition upon loss of the C-terminal N residue and additionally highlighted the importance of the residue at the –2 position on PCMT1 activity and CRBN engagement compared to the other residues upstream of the C-terminal N (**Figure 2c–d**).

To further profile the sequence preferences of both PCMT1 and CRBN recognition, two GLUL peptide libraries were examined, one with a variable –1 residue and the other with a variable –2 residue (**Figure 2e–f, EDF 3c–d**). At the –1 position, hydrophobic residues such as A, F, and Y moderately enhanced the recognition of both PCMT1 and CRBN relative to the native residue K (**Figure 2e, EDF 3c**). Varying the –2 residue produced greater divergence in MS conversion and K_D_ values between the residues, with Y (the WT residue), F, and L exhibiting the highest conversion by PCMT1 and strongest binding to CRBN (**Figure 2f, EDF 3d**). Conversion by PCMT1 largely correlated with subsequent CRBN binding, with the notable exception of the peptides bearing P or G at the –2 position, which had a weaker affinity to CRBN relative to other residues with comparable PCMT1-mediated methylation and cyclization, indicating that CRBN disfavors P and G residues at the –2 position (**Figure 2f**). Taken together, these data demonstrate a strong positive correlation between PCMT1 activity and CRBN binding for the examined GLUL-derived peptides, with some sequence preference at the –1 and –2 positions relative to the C-terminal N.

### GLUL is a substrate of PCMT1 and CRBN in vitro and in cells

We next investigated if the generation of GLUL-cN by PCMT1 affords a substrate of CRBN and results in binding, ubiquitination, and proteasomal degradation (**Figure 3a**). Full-length GST-GLUL incubated with PCMT1 displays robust binding to CRBN, which is abrogated if PCMT1 is omitted or if the C-terminal N of GLUL (N373) is mutated to alanine, indicating that GLUL is recognized by CRBN upon formation of the C-terminal cyclic imide by PCMT1 (**Figure 3b, Supplementary Table 3**). Treatment of GLUL with PCMT1 is both sufficient and necessary to induce CRL4^CRBN^-mediated ubiquitination in vitro (**Figure 3c**). To investigate whether GLUL-cN is degraded by CRBN in cells, we incubated recombinant GFP-GLUL with PCMT1, electroporated the product into HEK293T cells, then monitored GFP levels by flow cytometry. We observed a 70% decrease in PCMT1-treated GFP-GLUL after 6 h, in a manner that is dependent on the GLUL C-terminal N and PCMT1 treatment and is inhibited by lenalidomide (**Figure 3d**). Since GLUL exists as an oligomer, we verified that electroporation did not alter the oligomerization state of GFP-GLUL, which indicates that the electroporated protein was not affected by a separate quality control pathway (**EDF 4a**). Collectively, these data show that C-terminal cyclic imide generation by PCMT1 causes GLUL to become a CRBN substrate when introduced into cells.

**Figure 3.**
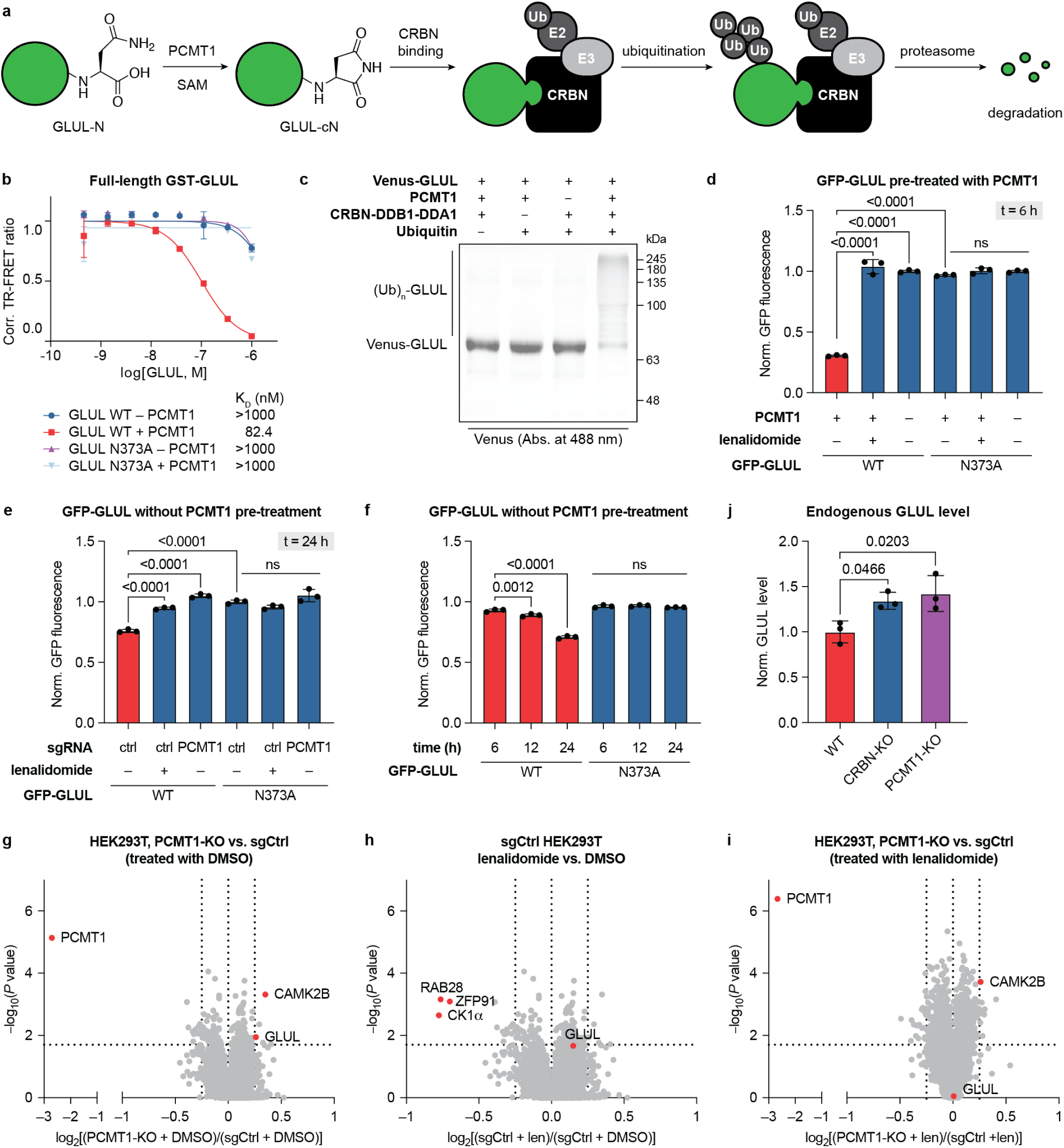
Co-regulation of GLUL by PCMT1 and CRBN in vitro and in cells. **(a)** Scheme of degron formation by PCMT1 and recognition by CRBN resulting in proteasomal degradation of GLUL. **(b)** TR-FRET assay of full-length GST-GLUL after 14 h incubation with or without PCMT1. **(c)** In vitro ubiquitination assay of Venus-GLUL with or without PCMT1. **(d)** Flow cytometry analysis of GFP levels in HEK293T cells 6 h after electroporation of GFP-GLUL protein that was pre-incubated with or without PCMT1 for 14 h. **(e)** Flow cytometry analysis of GFP levels in WT or PCMT1-KO HEK293T cells 24 h after electroporation of GFP-GLUL. **(f)** Time-course flow cytometry analysis of GFP levels in HEK293T cells 6–24 h after electroporation of GFP-GLUL. All flow cytometry experiments were performed with 3 biological replicates. Comparisons were performed using one-way ANOVA with Šidák’s multiple comparisons tests, and p-values are indicated. **(g–i)** Quantitative proteomics of sgCtrl or PCMT1-KO HEK293T cells after 48 h treatment with DMSO or 100 µM lenalidomide. The experiment was performed with 4 biological replicates. P-values for the abundance ratios were calculated by one-way ANOVA with TukeyHSD post-hoc test. **(j)** Quantification of endogenous GLUL levels in WT, PCMT1-KO, or CRBN-KO HEK293T cells by Western blot across 3 biological replicates. Comparisons were performed using a one-way ANOVA with Šidák’s multiple comparisons test, and p-values are indicated.

To investigate whether endogenous PCMT1 likewise promotes GLUL-cN formation for CRBN-dependent degradation, we directly electroporated unmodified GFP-GLUL into HEK293T cells with or without PCMT1 knockout (KO) and measured GFP levels after a longer incubation period. After 24 h, GFP-GLUL was reduced by 26% in HEK293T sgCtrl cells relative to GFP-GLUL(N373A), which was ablated by PCMT1 KO, analogously to the rescue from competitive inhibition of CRBN by lenalidomide (**Figure 3e**). GFP-GLUL levels gradually decrease over 24 h, which implies that the PCMT1-mediated cyclic imide formation is rate-limiting in the cellular degradation of GFP-GLUL (**Figure 3f**). These data establish that when introduced into cells, GLUL is recognized as a substrate of CRBN through the C-terminal cyclic imide degron generated by PCMT1.

### PCMT1 and CRBN regulate endogenous GLUL

To profile the endogenous substrates of PCMT1 and CRBN in cells, we performed global quantitative proteomics in HEK293T sgCtrl and PCMT1-KO cells treated with either DMSO or 100 µM lenalidomide for 48 h. We expected that substrates regulated by PCMT1 and CRBN would accumulate in PCMT1-KO cells or lenalidomide-treated cells relative to sgCtrl, but that there would be no apparent difference between PCMT1-KO and lenalidomide treatment since CRBN-mediated degradation of cN-bearing native substrates would be blocked in both conditions. Out of 7704 quantified proteins, we identified 10 that are significantly increased upon PCMT1-KO in the DMSO condition [log_2_(fold-change) > 0.25, p-value < 0.05]. Of these proteins, GLUL was the only protein with a C-terminal N residue that was upregulated upon PCMT1-KO or lenalidomide treatment, but not significantly changed when comparing these two conditions (**Figure 3g–i, Supplementary Table 4**). By contrast, for example, CAMK2B was upregulated upon PCMT1-KO, but not following lenalidomide treatment, indicating that its levels change in response to PCMT1 but not the thalidomide-binding domain of CRBN. Western blot analysis further confirmed the reproducible elevation of endogenous GLUL upon either PCMT1-KO or CRBN-KO in HEK293T cells (**Figure 3j, EDF 4b**). These data illustrate the regulation of endogenous GLUL by PCMT1 and CRBN in cells.

GLUL is the only enzyme capable of de novo synthesis of glutamine via an ATP-dependent condensation of glutamate and ammonium (**EDF 5a**).^34^ The cell regulates GLUL levels inversely proportionate to cellular glutamine concentrations to match the demand for de novo glutamine synthesis.^35^ A previous study reported that CRBN plays a role in glutamine-triggered GLUL degradation by recognizing glutamine-dependent acetylation of lysines on the N-terminus of GLUL.^22^ We thus investigated if the C-terminal cyclic imide modification installed by PCMT1 also participates in the glutamine-mediated regulation of GLUL. As expected, GLUL levels increased in HEK293T cells following glutamine starvation for 24–48 h (**EDF 5b**). However, glutamine refeeding with 4 mM glutamine resulted in time-dependent GLUL depletion that was independent of PCMT1 or CRBN (**EDF 5c**). The GLUL mRNA levels were identical in sgCtrl, PCMT1-KO, and CRBN-KO cells at steady glutamine concentrations, although the basal level shifted lower in response to glutamine starvation after 48 h (**EDF 5d**). These data imply that the regulation of GLUL by PCMT1 is not involved in the glutamine-triggered degradation of GLUL.

### CRBN ligands inhibit degradation of GLUL-cN

To investigate whether degradation of GLUL via the C-terminal cyclic imide is inhibited by CRBN ligands, we employed the cerebody method, which utilizes an engineered CRBN construct with improved stability in Western blot and affinity enrichment experiments, to enrich CRBN substrates like GLUL-cN from cell lysates (**Figure 4a**).^36^ The proportion of the cellular GLUL population bearing the C-terminal cyclic imide was clearly elevated after lenalidomide treatment or CRBN-KO but commuted in HEK293T PCMT1-KO cells, as shown by Western blot following cerebody enrichment (**Figure 4b, EDF 6a, Supplementary Table 5**). Importantly, no GLUL-cN is observed in the absence of PCMT1 suggesting that PCMT1 is the only enzyme capable of catalyzing C-terminal cyclic imide formation on GLUL in this cell type. Next, by performing selected ion monitoring (SIM) analysis for the cN-modified GLUL C-terminal peptide from HEK293T cell lysates after cerebody enrichment with a heavy internal standard, we verified the presence of the cyclic imide modification at the GLUL C-terminus only in lenalidomide-treated or CRBN-KO cells, but not in PCMT1-KO cells (**Figure 4c**). The levels of unmodified N peptide varied commensurately with the cN peptide, which could derive from hydrolysis of GLUL-cN during MS sample preparation or from unmodified GLUL monomers that are enriched in complex with GLUL-cN monomers. These data confirm that the generation of endogenous GLUL-cN is PCMT1-dependent and that its removal is blocked by loss or inhibition of CRBN.

**Figure 4.**
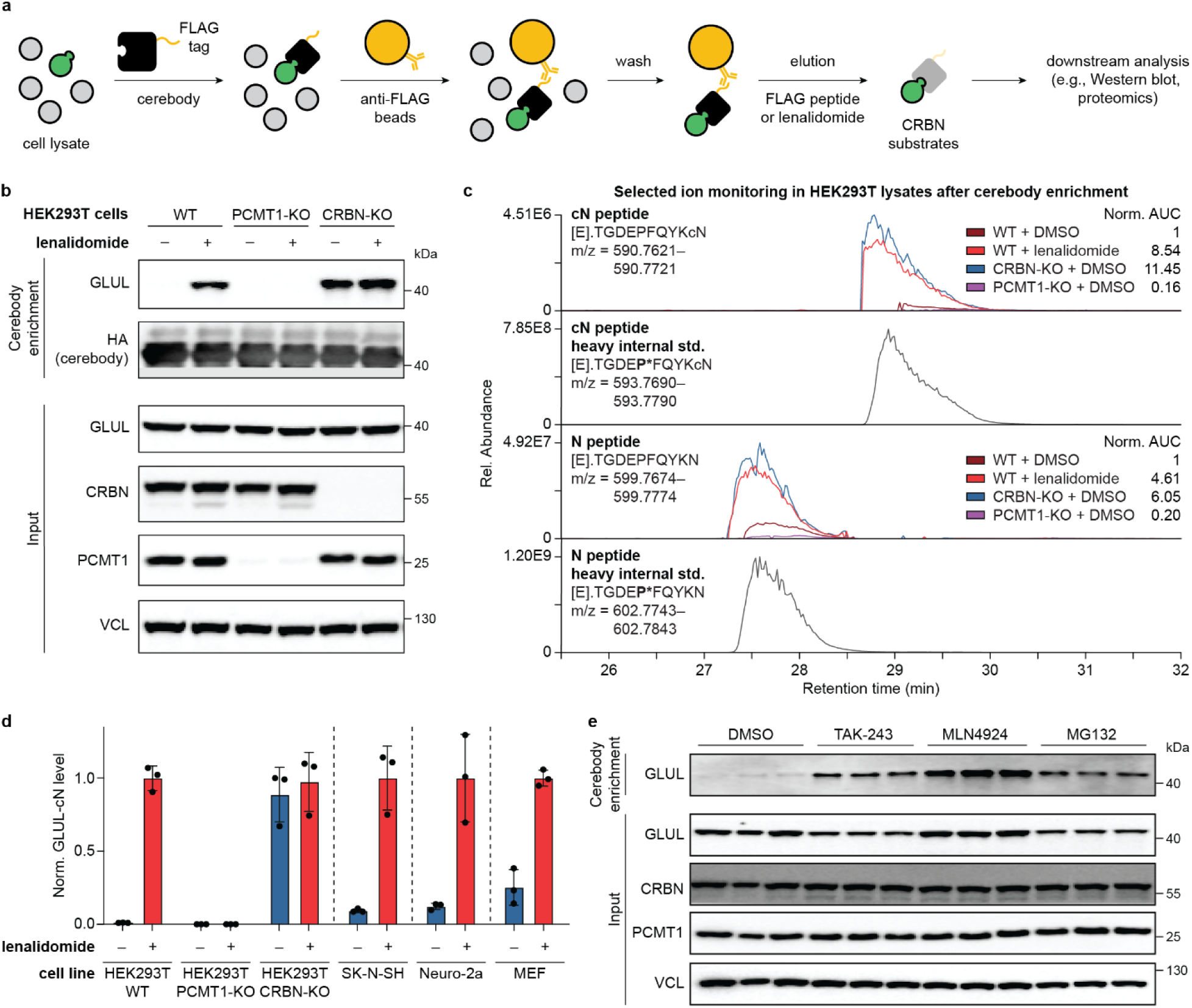
GLUL is an endogenous substrate of CRBN through the C-terminal cyclic imide degron that is impacted by clinical agents. **(a)** Scheme of workflow to enrich CRBN substrates for downstream analysis using cerebody. **(b)** Cerebody enrichment for GLUL-cN from WT, PCMT1-KO, or CRBN-KO HEK293T cells after treatment with DMSO or 100 µM lenalidomide for 24 h. **(c)** Selected ion monitoring of the cN-modified and unmodified GLUL C-terminal peptides after cerebody enrichment of WT, CRBN-KO, or PCMT1-KO HEK293T cells treated with DMSO or 100 µM lenalidomide for 48 h. Heavy standards labeled with proline-^13^C_5_,^15^N were spiked in to confirm the identities of the peptides of interest. (**d**) Endogenous GLUL-cN level in different cell lines after treatment with DMSO or 100 µM lenalidomide for 24 h, determined by cerebody enrichment. **(e)** Cerebody enrichment for GLUL-cN from Neuro-2a cells after treatment with DMSO, 5 µM TAK-243, 1 µM MLN2924, or 10 µM MG132 for 6 h.

We further evaluated the levels of GLUL-cN in several additional cell lines after inhibition of CRBN with 100 µM lenalidomide for 24 h, including the human neuroblastoma cell line SK-N-SH, the mouse neuronal cell line Neuro-2a, and the mouse embryonic fibroblast (MEF) cell line. GLUL-cN was readily enriched upon CRBN inhibition in all of these cell lines, demonstrating the activation of this pathway across a variety of cell lines and species (**Figure 4d**, **EDF 6b**). The accumulation of GLUL-cN in Neuro-2a cells following lenalidomide treatment was dose-dependent and plateaued by 24 h (**EDF 7a–b**). Moreover, inhibitors of the ubiquitin-proteasome pathway, including the E1 inhibitor TAK-243, neddylation inhibitor MLN4924, and proteasome inhibitor MG132, all rescued GLUL-cN in Neuro-2a cells, confirming the degradation of GLUL-cN by the ubiquitin-proteasome system under native conditions (**Figure 4e**). Together, these data demonstrate the bona fide regulatory mechanism of endogenous GLUL mediated by PCMT1 and CRBN through the C-terminal cyclic imide degron across several cell lines and species, which is competitively inhibited by CRBN-targeting clinical agents.

### Structural basis of GLUL-cN binding to CRBN

To reveal the mechanism of GLUL-cN recognition by CRBN and further map key structural determinants of the GLUL-cN degron, we solved a crystal structure of human CRBN-DDB1 in complex with the GLUL-cN C-terminal 6-mer peptide, PFQYKcN, at a 2.9 Å resolution (**Figure 5a, EDF 8a**, **Extended Data Table 1**). The C-terminal cN and the three upstream residues were well resolved with continuous density, whereas the remaining two residues of the 6-mer peptide were invisible, presumably due to their disordered nature (**EDF 8a**). As expected, the C-terminal cyclic imide is recognized by the tri-tryptophan cage in the thalidomide-binding domain of CRBN (**Figure 5b**), analogous to the binding mode of thalidomide derivatives to CRBN (**EDF 8b**). Single amino acid mutations of the W380, W386, and W400 residues in CRBN abolished its interaction with GLUL-cN (**Figure 5c**), indicating that the tri-tryptophan cage plays a critical role in recognizing the native substrate via the cyclic imide. Interestingly, the three residues in GLUL-cN upstream of cN also make contacts with CRBN (**Figure 5b**). Specifically, N351 in CRBN donates a hydrogen bond to the backbone carbonyl group of the –1 residue in GLUL-cN, while the tyrosine residue at the –2 position closely packs against H397 in CRBN. Additional van der Waals interactions are made between the peptide backbone of the GLUL-cN C-terminus and Y355 and H357 in CRBN. In support of this extended GLUL-cN-CRBN interface, individual mutations of the CRBN residues involved in these interactions impaired binding to GLUL-cN, as measured by biolayer interferometry (BLI, **Figure 5c–d, EDF 8c–g**). These data confirm the optimal geometry of the GLUL-cN peptide backbone at the –1 position and the involvement of the side chain at the –2 position when bound to CRBN.

**Figure 5.**
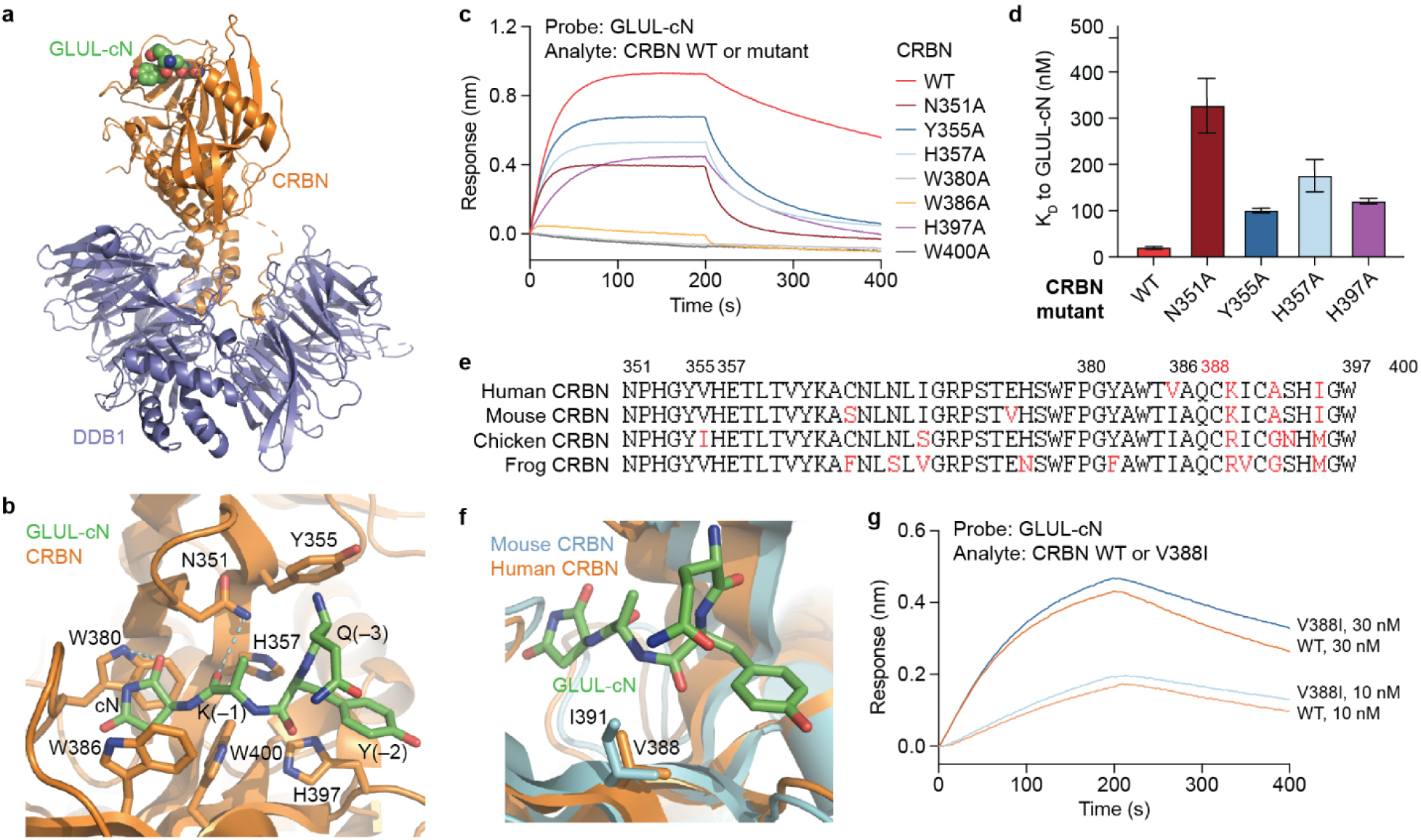
Structural basis of GLUL-cN binding to CRBN. **(a)** Crystal structure of the DDB1-CRBN-GLUL-cN complex. GLUL-cN is shown as green, red and blue spheres. DDB1 and CRBN are shown as cartoon diagrams and colored slate and orange, respectively. **(b)** Close-up view of CRBN residues interacting with GLUL-cN. GLUL-cN is shown as green sticks. The electron density for the side chain of lysine at the –1 position is incomplete (likely due to flexibility) and is thus not included in the model. CRBN is shown as a cartoon diagram with interface residues highlighted. The hydrogen bonds are represented by cyan dashes. **(c)** BLI measurements of GLUL-cN binding to CRBN WT or mutants. **(d)** The K_D_ values for GLUL-cN binding to CRBN WT or mutants, measured by BLI. **(e)** Sequence alignment of four vertebrate CRBN orthologues. Key residue numbers of human CRBN are labeled on top of the sequence. **(f)** Mouse CRBN (cyan, AlphaFold model) aligned with human CRBN-GLUL-cN complex. Human CRBN is colored orange. GLUL-cN is shown as green sticks. The side chains of human CRBN V388 and its counterpart in mouse, I391, are highlighted. **(g)** BLI measurements of GLUL-cN binding to the CRBN WT and V388I variant.

Residues N351, Y355, and H357, which are among the key GLUL-binding residues, are part of the CRBN sensor loop (341–361), whose conformation can be modulated by thalidomide derivatives.^37^ Upon binding CRBN, these ligands trigger a conformational change in CRBN from an open to a closed state, where stabilization of the closed conformation positively correlates with induced neosubstrate degradation outcomes. By contrast, GLUL-cN directly interacts with the CRBN sensor loop to promote stronger binding interactions with CRBN (**Figure 5b**). Therefore, the native substrate itself can modulate the sensor loop of CRBN, leading to efficient substrate recruitment and ubiquitination.

Importantly, sequence alignment reveals that both the C-terminal 3-mer of GLUL and all seven CRBN residues involved in recognizing GLUL-cN are strictly conserved from humans to amphibians (**Figure 5e, EDF 8h**). CRBN can be artificially rewired by thalidomide and lenalidomide to recruit neosubstrates for degradation in humans, but not in mice. This differential activity has been attributed to the V388 residue in human CRBN, which is replaced by isoleucine in mice and other species. This alteration clashes with neosubstrates and inhibits their binding in these species (**Figure 5e**).^38^ Considering that native substrates are more likely to be conserved between species, we anticipated that the mouse CRBN isoleucine mutation would not cause steric hinderance with GLUL-cN. Indeed, GLUL-cN is structurally compatible with an isoleucine residue at the V388 position in CRBN (**Figure 5f**), and a human CRBN V388I mutant retained robust binding to GLUL-cN (**Figure 5g**). These data suggest that the function of CRBN in recognizing native substrates bearing the C-terminal imide is conserved across species.

### Loss of CRBN function affects the brain

As dysregulation of CRBN,^20,39^ PCMT1,^18,40^ and GLUL^41^ are linked to negative neurological effects, including intellectual disability, learning and memory impairment, and seizures, we investigated whether the loss of CRBN results in accumulation of GLUL-cN in the brain. To specifically probe the consequences of CRBN loss in the brain, we performed a quantitative proteomics investigation of the hippocampus in a full Crbn^−/–^ mouse line.^39^ This brain region expresses high levels of CRBN^39^ and is primarily associated with learning and memory,^42^ as well as with the control of seizures.^43^ Comparison of the hippocampi from four WT and four Crbn^−/–^ mice (two male, two female) identified 6845 proteins. Of these, the two most significantly upregulated proteins, Glul and inorganic pyrophosphatase 1 (Ppa1), bear C-terminal N residues [log_2_(fold-change) > 0.30 and p-value < 0.05, **Figure 6a, Supplementary Table 6**]. Upon quantification of tryptic peptides that bear a C-terminal N or Q and are derived from a protein C-terminus, Glul[358–373] was the only peptide significantly upregulated upon Crbn loss (**EDF 9a**). Upregulation of Glul and Ppa1, but not Pcmt1, in Crbn^−/–^ hippocampi was validated by Western blot (**Figure 6b, EDF 9b, Supplementary Table 7**). The mRNA levels of Glul, Ppa1, and Pcmt1 were not significantly changed between WT and Crbn^−/–^ samples (**EDF 9c**), confirming that the accumulation of Glul and Ppa1 is a post-transcriptional effect. Cerebody enrichment revealed elevated levels of Glul-cN and cN-modified Ppa1 (Ppa1-cN) in the hippocampi of Crbn^−/–^ mice relative to WT hippocampi (**Figure 6c**), validating that the regulation of Glul and Ppa1 by Crbn occurs via the C-terminal cyclic imide in the brain. Collectively, these data point to the neurobiological connection of GLUL and a novel substrate PPA1 with PCMT1 and CRBN in vivo.

**Figure 6.**
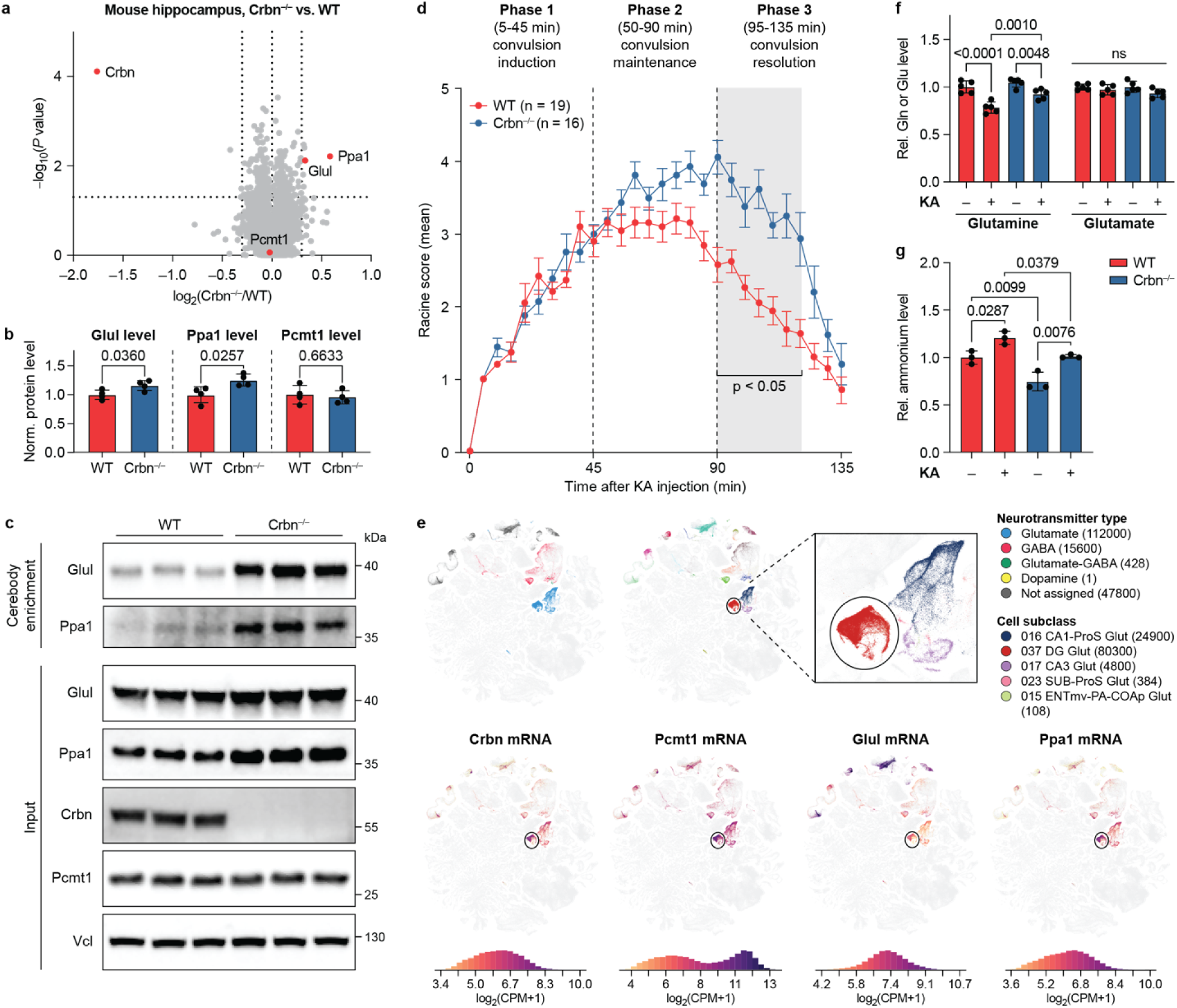
Loss of CRBN function affects the brain. **(a)** Quantitative proteomics of the hippocampus from 4 individual WT or Crbn^−/–^ mice. P-values for the abundance ratios were calculated by one-way ANOVA with TukeyHSD post-hoc tests. **(b)** Quantification of Glul, Ppa1, and Pcmt1 levels in WT or Crbn^−/–^ mouse hippocampus by Western blot across 4 biological replicates. Comparisons were performed using unpaired two-tailed t-tests, and p-values are indicated. **(c)** Cerebody enrichment for Glul-cN and Ppa1-cN from WT or Crbn^−/–^ mouse hippocampi across 3 biological replicates. **(d)** Time-course change in Racine scores of WT or Crbn^−/–^ mice after kainic acid (KA) injection. The shaded area indicates all the time points at which p < 0.05 by two-way ANOVA with Sidák’s multiple comparisons test. **(e)** Map of Crbn, Pcmt1, Glul, and Ppa1 expression in mouse hippocampus. The upper graphs are color coded according to the neurotransmitter type or cell subclass, while the lower graphs are color coded according to the normalized mRNA level of each gene. RNA sequencing data were extracted from the Brain Knowledge Platform. **(f)** Quantification of glutamine and glutamate levels from vehicle- or KA-treated (2 h-treatment) WT or Crbn^−/–^ mouse hippocampus by Glutamine/Glutamate-Glo assay. Comparisons were performed using two-way ANOVA with Šidák’s multiple comparisons tests, and p-values are indicated. **(g)** Quantification of ammonium levels from vehicle- or KA-treated (2 h-treatment) WT or Crbn^−/–^ mouse hippocampus. Comparisons were performed using one-way ANOVA with Šidák’s multiple comparisons tests, and p-values are indicated.

We confirmed the regulation of PPA1 by PCMT1 and CRBN in vitro and in cells. PPA1 is an enzyme that catalyzes the hydrolysis of pyrophosphate to inorganic phosphate^44^ and has a C-terminal N that is conserved back through fish (**EDF 9d**). The C-terminal 5-mer peptides from human and mouse PPA1 are methylated and cyclized by PCMT1 under physiological conditions (20 mM Tris, pH 7.4 at 37 °C), as monitored by mass spectrometry (**EDF 10a–b**), and these peptides bind CRBN only after incubation with PCMT1 (**EDF 10c**). Furthermore, accumulation of PPA1-cN was observed in HEK293T, SK-N-SH, and Neuro-2a cells upon loss or inhibition of CRBN (**EDF 10d**). These data establish that, like GLUL, PPA1 is regulated by CRBN via the C-terminal cyclic imide in vitro, in cells, and in vivo.

Epileptic seizure is one of the most prevalent neurological disorders and is proposed to stem from aberrant neuronal activity and the imbalance between the neurotransmitters glutamate and GABA.^45^ As both GLUL and PPA1 play a critical role in metabolism and the brain,^46–48^ perturbation of the CRBN-mediated regulation of these proteins may impact neuronal activity and hence the response to seizures. We treated WT and Crbn^−/–^ mice with kainic acid (KA), a glutamate homolog and neurotoxin that induces seizures,^49^ and evaluated the severity of seizures based on a modified Racine scale.^50^ Upon KA injection (20 mg/kg body weight, i.p.), the WT and Crbn^−/–^ mice initially showed similar latency to convulsions in Phase 1 of the experiment, but the Crbn^−/–^ mice maintained a higher Racine score indicative of more severe seizures in Phase 2 and exhibited a remarkably slower resolution of the convulsions in Phase 3 (**Figure 6d, EDF 9e–f**). These data show that loss of Crbn, like loss of Pcmt1,^18^ gives rise to a proepileptic phenotype in mice. Intriguingly, single cell RNA sequencing data in the mouse hippocampus retrieved from the Brain Knowledge Platform^51^ shows high co-localization and enrichment of Crbn, Pcmt1, Glul, and Ppa1 in the glutamatergic neurons (Glut) of the dentate gyrus (DG), strengthening the in vivo association between these proteins (**Figure 6e**).

We interrogated the molecular connection between the proepileptic phenotype and the elevated Glul levels in Crbn^−/–^ mice by monitoring levels of enzyme substrates (glutamate and ammonium) and product (glutamine) in the hippocampus. No change in glutamine or glutamate was observed between WT and Crbn^−/–^ mice at steady-state by a luminescent reporter assay (**Figure 6f**), which may be due to compensatory mechanisms. By contrast, glutamine levels decreased on proepileptic KA challenge, which was attenuated in the Crbn^−/–^ mice, an effect consistent with the elevated Glul levels measured in the hippocampus (**Figure 6f**). In addition, basal ammonium levels were reduced in Crbn^−/–^ mice relative to WT, and these levels increased on KA injection in both genotypes (**Figure 6g**). Taken together, these metabolic changes are consistent with elevated Glul levels on loss of Crbn that may influence response during recovery from KA-induced seizures, thereby illuminating a previously unknown connection between the protein damage–repair enzyme PCMT1, the E3 ligase CRBN, the regulation of the metabolic enzymes GLUL and PPA1, and effects on metabolic neuroadaptations during seizure response in the brain.

## Discussion

We discovered that PCMT1 enzymatically installs the C-terminal cyclic imide degron on proteins with C-terminal N, enabling their recognition and degradation by CRBN. This newly identified enzymatic mechanism for generating the C-terminal cyclic imide degron is distinct from the previously annotated roles of both PCMT1 and CRBN in mitigating non-enzymatic protein damage events, highlighting a previously overlooked function of these proteins in maintaining homeostasis of proteins like GLUL and PPA1. Our data point to the specific activity of this pathway on GLUL and PPA1 in the mouse hippocampus, likely on excitatory neurons, although additional substrates may be identified on further examination. These data indicate the likely existence of additional regulatory factors that cause substrates to enter this pathway, which may be dependent on the substrate or activity of PCMT1 and CRBN in specific contexts or cell types, analogous to the regulation of other protein modifications. Furthermore, both GLUL and PPA1 have been observed to respond to small molecule CRBN ligands in our hands and in other studies,^36,52^ indicating that endogenous substrates may be impacted by CRBN ligands in the clinic. Thus, future efforts will focus on illuminating substrates that are regulated by PCMT1, CRBN, and other contributing factors in different cell types and settings, including those that are more relevant to CRBN ligands in the clinic, and their potential contribution to desirable or undesirable clinical outcomes.

Interestingly, the substrates that we have thus far demonstrated to be regulated by PCMT1 and CRBN through the C-terminal cyclic imide are all highly conserved metabolic enzymes that form homo-oligomers and to our knowledge have not been previously associated with each other. GLUL is present in virtually all species from prokaryotes to eukaryotes and is expressed as a homodecamer in humans.^53^ PPA1 is also a highly conserved homodimeric protein involved in phosphate metabolism, and its loss is embryonically lethal.^54,55^ In parallel efforts to discover novel substrates, we have further annotated uroporphyrinogen decarboxylase (UROD) as a homodimeric substrate of both PCMT1 and CRBN.^36^ Further studies on additional substrates of PCMT1 and CRBN across species will reveal whether these features identified thus far from human and mouse models are important to the evolution and physiological function of this pathway.

Finally, this study of CRBN substrates that are regulated by PCMT1 will shed light on substrates regulated by CRBN that either do not have a C-terminal N or are not recognized through the thalidomide-binding domain. These proteins may be recruited to CRBN via a C-terminal cyclic imide-modified substrate, such as through a gain-of-function protein–protein interaction or through allosteric promotion of a conformational change in CRBN,^37^ comprise a downstream response to degradation of the substrate, or participate in unrelated regulatory pathways mediated by other enzymes. A unique feature of the C-terminal cyclic imide degron is the temporal nature of the modification’s formation. The time required for the PCMT1-mediated formation of the C-terminal cyclic imide ranges from 12 to 24 h under physiological conditions, suggesting a potential role of this protein degradation pathway as a molecular timer for processes of similar duration, such as the cell cycle, development, differentiation, or circadian rhythm. These insights into the enzymatic formation of the C-terminal cyclic imide degron on endogenous substrates and the associated biological consequences thus shed light on the role of PCMT1 and CRBN in signaling and metabolism and reveal the potential of CRBN ligands to influence these functions during therapy.

## Extended Data and Supplementary Information

are available for this paper.

## Acknowledgments

We thank S. Ichikawa, H. A. Flaxman, N. C. Payne, M. A. Leon-Duque, and T. Long for helpful discussions, T. Wang for support with peptide synthesis, M. Chen and S. Trager from the Harvard University Mass Spectrometry and Proteomics Resource Laboratory for support with MS experiments, D. Cui and B. Tresco from the Harvard University Laukien-Purcell Instrumentation Center for support with peptide MS experiments, and J. A. Nelson and Z. T. Niziolek from the Harvard University Bauer Core for support with flow cytometry. His_6_-CRBN/DDB1 was a generous gift from Boehringer Ingelheim. Support from the National Institutes of Health (R01GM141406, C.M.W.), the Blavatnik Biomedical Accelerator at Harvard University (C.M.W.), Mark Foundation for Cancer Research (C.M.W.), Merkin Family Foundation (C.M.W.), the Starr Foundation (C.M.W.), and Spanish Ministerio de Ciencia, Innovación y Universidades (MICINU/FEDER; PID2021-125118OB-I00, M.G.) is gratefully acknowledged. N.Z. is supported by Howard Huges Medical Institute.

## Author Contributions

Z.Z., W.X., A.K.D.P. designed and synthesized peptides. E.Y.F., S.C., W.X., H.C.L., A.K.D.P. generated recombinant and engineered proteins. Z.Z., W.X., E.Y.F. performed in vitro enzyme activity and TR-FRET binding studies. S.C., N.Z. designed and conducted structural, BLI, and in vitro ubiquitination studies. Z.Z., W.X., E.Y.F. performed cellular studies. H.C.L. developed cerebody methods. A.H.L., P.P.V., M.G. designed and conducted in vivo measurements. E.Y.F., W.X. analyzed mouse tissue samples. C.M.W. conceived of the project. C.M.W., Z.Z., W.X., E.Y.F., S.C. drafted the manuscript. All authors reviewed and edited the manuscript.

## Data Availability

All data are available in the main text, Supplementary Information, and Supplementary Tables. Proteomics data have been deposited to the PRIDE repository with the dataset identifier PXD061730. Crystal structure of human CRBN/DDB1 in complex with GLUL-cN has been deposited to the Protein Data Bank (PDB) with the accession number 9NR3.

## Code Availability

All code used in this study is commercially available or provided in a public repository. A full description of software used is provided in the Supplementary Information.

---

**Extended Data Figure 1.**
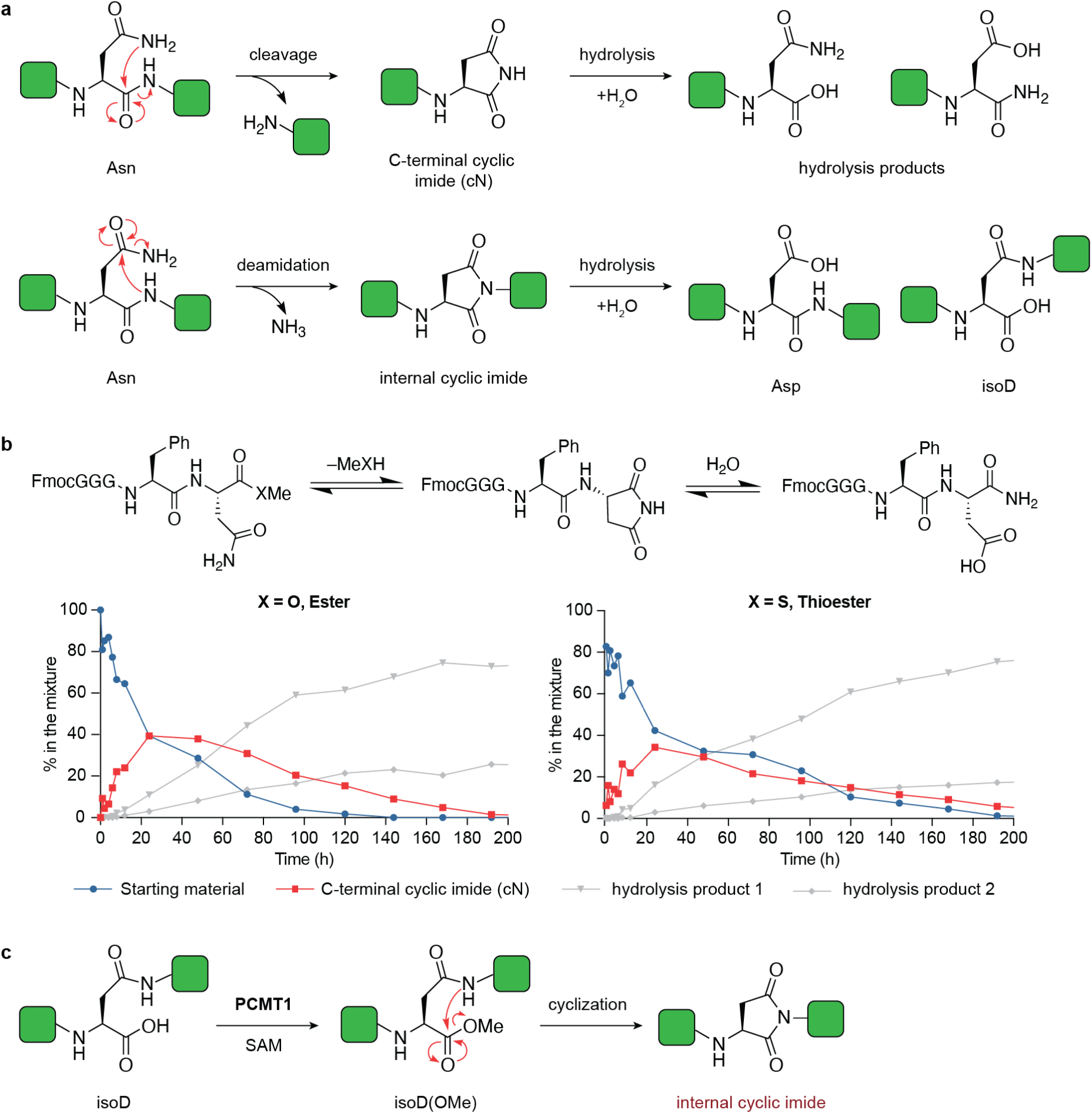
Formation of cyclic imides from asparagine. **(a)** Schematic of protein damage events that occur through Asn residues that result in the C-terminal cyclic imide or isoaspartate post-translational modifications. **(b)** Chemical scheme and conversion of model peptides incubated in 20 mM NH_4_OAc, pH 7.4 buffer at 37 °C. The experiment was performed in triplicate. **(c)** Schematic of PCMT1 correction of isoaspartate modifications.

**Extended Data Figure 2.**
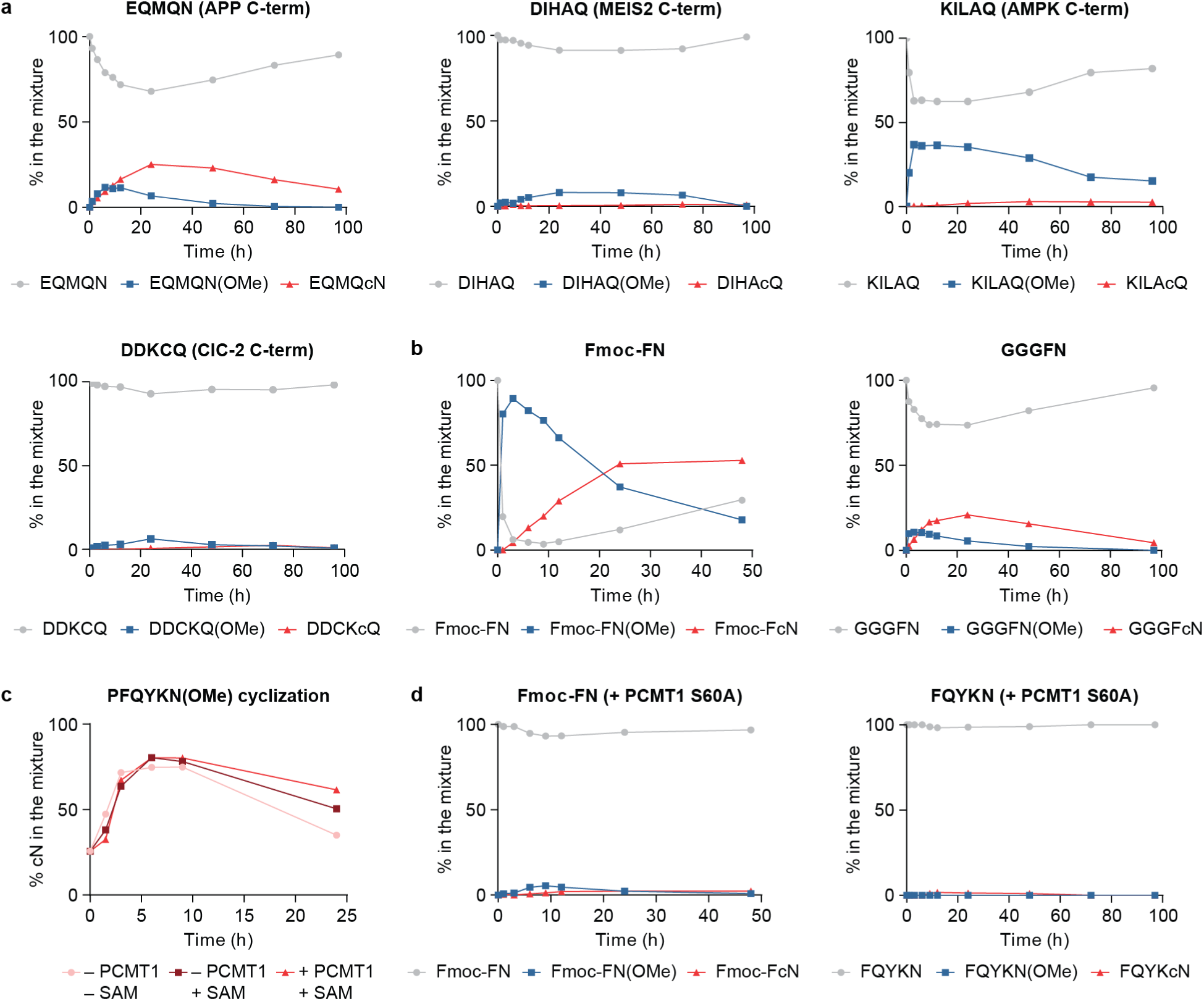
Efficiency of PCMT1-mediated C-terminal cyclic imide formation on various peptides. **(a)** Time-course incubation of peptides representing the C-termini of CRBN substrates with PCMT1 and SAM in 50 mM Tris-Cl, pH 7.4 buffer at 37 °C. **(b)** Time-course incubation of Fmoc-FN and GGGFN with PCMT1 and SAM in 50 mM Tris-Cl, pH 7.4 buffer at 37 °C. **(c)** Time-course incubation of fully methylated peptide PFQYKN(OMe) in 50 mM Tris-Cl, pH 7.4 buffer at 37 °C with or without PCMT1 and SAM. **(d)** Time-course incubation of Fmoc-FN and FQYKN with the catalytically inactive PCMT1 and SAM in 50 mM Tris-Cl, pH 7.4 buffer at 37 °C.

**Extended Data Figure 3.**
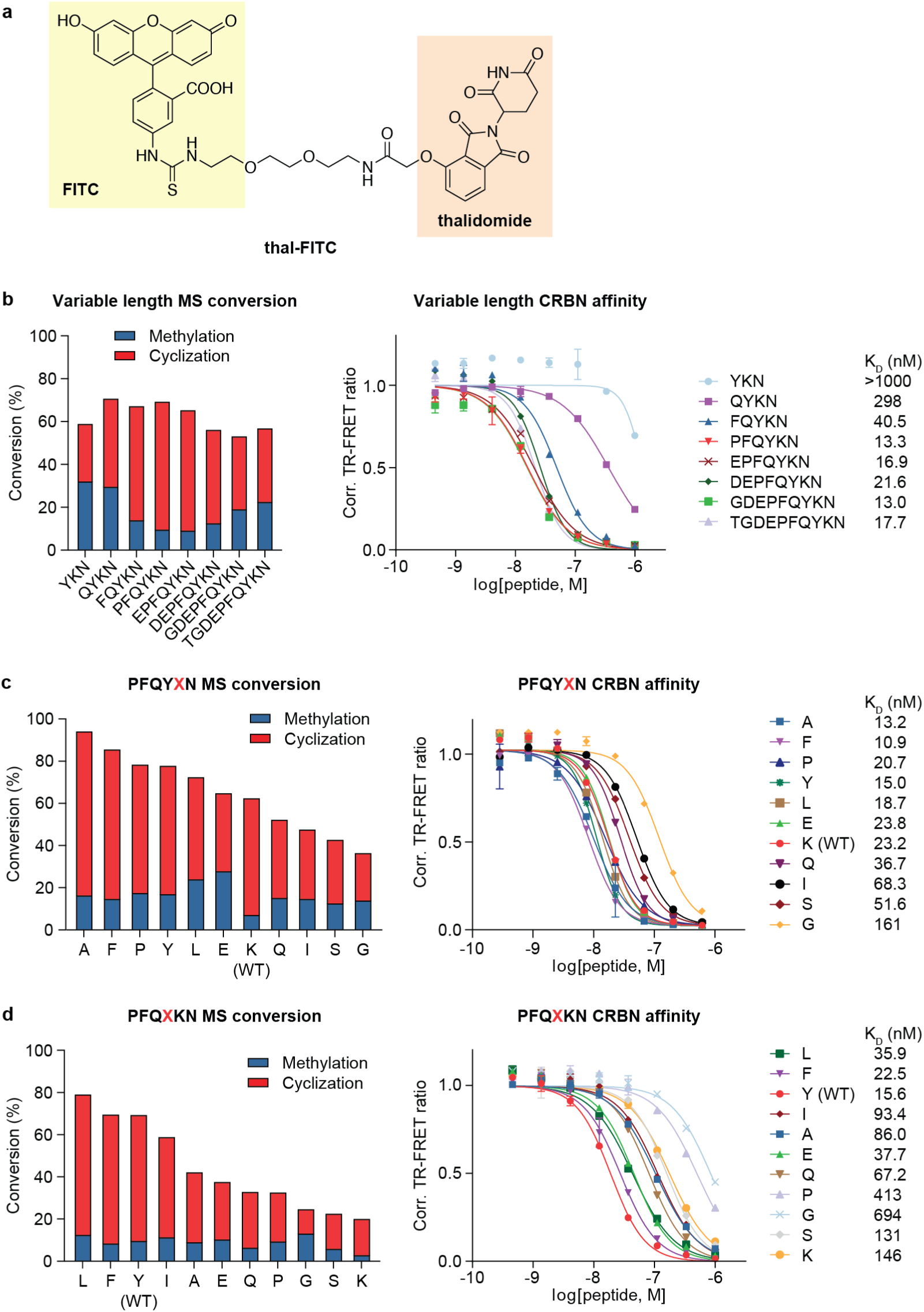
Sequence preference of PCMT1 activity and CRBN recognition. **(a)** Structure of thalidomide-FITC, the TR-FRET tracer used to assess the engagement of ligands with His_6_-CRBN/DDB1 in this study. **(b)** Conversion efficiency and CRBN binding of GLUL peptides with variable lengths after 14 h incubation with PCMT1 by mass spectrometry and TR-FRET assay, respectively. **(c)** Conversion efficiency and CRBN binding of GLUL peptides with variable –1 residue after 14 h incubation with PCMT1 by mass spectrometry and TR-FRET assay, respectively. **(d)** Conversion efficiency and CRBN binding of GLUL peptides with variable –2 residue after 14 h incubation with PCMT1 by mass spectrometry and TR-FRET assay, respectively. All TR-FRET experiments were performed with 3 technical replicates, and data are presented as mean ± s.d. (n = 3).

**Extended Data Figure 4.**
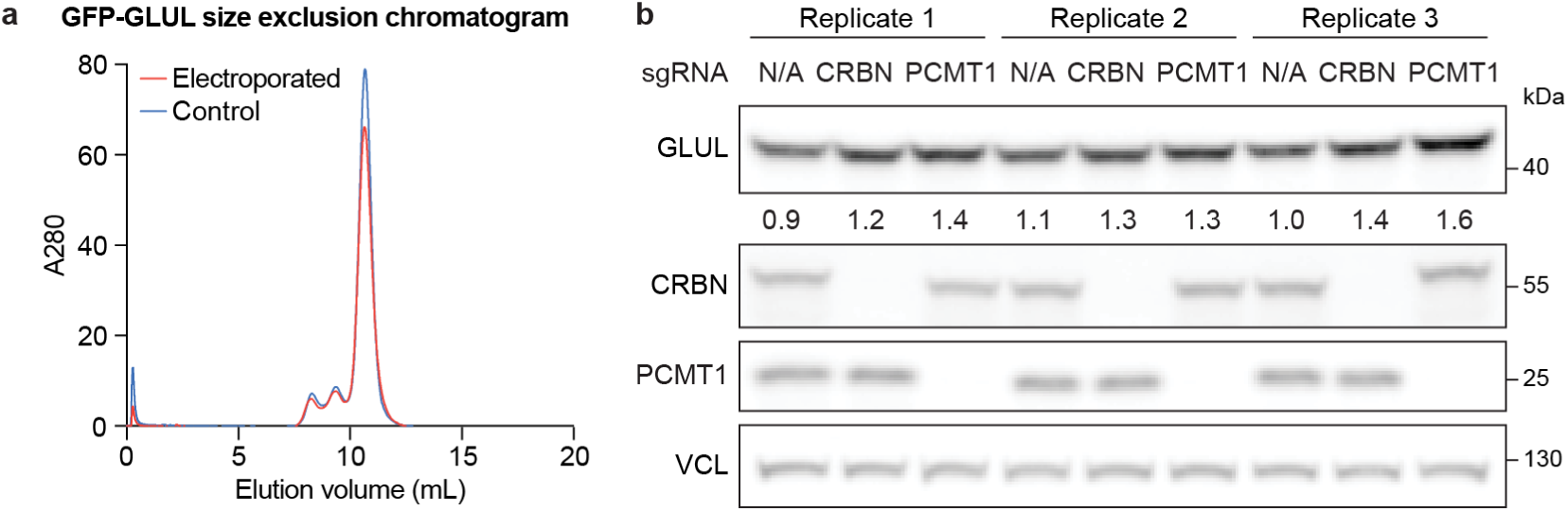
Full-length GLUL is recognized and regulated by PCMT1 and CRBN. **(a)** Size exclusion chromatography of the GFP-GLUL protein before or after electroporation. The oligomerization state of the protein did not change upon electroporation. **(b)** Western blot of endogenous GLUL levels in WT, PCMT1-KO, or CRBN-KO HEK293T cells across 3 biological replicates.

**Extended Data Figure 5.**
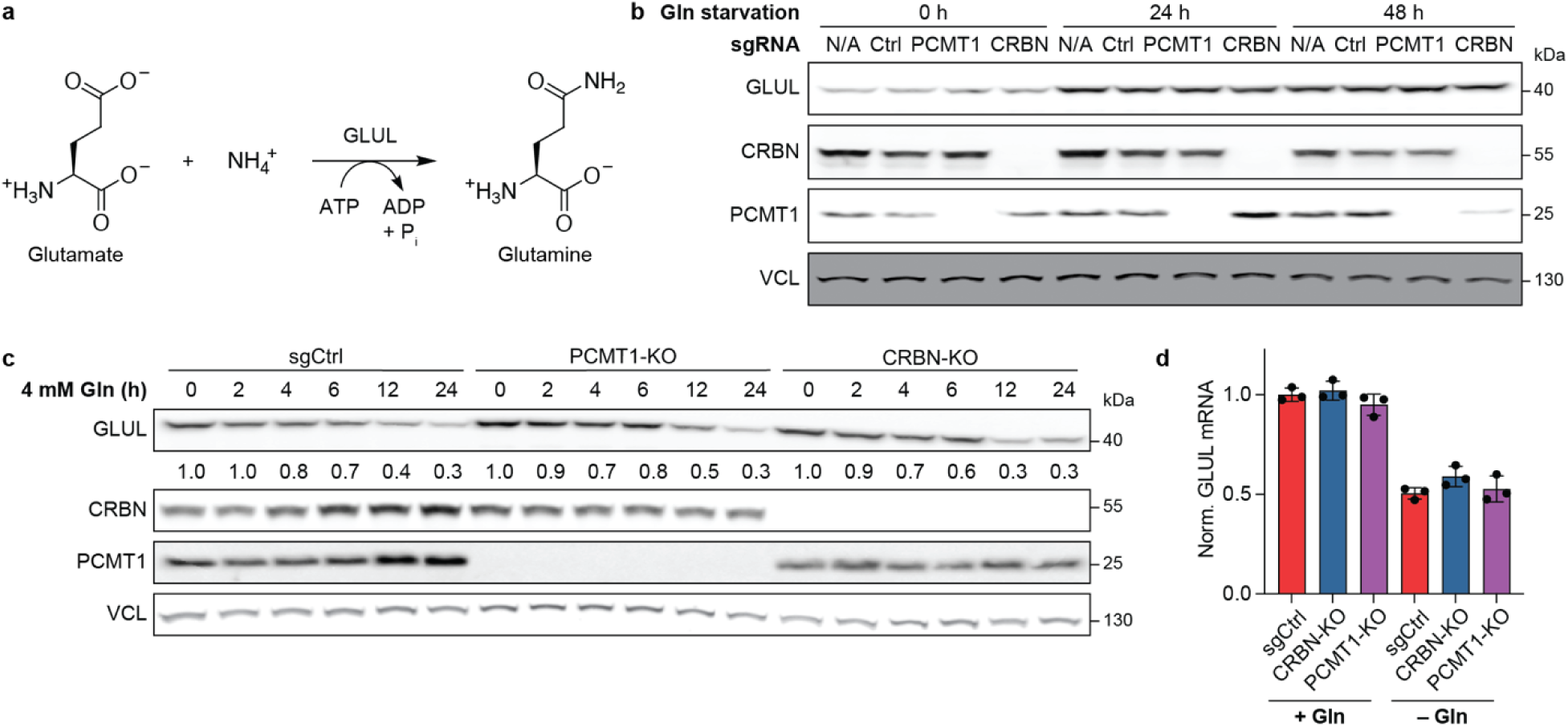
Glutamine-triggered GLUL degradation is independent of PCMT1. **(a)** Scheme of GLUL-catalyzed condensation of glutamate and ammonium into glutamine. **(b)** Time-course GLUL buildup in sgCtrl, PCMT1-KO, or CRBN-KO HEK293T cells incubated in glutamine-free media. **(c)** Time-course GLUL degradation of glutamine-starved sgCtrl, PCMT1-KO, or CRBN-KO HEK293T cells after refeeding with 4 mM glutamine. **(d)** Normalized mRNA level of GLUL in HEK293T cells with or without glutamine starvation for 48 h. The experiment was performed with 3 biological replicates.

**Extended Data Figure 6.**
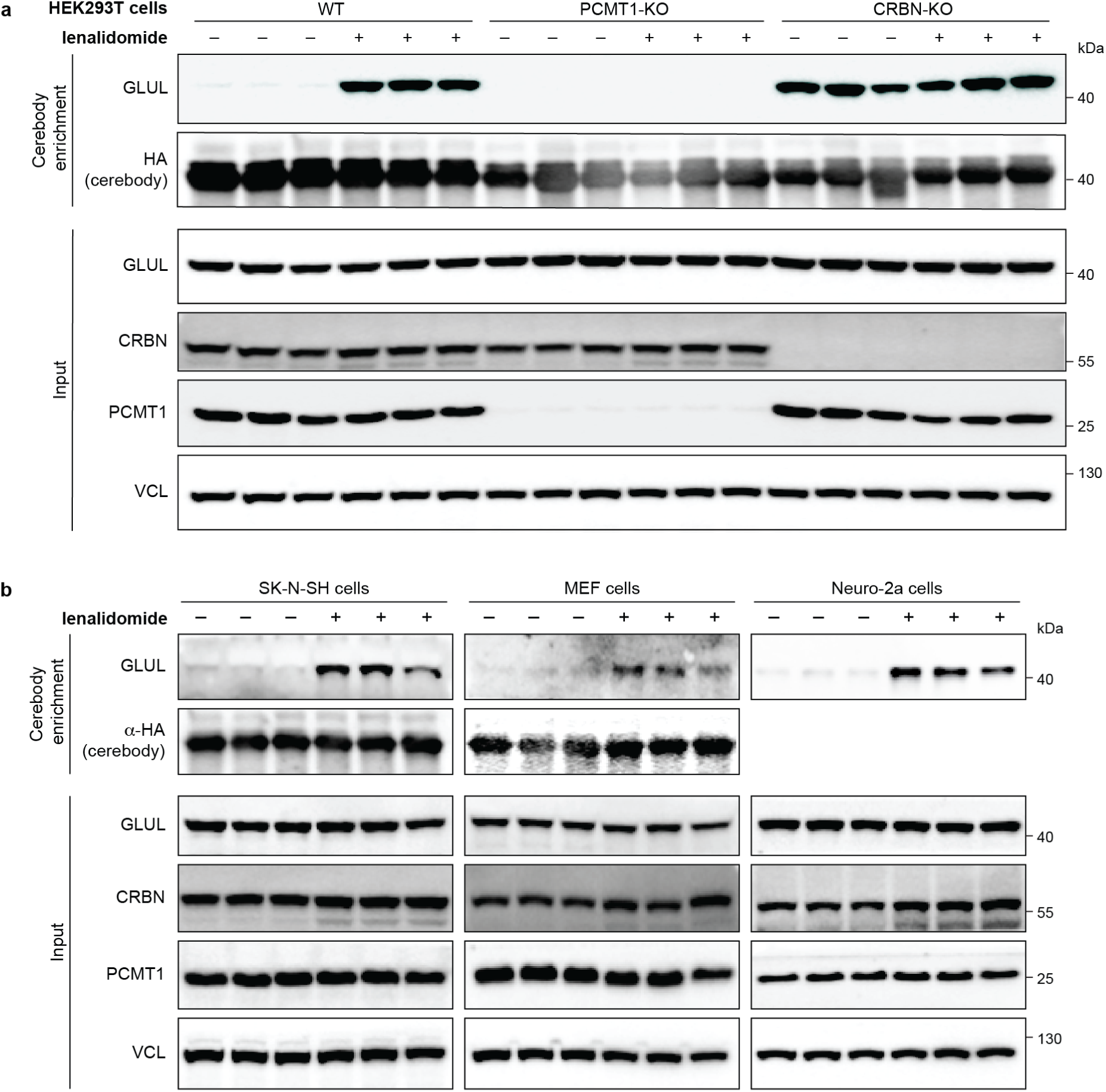
Impact of clinical agents on the GLUL-cN level in cells. **(a)** Cerebody enrichment for GLUL-cN from WT, PCMT1-KO, or CRBN-KO HEK293T cells after treatment with DMSO or 100 µM lenalidomide for 24 h across 3 biological replicates. **(b)** Cerebody enrichment for GLUL-cN from different human and mouse cell lines after treatment with DMSO or 100 µM lenalidomide for 24 h across 3 biological replicates. Elution for SK-N-SH and MEF was performed with 3× FLAG peptide, while elution for Neuro-2a was performed with 200 µM lenalidomide.

**Extended Data Figure 7.**
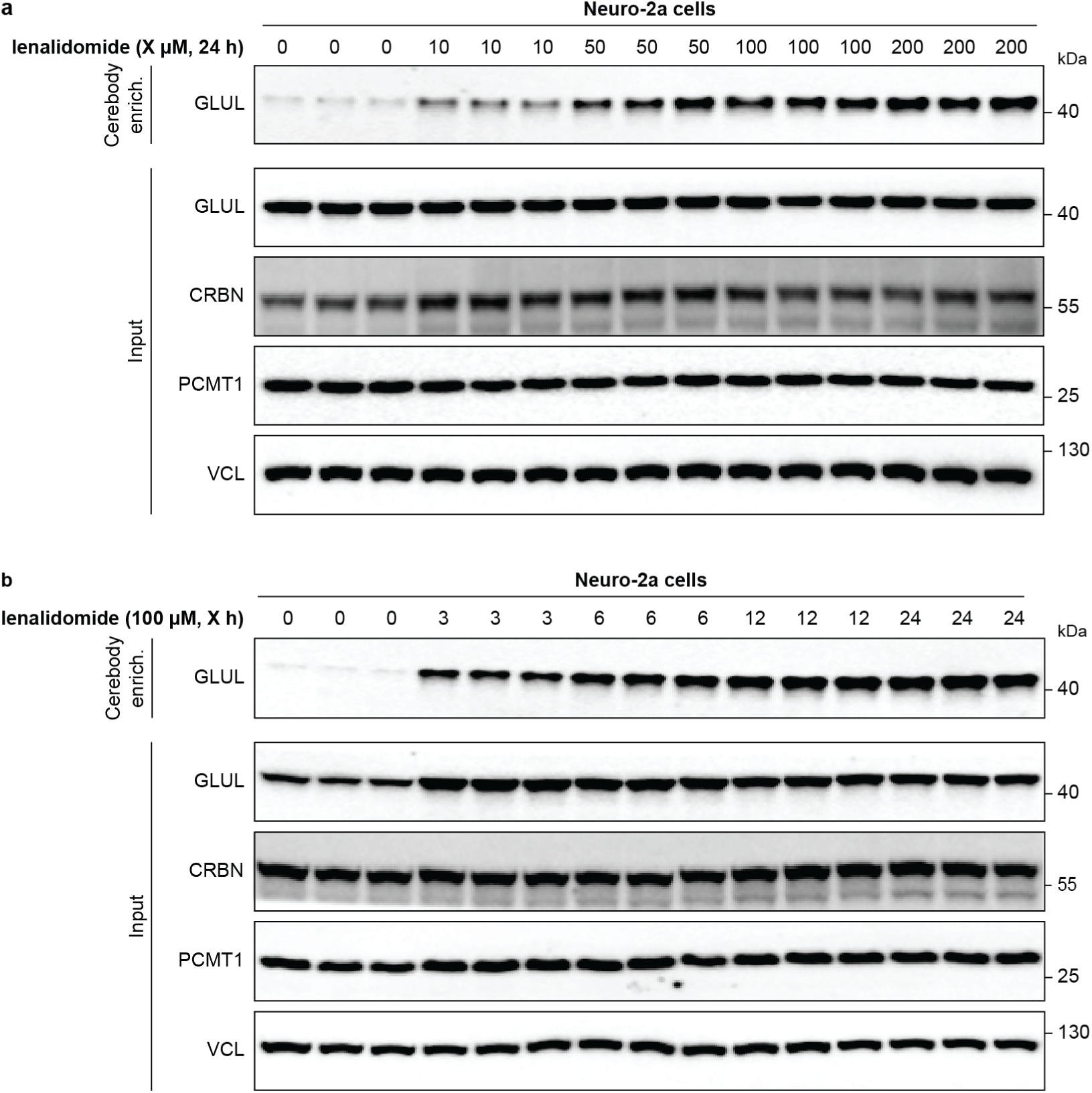
Time- and dose-dependence of clinical agents’ impact on GLUL-cN in cells. **(a)** Cerebody enrichment for GLUL-cN from Neuro-2a cells after treatment with varying doses of lenalidomide for 24 h across 3 biological replicates. **(b)** Cerebody enrichment for GLUL-cN from Neuro-2a cells after treatment with 100 µM lenalidomide for varying durations across 3 biological replicates.

**Extended Data Figure 8.**
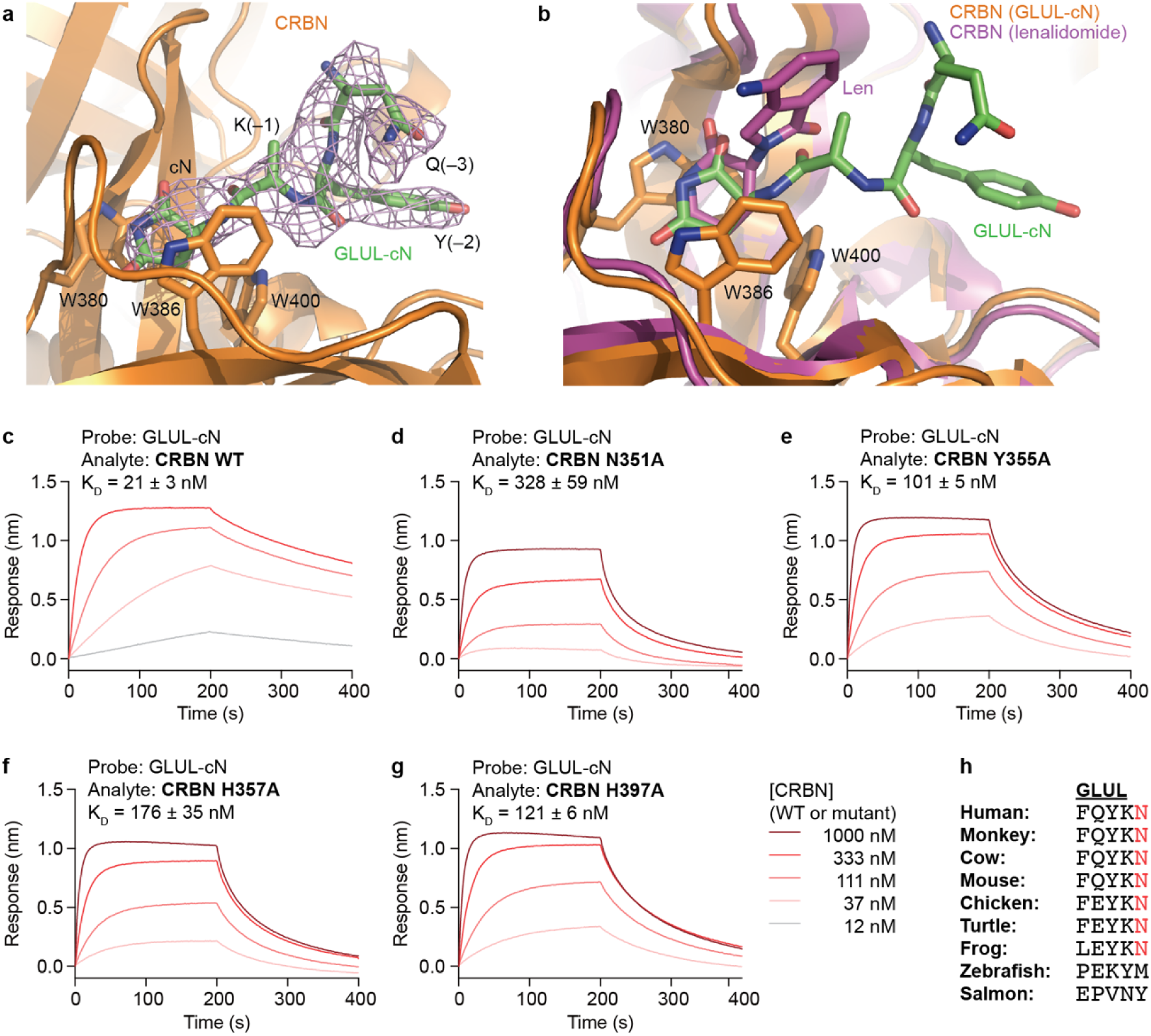
Characterization of GLUL-cN binding to CRBN. **(a)** Close-up view of GLUL-cN in complex with CRBN. GLUL-cN is shown as a stick model, together with its positive F_o_ – F_c_ electron density (purple mesh) calculated and contoured at 2.5σ before it was built into the CRBN-DDB1 model. The electron density for the side chain of lysine at the –1 position is incomplete (likely due to flexibility) and is thus not included in the model. CRBN is show as an orange cartoon with the tri-Trp side chains highlighted. **(b)** CRBN-GLUL-cN aligned with CRBN-lenalidomide (PDB: 5FQD). For CRBN-GLUL-cN, CRBN is shown as an orange cartoon with the tri-Trp side chains highlighted. GLUL-cN is shown as sticks (carbon: green, oxygen: red, nitrogen: blue). For CRBN-lenalidomide, CRBN is shown as a purple cartoon, and lenalidomide (Len) is show as sticks (carbon: purple, oxygen: red, nitrogen: blue). **(c–g)** BLI measurements of GLUL-cN binding to CRBN WT or mutants. **(h)** The C-terminal residues of GLUL across species from different classes of vertebrates. The conserved C-terminal N residues are highlighted in red.

**Extended Data Table 1.** X-ray crystallography data collection and refinement statistics.

| DDB1-CRBN-GLUL-cN |  |
| --- | --- |
| **Data collection** |  |
| Space group | P 2<sub>1</sub> 2<sub>1</sub> 2<sub>1</sub> |
| Cell dimensions |  |
| *a*, *b*, *c* (Å) | 76.435, 100.364, 169.672 |
| $\alpha$, $\beta$, $\gamma$ (°) | 90, 90, 90 |
| Resolution (Å) | 49.27 - 2.93 (2.96 - 2.93) * |
| *R*<sub>merge</sub> | 0.186 (2.055) |
| *I* / $\sigma$ *I* | 7.8 (0.9) |
| Completeness (%) | 99.73 (99.46) |
| Redundancy | 6.5 (6.3) |
| **Refinement** |  |
| Resolution (Å) | 49.27 - 2.93 |
| No. reflections | 28693 (923) |
| *R*<sub>work</sub> / *R*<sub>free</sub> | 0.2502/ 0.2780 |
| No. atoms | 9115 |
| Protein | 9080 |
| Ligand/ion | 35 |
| Water | 0 |
| *B*-factors | 94.54 |
| Protein | 94.56 |
| Ligand/ion | 88.99 |
| Water | N.A. |
| R.m.s. deviations |  |
| Bond lengths (Å) | 0.004 |
| Bond angles (°) | 0.611 |

Note: A single crystal was used for diffraction data collection and structure determination.
\*Values in parentheses are for the highest-resolution shell.

**Extended Data Figure 9.**
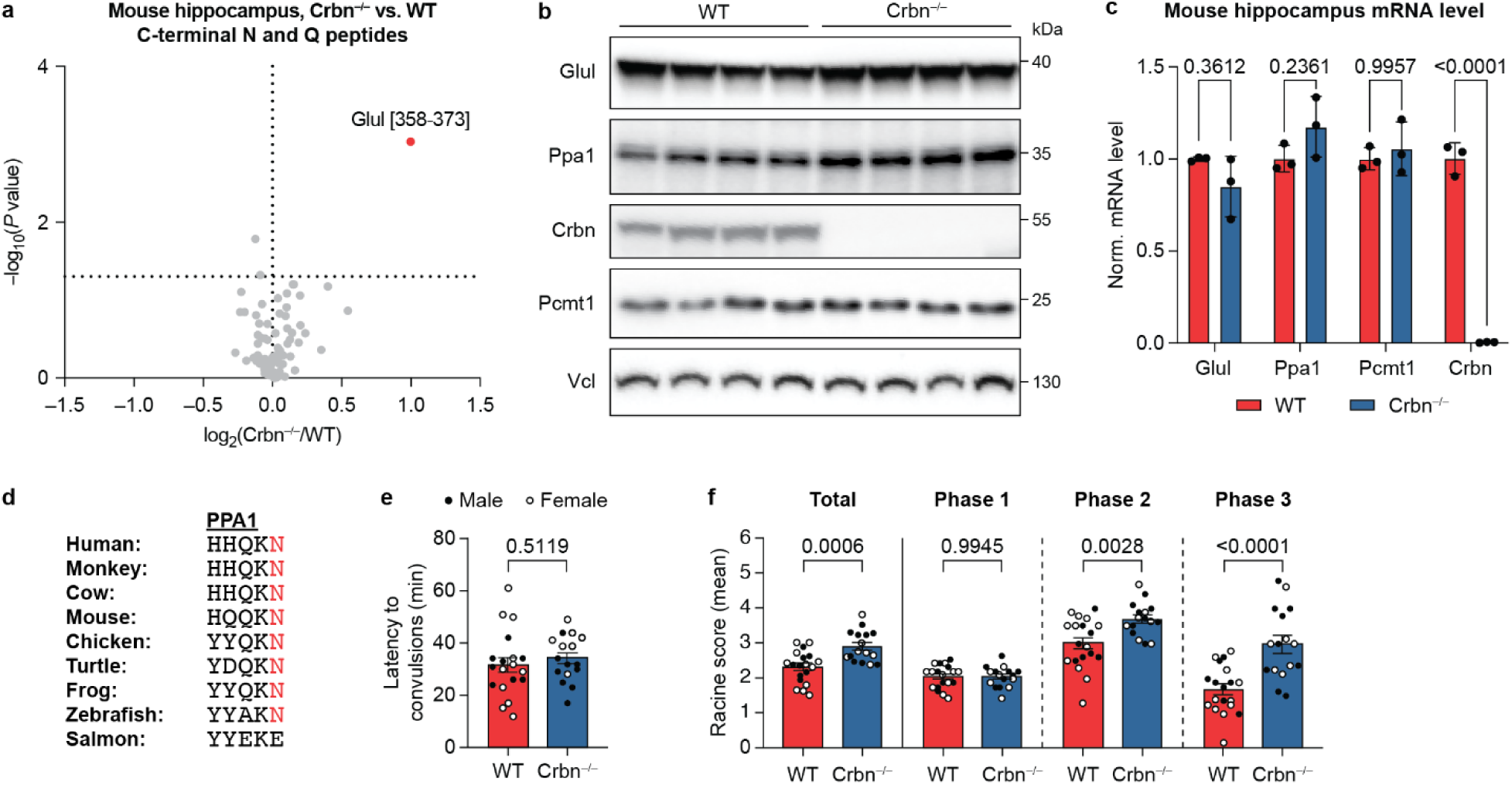
Genetic knockout of Crbn leads to accumulation of Glul and Ppa1 and neurological phenotypes in mouse models. **(a)** Volcano plot of tryptic peptides derived from protein C-termini that bear a C-terminal N or Q identified from the hippocampi of 4 individual WT or Crbn^−/–^ mice. P-values for the abundance ratios were calculated by one-way ANOVA with TukeyHSD post-hoc tests. **(b)** Western blot of Glul and Ppa1 levels in WT or Crbn^−/–^ mouse hippocampus. **(c)** Normalized mRNA levels of Glul, Ppa1, Pcmt1, and Crbn in WT or Crbn^−/–^ mouse hippocampus across 3 biological replicates. Comparisons were performed using unpaired two-tailed t-tests, and p-values are indicated. **(d)** The C-terminal residues of PPA1 across species from different classes of vertebrates. The conserved C-terminal N residues are highlighted in red. **(e)** Comparison of the time from KA injection to the first observation of convulsions (stage 3 of Racine scale) in WT and Crbn^−/–^ mice. **(f)** Comparison of Racine scores of WT and Crbn^−/–^ mice during each phase of the convulsion response. Comparisons were performed using unpaired two-tailed t-tests, and p-values are indicated.

**Extended Data Figure 10.**
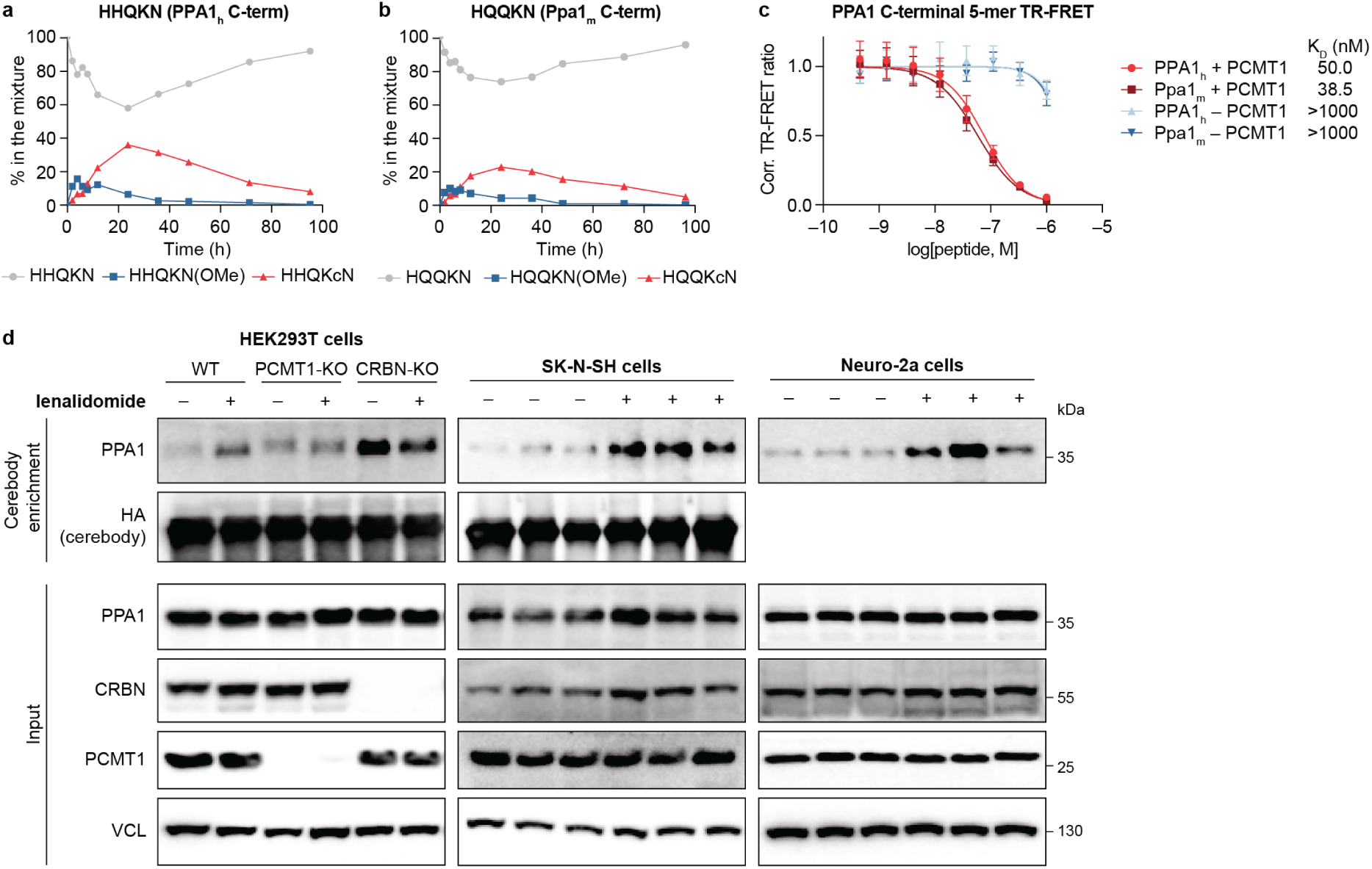
PPA1 is a substrate of PCMT1 and CRBN. **(a)** Time-course incubation of the peptide representing the C-terminus of human PPA1 with PCMT1 and SAM in 50 mM Tris-Cl, pH 7.4 buffer at 37 °C. **(b)** Time-course incubation of the peptide representing the C-terminus of mouse Ppa1 with PCMT1 and SAM in 50 mM Tris-Cl, pH 7.4 buffer at 37 °C. **(c)** TR-FRET assay of the human or mouse PPA1 C-terminal peptides incubated with or without PCMT1 to measure their affinity to His_6_-CRBN/DDB1. **(d)** Cerebody enrichment for PPA1-cN from different human and mouse cell lines after treatment with DMSO or 100 µM lenalidomide for 24 h across 3 biological replicates. Elution for HEK293T and SK-N-SH was performed with 3× FLAG peptide, while elution for Neuro-2a was performed with 200 µM lenalidomide.

