## Supplementary Information for "PCMT1 generates the C-terminal cyclic imide degron on CRBN substrates"

**This PDF file includes:**

Supplementary Figures 1 to 4
Methods
Materials and Instrumentation
Synthetic Procedures
NMR Spectra and LC-MS Traces
SI References

**Other Supplementary Materials for this manuscript include the following:**

Supplementary Information Tables 1 to 7 (Excel)

### I. Supplementary Figures

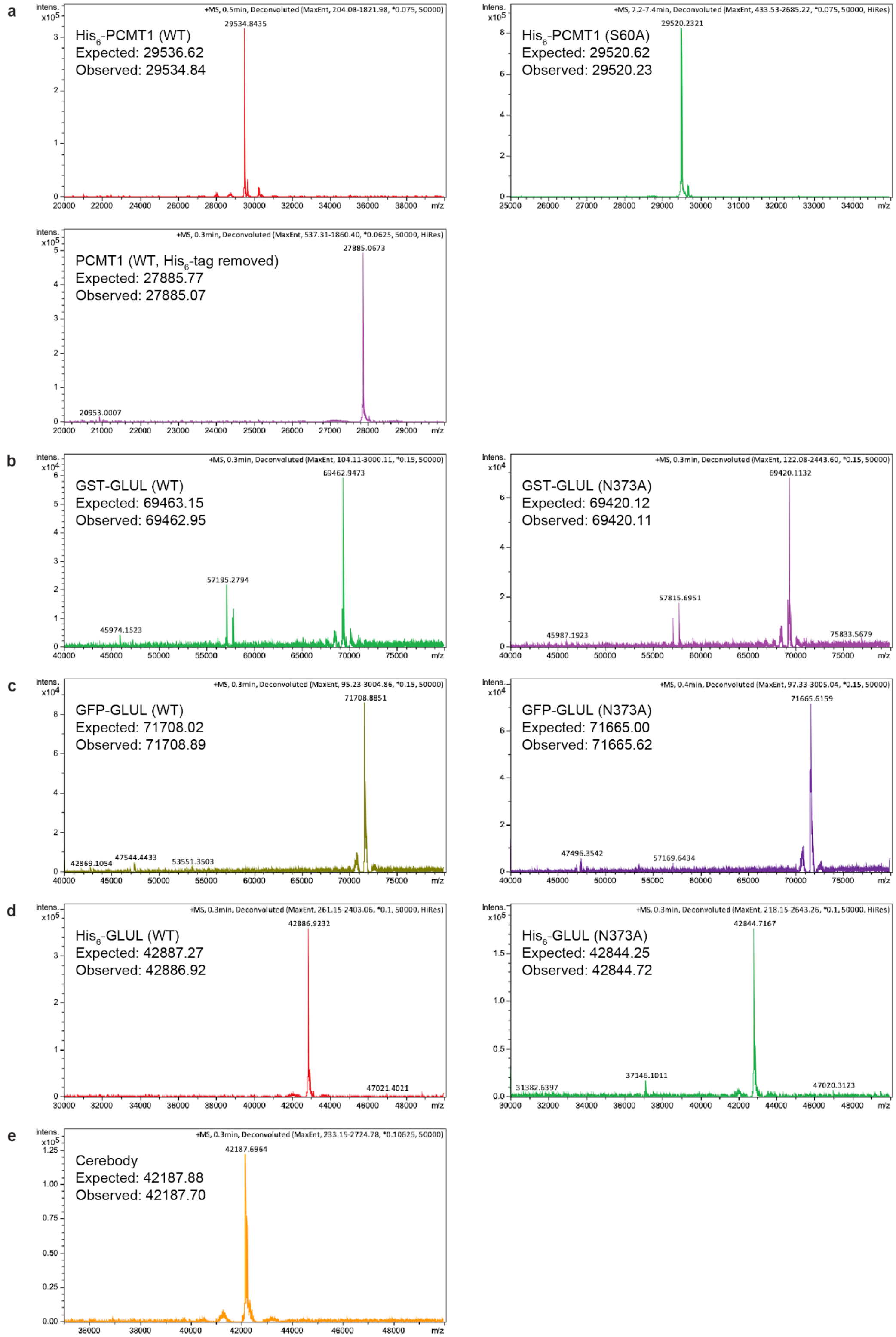

**Supplementary Figure 1. (a)** Intact MS spectra of PCMT1 constructs. **(b)** Intact MS spectra of GST-GLUL constructs. **(c)** Intact MS spectra of GFP-GLUL constructs. **(d)** Intact MS spectra of $His_6$-GLUL constructs. **(c)** Intact MS spectra of cerebody.

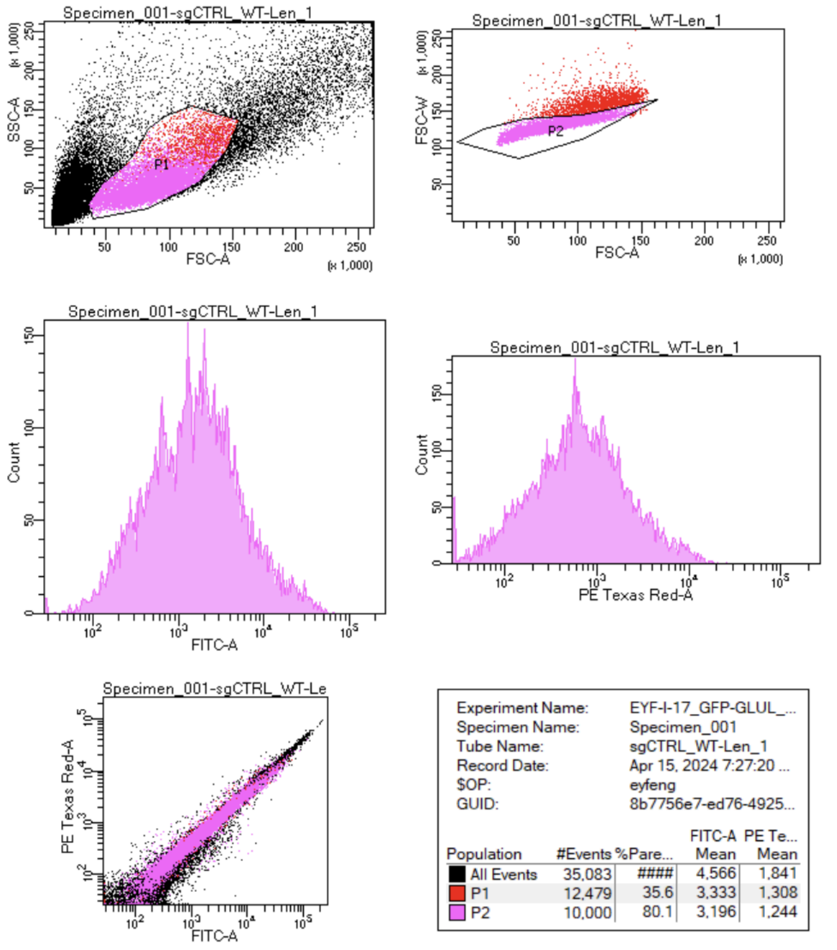


**Supplementary Figure 2.** Representative flow cytometry gating for GFP-GLUL electroporation experiments.

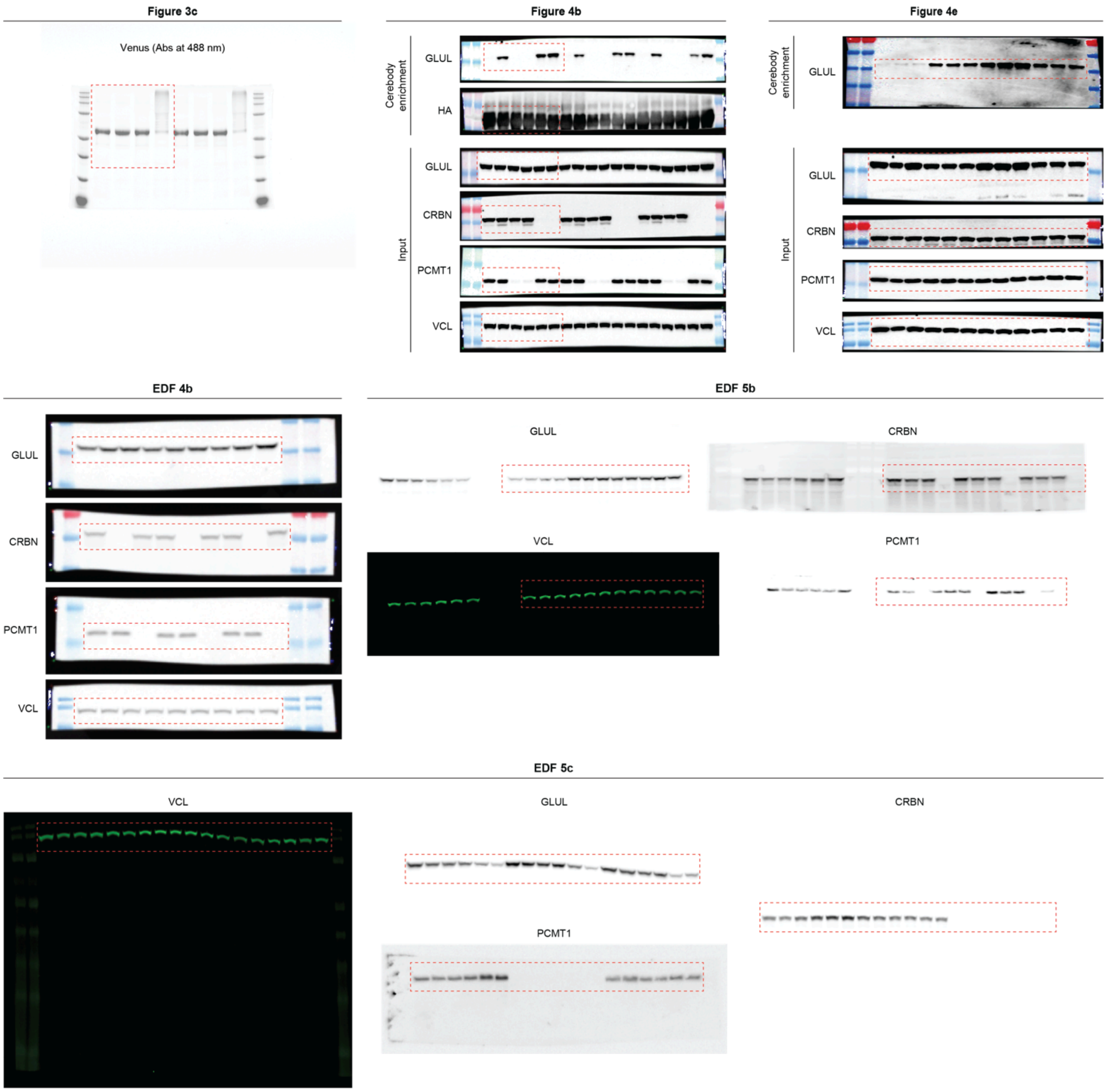


**Supplementary Figure 3.** Uncropped images for western blots shown in Figure 3, Figure 4, EDF 4, and EDF 5.

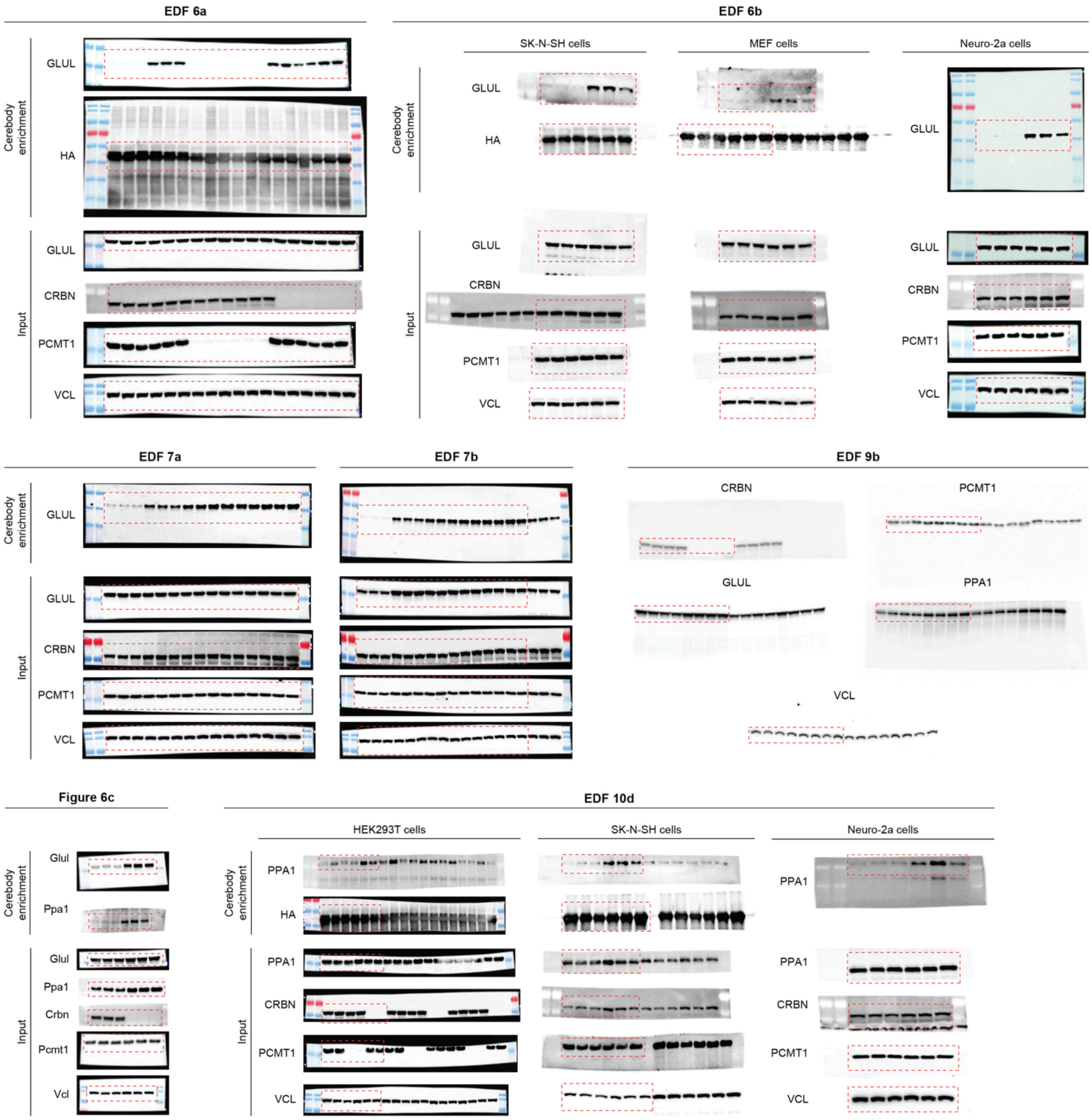


**Supplementary Figure 4.** Uncropped images for western blots shown in Figure 6, EDF 6, EDF 7, EDF 9, and EDF 10.

#### II. Methods

General cell culture protocol

All cell lines were cultured in DMEM supplemented with 10% heat-inactivated fetal bovine serum (FBS) and 1× penicillin-streptomycin (denoted DMEM+/+), except for MEF cell lines which were cultured in DMEM supplemented with 15% heat-inactivated FBS and 1× penicillin-streptomycin. Cells were grown at 37 °C in a humidified atmosphere with 5% $CO_2$ unless otherwise noted. Mycoplasma testing was performed regularly for all cell lines to check for contamination. For collection of cell pellets, cells were dissociated with 0.25% trypsin-EDTA for 3 min at 37 °C, then washed twice in PBS with centrifugation at 500 × g, 24 °C, 3 min. The pellets were flash-frozen with liquid nitrogen and stored at –80 °C until use.

General western blotting protocol

For analysis of cell samples, cells were lysed in 2% SDS/PBS with probe sonication (5 sec on, 3 sec off, 15 sec total, 10%), or in Pierce IP lysis buffer supplemented with 1× protease/phosphatase inhibitor cocktail. After clearing the lysates by centrifugation (21,000 × g, 4 °C, 10 min), the supernatant of each sample was transferred to a new tube, and the total protein concentration was measured by BCA assay. The lysates were diluted to 1–3 mg/mL, mixed with 5× SDS-PAGE loading buffer (1× concentration: 50 mM Tris–HCl, 2% SDS, 10% glycerol, 1% β-mercaptoethanol, 0.02% bromophenol blue) and boiled at 95 °C for 5 min. For analysis of recombinant proteins, the protein solution was directly mixed with SDS-PAGE loading buffer and boiled at 95 °C for 5 min. The denatured samples (8–10 µL) were loaded on a 12% Criterion TGX 26-well precast gel and run at 150 V for 60 min in Tris/Glycine/SDS buffer. The gel was transferred to a nitrocellulose membrane using an Invitrogen iBlot 2 dry blotting system with the preset P0 program. The membrane was cut, blocked with 5% BSA/TBST or 5% milk/TBST, and incubated with the primary antibody at a 1:1000 dilution at 4°C overnight. The membrane was washed 3× with TBST and incubated with the secondary antibody at a 1:10,000 dilution at 24 °C for 1 h, after which it was washed 3× with TBST again. The washed membrane was then imaged using the chemiluminescence or IR800 channel on an Azure imager.

Formation study of model peptides

Fmoc-GGGFN(OMe) and Fmoc-GGGFN(SMe) were dissolved and aliquoted in DMSO, then resuspended and incubated separately in 20 mM $NH_4OAc$ buffer, pH 7.4 at a final peptide concentration of 67 µM. Samples were incubated in a sand bath pre-heated to 37 °C. An aliquot was taken at t = 0 h (collected immediately after resuspension in buffer), then at subsequent desired time points. The samples were quenched by addition of formic acid to pH < 4 and immediately moved to –20 °C and stored until analysis. MS samples were prepared by mixing 7 µL of the incubation mixture with 43 µL acetonitrile and analyzed on a Waters ACQUITY UPLC system equipped with an SQ Detector 2 mass spectrometer. The extracted ion chromatograms of each species (with mass tolerance of ± 0.01) were background-subtracted and integrated on Mnova.

Protein preparation

Plasmids were constructed with HiFi DNA Assembly Kit (NEB), and mutations were generated using QuikChange Lightning kit (Agilent) or Q5 site-directed mutagenesis kit (NEB). All plasmids were validated by whole plasmid sequencing.

$His_6$-PCMT1, $His_6$-GLUL (WT or N373A), GST-GLUL (WT or N373A), $His_6$-GFP-GLUL (WT or N373A), and $His_6$-mCherry were expressed in *E. coli* BL21(DE3) cells, which were inoculated in an overnight starter culture in LB with 100 µg/mL carbenicillin or 50 µg/mL kanamycin. Large-scale overexpression cultures (1 L × 5) were inoculated with the saturated overnight culture diluted 1:100. The overexpression cultures were incubated at 37 °C with shaking until the OD600 reached 0.6–0.8, at which point the temperature was reduced to 16–20 °C and IPTG was added to a final concentration of approximately 0.4 mM. The bacteria were allowed to express the protein overnight, after which they were harvested by centrifugation. The cell pellet was resuspended in 40 mL lysis buffer (20 mM imidazole, 1× protease inhibitor, 1% Triton-X 100 in PBS), then sonicated (6 sec on, 60 sec off, 5 min total on time, 70% amplitude using a Fisherbrand model 120 sonicator) on ice. The lysate was then clarified by centrifugation (20,000 × g, 4 °C, 10 min) and syringe filtration (0.45 µm). $His_6$-tagged protein was purified by Ni-NTA affinity chromatography followed by size exclusion chromatography. GST-tagged protein was purified with glutathione sepharose 4B resin (Cytiva), using 0.5 mL resin per 1 L of expression culture. Protein-containing fractions were pooled and concentrated using Vivaspin 10 kDa MWCO spin concentrators (Cytiva). Glycerol was added to a final concentration of 10%, after which the protein was aliquoted, flash-frozen, and stored at –80 °C.

The $His_6$-tag on $His_6$-PCMT1 was cleaved with thrombin agarose resin (Sigma) following the manufacturer's protocol.

Cerebody was overexpressed and purified as previously described.[1]

$His_6$-tagged CRBNΔ1-40 and tag-free DDB1 (where BPB domain 394-708 is replaced by a glycine-linker) were co-expressed in Hi5 monolayer insect cells. $His_6$-tagged CRBNΔ1-40, DDB1 (full length), and DDA1 were co-expressed in Hi5 monolayer insect cells. The CRBN-DDB1 complex or the CRBN-DDB1-DDA1 complex was isolated from cell lysate by Ni-NTA affinity chromatography. After cleavage by TEV protease, tag-free CRBN-DDB1 or CRBN-DDB1-DDA1 complex was further purified by anion exchange and size exclusion chromatography. GST-tagged or $His_6$-Venus-tagged GLUL (full-length or C-terminal 6-mer peptide) was expressed alone or co-expressed with PCMT1 in *E. coli* BL21(DE3) cells. For PCMT1 co-expression, the LB broth was supplemented with 6 µM SAM. $His_6$-tagged CRBN[2] (WT or mutant) was expressed in *E. coli* BL21(DE3) cells in LB broth was supplemented with 50 µM $ZnCl_2$. $His_6$-CRBN was purified by Ni-NTA affinity chromatography followed by size exclusion chromatography.

In vitro PCMT1 incubation of peptides and MS analysis

Unless otherwise noted, the peptide (10 µM) was incubated with $His_6$-PCMT1 (10 µM, WT or S60A) and SAM (200 µM) in 50 mM Tris-Cl buffer, pH 7.4 at 37 °C. Samples were collected after 14 h for an end-point analysis, or at the desired time points for a time-course study. The samples were quenched by addition of formic acid or trifluoroacetic acid to pH < 4 and stored at –80 °C until use. The MS samples were prepared by mixing 2.5 µL of the incubation mixture and 20 µL acetonitrile and analyzed on a Bruker micrOTOF II LC-MS system. The extracted ion chromatograms of the acyclic, methyl ester, and C-terminal cyclic imide species (with mass tolerance of ± 0.01) were background-subtracted and integrated on Mnova.

MTase Glo assay

MTase Glo assay (Promega), which detects the level of SAH, was performed as described in the supplied protocol. In each well of a solid white 96-well plate, the reaction buffer (1×: 20 mM Tris-Cl pH 8.0, 50 mM NaCl, 1 mM EDTA, 3 mM $MgCl_2$, 0.1 mg/mL BSA, 1 mM DTT), SAM (1×: 30 µM) and water were mixed to make the 2× substrate dilution solution. Then, the substrate solution was prepared by mixing the substrate at the highest desired concentration with the reaction buffer and SAM at 2× concentration. 1:2 serial dilutions of the substrate solution were performed by mixing equal volumes of 2× substrate dilution solution and substrate solution. 10 µL of the serial dilution of substrates and 10 µL of $His_6$-PCMT1 (1×: 0.05 µM for isoD peptide and 5 µM for C-terminal N peptide) were added to each well to initiate the reaction. The plate was incubated at 37 °C for 30 min for isoD peptide and for 3 h for C-terminal N peptide. The reaction was quenched by adding 5 µL 0.5% trifluoroacetic acid and the plate was incubated for 5 min at room temperature. MTase Glo reagent (5 µL of 6× stock) was added to each well, and the plate was centrifuged (1000 × g, 2 min) and incubated for 30 min at room temperature. MTase Glo detection solution (30 µL of 2× stock) was added to each well and the plate was again centrifuged (1000 × g, 2 min) and incubated for 30 min at room temperature. Before the substrates were tested, a standard curve was generated with the luminescence measurements of SAH incubated with the MTase Glo reagent and detection solution to determine the slope $k_{SAH}$. The measurements from the substrates were graphed and fitted to the Michaelis-Menten model in Prism 10 to determine the $y_{max}$ (maximum luminescence) and $K_m$ values. $K_{cat}$ was then calculated using the equations below:

$$v_{max} = \frac{y_{max}}{k_{SAH} \times t_{incubation}} \qquad K_{cat} = \frac{v_{max}}{[PCMT1, \mu M]}$$

TR-FRET displacement assay

Peptide or recombinant protein (10 µM) was incubated with $His_6$-tag-free PCMT1 (10 µM) and SAM (200 µM) in 50 mM Tris-Cl buffer, pH 7.4 at 37 °C for 14 h. Triton X-100 was added to the mixture to final concentration of 0.1% for ease of dispense.

Anti-$His_6$ antibody (Abcam ab18184) was labeled with CoraFluor-1-Pfp ester as previously described to generate the TR-FRET donor.[3] TR-FRET experiments were performed in Proxiplate-384 Plus (VWR PERK6008280) with 15 µL assay volume. TR-FRET measurements were acquired on a SpectraMax iD5 plate reader with SoftMax Pro software version 7.1.2, with the following settings: 350 nm excitation, 490 nm (Tb), and 520 nm (FITC) emission, 110 flashes per read, 0.05 ms excitation time, 0.05 ms delay and 0.2 ms integration. The TR-FRET ratio was taken as the 520/490 nm intensity ratio for each well.

To determine the tracer $K_D$, 5 nM CoraFluor-1-labeled anti-$His_6$ antibody and 10 nM $His_6$-CRBN/DDB1 complex were mixed in assay buffer (25 mM HEPES, 150 mM NaCl, 0.5 mg/mL BSA, 0.005% TWEEN 20, pH 7.5). Thal-FITC was first added in a serial dilution (1:2 titration, 8-point, $c_{max}$ = 1.33 µM) using an HP D300 digital dispenser and allowed to equilibrate for 10 min at room temperature before TR-FRET measurements were taken. The $K_D$ determined from the one-site model calculated in Prism 10 was used in the equation below to adjust for a two-site model due to the presence of the bivalent anti-$His_6$ antibody:

$$K_{D,\,adj} = K_D \times (1 + \sqrt{2})$$

For ligand displacement assays, 5 nM CoraFluor-1-labeled anti-$His_6$ antibody,10 nM $His_6$-CRBN/DDB1 complex, and 50 nM Thal-FITC were mixed in the assay buffer. In all cases, the incubation mixture was added in a serial dilution (1:3 titration, 8-point, $c_{max}$ = 1 µM for the peptide or protein substrate) using an HP D300 digital dispenser and allowed to equilibrate for 10 min at room temperature before TR-FRET measurements were taken. All experiments were performed in technical triplicates. Background signal (bottom) was determined from wells containing 100 µM 5-$NH_2$-lenalidomide to represent complete displacement. The assay ceiling (top) was defined by a no-ligand control. Data were background-subtracted, normalized, and fitted to a four-parameter dose-response model [log(inhibitor) vs. response – Variable slope (four parameters)] Prism with constraints of Top = 1 and Bottom = 0 in 10 to derive the ligand $IC_{50}$ values. Ligand $K_D$ values were calculated using Cheng-Prusoff principles according to the equation below:

$$K_D = \frac{IC_{50}}{1+\frac{[S]}{K_x}}$$

where $IC_{50}$ is the measured $IC_{50}$ value, [S] is the concentration of fluorescent tracer, and $K_X$ is the adjusted $K_D$ of the fluorescent tracer.

Biolayer interferometry

The binding affinity between CRBN and GLUL-cN was measured using the Octet Red 96 (ForteBio, Pall Life Sciences) following the manufacturer’s procedures. The optical probes were coated with anti-GST antibody, then loaded with 500 nM GST-tagged GLUL-cN. Subsequently, the probes were quenched with 0.5 mM biocytin or 1 µM GST protein prior to kinetic binding analysis. The reactions were carried out in black 96 well plates maintained at 30°C. The reaction volume was 200 µL in each well. The binding buffer contained 25 mM HEPES, pH 7.4, 100 mM NaCl, 1 mM TCEP, 0.1% Tween-20, and 0.05 mg/mL BSA. CRBN was used as the analyte at various concentrations. Binding kinetics of the analyte at different concentrations were measured simultaneously. The data were analyzed using the Octet data analysis software. The $k_{on}$

and $k_{off}$ values were used to calculate the dissociation constant ($K_D$) with kinetic analysis of direct binding. All BLI experiments were repeated at least 2 times.

In vitro ubiquitination

A reaction mixture containing 1.7 μM Venus-GLUL without or with PCMT1 activation, 1 μM CRBN-DDB1-DDA1, 0.6 μM NEDD8~CUL4-RBX1, 0.6 μM E2 (UBCH5), 0.3 μM UBE1, 50 μM ubiquitin, and 2 mM ATP/10 mM $MgCl_2$ was incubated at 37 °C for 30 min. Control experiments were performed by systematically omitting one component of the reaction at a time. The reaction mixtures were resolved by a 10% SDS–PAGE gel. The GLUL signal was monitored by detecting Venus absorbance at 488 nm. The absorbance at 647 nm was used to detect the protein markers.

Generation of PCMT1 knockout HEK293T cells by CRISPR/Cas9

The CRISPR/Cas9 plasmids, which contain the Cas9 and puromycin resistance genes and either a non-targeting sgRNA sequence or a sgRNA sequence targeting PCMT1 (Sequences 1–2), were purchased from Genscript. The transfection and selection procedures were adapted from Ran and co-workers.[4] $1.5\times10^5$ WT HEK293T cells were seeded in 24-well plates containing 0.5 mL DMEM+/– (supplemented with FBS but without antibiotics). For each well, 0.5 μg CRISPR/Cas9 plasmid was diluted in 50 μL Opti-MEM I, then 0.5 μL TransIT-Pro was added. The mixture was incubated for 20 min at room temperature, then added dropwise into each well. After 24 h, 0.5 mL fresh DMEM+/– was added into each well. After another 24 h, the media was changed to 1 mL DMEM+/+ containing 2 μg/mL puromycin and further incubated for 72 h to select the cells that took up the puromycin resistance gene. The selected cells were expanded and validated for PCMT1 knockout by Western blotting.

Electroporation of GFP-GLUL

For sampled pre-treated with PCMT1, $His_6$-GFP-GLUL (WT or N373A, 25 μM) was incubated with $His_6$-PCMT1 (50 μM) and SAM (500 μM) in 50 mM Tris-Cl buffer, pH 7.4 at 37 °C for 14 h. The incubation mixtures were directly used for electroporation. For samples without PCMT1 pre-treatment, $His_6$-GFP-GLUL (WT or N373A) was diluted to 50 μM in PBS.

HEK293T cells were grown to 80–90% confluency in DMEM+/+ prior to electroporation. Cells were detached by trypsinization as necessary and washed with PBS, then resuspended in PBS and counted. For each sample, an equal number of cells ($7.0\times10^5$) were aliquoted into Eppendorf tubes and pelleted. Electroporation mixes were prepared for each sample type. Electroporation mixes were prepared by combining 41.6 μL Neon buffer R containing DMSO or 100 μM lenalidomide, 4.2 μL of 50 μM mCherry protein as the internal standard, and 4.2 μL of 50 μM GFP-GLUL (WT or N373A) or PCMT1 incubation mixture. A mock electroporation mix was also prepared by substituting the mCherry and GFP-GLUL with PBS. Immediately prior to each electroporation, the PBS was removed from the pelleted cells, and the pellet was resuspended in 12 μL of electroporation mix. The sample was taken up into a 10 μL tip attached to a Neon pipette, and the pipette tip was submerged in a Neon cuvette containing 3 mL Neon buffer E2. The sample was then electroporated (1150 V, 20 msec, 2 pulses). The cells were then dispensed into 100 μL PBS pre-warmed to 37 °C and flicked to mix. This process was repeated for each sample, with each tip reused within triplicate samples. Cells were then pelleted by centrifugation, and the supernatant was discarded. Cells were resuspended in 50 μL colorless trypsin-EDTA solution and incubated at 37 °C for 3 min. Trypsinization was quenched by addition of 0.5 mL colorless DMEM+/+ containing DMSO or 100 μM lenalidomide, and the mixture was transferred to a well of a 24-well plate. The samples were then incubated at 37 °C, 5% $CO_2$ for the desired time. After incubation, the media was removed, and each sample was resuspended in 500 μL PBS. The samples were then analyzed by flow cytometry (PE Texas Red and FITC channels on FACSymphony A3 Lite). To calculate normalized GFP level, the arithmetic mean of the GFP and mCherry fluorescence intensities for each sample were corrected by subtracting the corresponding intensity values in the mock sample; then, these corrected values were used to calculate a GFP/mCherry fluorescence intensity ratio for each sample. Finally, the resulting values were normalized to the mean ratio for GFP-GLUL (N373A) in the presence of exogenous or endogenous PCMT1 without lenalidomide competition.

<u>Global quantitative proteomics sample preparation</u>

The proteomics sample preparation protocol was adapted from Donovan and coworkers,[5] and the off-line fractionation protocol was adapted from Batth and coworkers.[6] Cellular samples were prepared in biological quadruplicate for each of the 4 conditions (sgCtrl + DMSO, sgCtrl + lenalidomide, PCMT1-KO + DMSO, PCMT1-KO + lenalidomide). $3.0\times10^6$ cells were seeded in 6-well plates containing 1.5 mL DMEM+/+ and treated with DMSO or 100 μM lenalidomide. After 24 h, the media was replaced with fresh DMEM+/+ containing DMSO or 100 μM lenalidomide and further incubated at 37 °C for 24 h (total treatment time: 48 h). Cells were collected by trypsinization, washed with PBS, lysed by probe sonication (5 sec on, 3 sec off, 15 sec total on time, 11% amplitude) in lysis buffer (8 M urea, 50 mM NaCl, 50 mM HEPES, 1×protease/phosphatase inhibitor cocktail, pH 7.4), and cleared by centrifugation (21,000 × g, 4 °C, 10 min). Mouse brain samples were prepared from the hippocampus from 4 WT and 4 Crbn-KO individual mice. The hippocampus from each mouse was transferred into a 2.0 mL tube using forceps, to which 120 μL lysis buffer was added. Tissues were lysed by probe sonication (5 sec on, 15% amplitude) and cleared by centrifugation (21,000 × g, 4 °C, 10 min).

After total protein quantification by BCA protein assay, the cell or tissue lysates were diluted to 2 mg/mL with additional lysis buffer. The samples (200 μg protein per sample) were reduced by adding dithiothreitol (5 mM) and incubating at 24 °C for 30 min, then alkylated by adding iodoacetamide (15 mM) and incubating at 24 °C for 30 min protected from light. Proteins were precipitated by methanol-chloroform precipitation. Four volumes of methanol were added to the lysate, followed by one volume of chloroform, and finally three volumes of water. The mixture was vortexed and centrifuged at 14,000 × g for 5 min at 4 °C, and the upper layer was carefully removed.

The precipitated protein was then washed with three volumes of methanol and centrifuged at 14,000 × g for 5 min at 4 °C, and the resulting protein pellet was dried for 10 min in a vacufuge. The protein pellet was resuspended in 25 µL of 4 M urea, 50 mM HEPES, pH 7.4, after which 75 µL of 200 mM HEPES, pH 7.4 was added to lower the final urea concentration to 1 M. The samples were first digested with 2 µg of Lys-C for 4 h at 22 °C. The mixture was then diluted by addition of 100 µL of 200 mM HEPES, pH 7.4, and further digested with 4 µg of trypsin for 16 h at 37 °C. For each sample, half of the digested peptide mixture (ca. 100 µg peptides) was taken for labeling with 400 µg TMTpro 16-plex reagent. The labeling reactions were incubated for 1.5 h at 22 °C, then quenched by adding 20 µL of 1 M Tris-Cl, pH 7.6 and incubating for 15 min at 22 °C. The channels were combined, dried in a vacufuge, and desalted using a Pierce peptide desalting spin column (Thermo) according to the manufacturer's protocol. The desalted sample was offline fractionated into 80 fractions by high pH reverse-phase HPLC (Agilent 1260 Infinity II) through an Aeris peptide XB-C18 column (Phenomenex 00G-4507-E0) with mobile phase A containing 5% acetonitrile and 10 mM ammonium bicarbonate, and mobile phase B containing 90% acetonitrile and 1 mM ammonium bicarbonate (both pH 7.4). Fractions (0.7 mL each) were collected using a fraction collector in a deep 96-well plate. Samples were initially loaded onto the column at 1 mL/min for 4 min, after which the fractionation gradient was set as follows: 1% B to 27% B over 50 min, ramp to 60% B over 4 min, and ramp to 70% B over 2 min. Fraction collection was stopped at this point, and the gradient was held at 70% B for 5 min before being ramped back to 1% B to wash and equilibrate the column. The 80 resulting fractions were pooled into 20 final fractions in a non-continuous manner by combining fractions n, n+20, n+40, and n+60. The 20 final fractions were dried, then resuspended in 40 µL of 0.1% formic acid.

Proteomics mass spectrometry acquisition procedures

The resuspended samples (2 µL) were run on an Orbitrap Eclipse Tribrid Mass Spectrometer coupled with a Vanquish Neo HPLC system (Thermo) at the Harvard Center for Mass Spectrometry. The peptides were first trapped on a trapping cartridge (300 µm x 5 mm PepMap™ Neo C18 Trap Cartridge, Thermo) prior to separation on a silica-chip-based micropillar column (µPAC, C18 pillar surface, 50 cm bed, Thermo). The column oven temperature was maintained at 35 °C. Peptides were eluted using a multi-step gradient at a flow rate of 300 nL/min over 180 min. The mobile phase A and weak wash liquid was water with 0.1% formic acid, the mobile phase B was acetonitrile with 0.1% formic acid, and the strong wash liquid was 80% acetonitrile with 0.1% formic acid. The mobile phase gradient consisted of a linear 125 min gradient from 2% to 20% mobile phase B, followed by a 24 min increase to 35% B, a further 10 min increase to 95% B, and a 20 min plateau phase at 95% B. The autosampler temperature was 7 ºC. The column "Fast equilibration" was enabled. The Orbitrap Eclipse MS was operated in DDA mode with 3 s cycle time. The electrospray ionization was in positive mode with voltage of 2.1 kV and the capillary temperature at 275 ºC. Dynamic exclusion was enabled with a mass tolerance of 10 ppm and exclusion duration of 60 s. Full scan was performed in the range of 400–1600 m/z at a resolution of 120,000, RF lens 30%, normalized AGC target 200%, and maximum injection time set to auto. Precursors were isolated in a window of 0.7 m/z and charge states of 2–6. MS2 fragmentation was performed by HCD using a normalized collision energy of 38% at a resolution of 50,000. The normalized AGC target was set to 250%, and the maximum ion injection time was set to 200 ms.

Mass spectrometry data analysis for protein level quantitation

Analysis was performed in Thermo Scientific Proteome Discoverer version 2.4.1.15. The raw data were searched against SwissProt human (Homo sapiens) protein database (21 February 2019; 20,355 total entries) or SwissProt mouse (Mus musculus) protein database (30 May 2018; 17,372 total entries) and contaminant proteins using the Sequest HT algorithm. Searches were performed with the following guidelines: spectra with a signal-to-noise ratio greater than 1.5; mass tolerance of 20 ppm for the precursor ions and 0.02 Da for the fragment ions; full trypsin digestion; fewer than 2 missed cleavages; static carboxyamidomethylation of cysteine residues (+57.021 Da); static TMTpro 16-plex labeling (+304.207 Da) at lysine residues and N-termini; variable oxidation on methionine residues (+15.995 Da); variable dehydration on asparagine and glutamine residues at protein C-terminus only (–18.015 Da). The TMT reporter ions were quantified using the Reporter Ions Quantifier node and normalized to the total peptide amount. Peptide spectral matches (PSMs) were filtered using a 1% false discovery rate (FDR) using Target Decoy PSM Validator. For the quantification of total proteins, the data were further filtered to include only master proteins with high protein FDR confidence, at least 3 unique peptides, and exclude all contaminant proteins. The abundance ratios and their associated p-values were calculated by one-way ANOVA with TukeyHSD post-hoc test.

Glutamine starvation and refeeding

For glutamine starvation experiments, $0.8\times10^6$ HEK293T cells per condition were seeded in 6-well plates containing 2 mL DMEM+/+ without glutamine supplementation and incubated for the desired period of time until collection. For glutamine refeeding experiments, $9\times10^6$ HEK293T cells were seeded in 15 cm plates containing 25 mL DMEM+/+ without glutamine supplementation and incubated for 40–48 h. Then, these glutamine-starved cells were trypsinized, and $2\times10^6$ cells per condition were seeded in 6-well plates containing 1 mL DMEM+/+ with 4 mM glutamine. The samples were further incubated for the desired period of time. Cells were then collected and analyzed by Western blotting.

RT-qPCR

Total RNA was extracted from cells or mouse brain tissues using the RNeasy kit or RNeasy lipid tissue kit (both Qiagen), respectively. RNA samples were diluted with water to the same final concentration of 50–100 ng/µL. Reverse transcription and real-time PCR were performed with the Luna universal one-step RT-qPCR kit (NEB) using the supplied protocol on the iQ5 Multicolor Real-Time PCR Detection System. The relative mRNA level was calculated using the $2^{(-\Delta\Delta Cq)}$ method with β-actin as the reference gene. Sequences 3–16 (3–6 for human mRNA, 7–16 for mouse mRNA) were used as primers.

Cerebody enrichment

Cell samples were lysed in Pierce IP lysis buffer supplemented with 1× protease/phosphatase inhibitor cocktail. Tissue samples were lysed by sonication (5 sec on, 15% amplitude) in T-PER buffer (Thermo) supplemented with 1× protease/phosphatase inhibitor cocktail. Lysates were then cleared by centrifugation (20,000 × g, 4 °C, 10 min). After total protein quantification by BCA protein assay, the lysates were diluted to 2 mg/mL. For each sample, 800 µg of lysate was transferred to a new tube, to which cerebody (5 µM final concentration) and 50 µL of TBS-washed anti-FLAG M2 magnetic beads (Sigma) were added. The samples were incubated on a roller at room temperature for 80 min. After pelleting the beads using a magnetic rack and removing the supernatant, 150~400 µL of PBS or 1% Triton X-100 in PBS was added, and the beads were washed either by pipetting up-and-down or vortexing at the lowest speed for 1-5 min (depending on the target protein and sample type). After again pelleting the beads using a magnetic rack and removing the supernatant, the samples were eluted by adding 50 µL of 150 ng/µL 3× FLAG peptide in TBS or 50 µL of 200 µM lenalidomide in PBS and incubating at room temperature for 1 h with agitation. The beads were pelleted, and the eluates were collected for Western blotting or further analysis.

Selected ion monitoring of GLUL C-terminal peptides

To generate the heavy internal standard, TGDEP*FQYKN (10 µM) was incubated with $His_6$-PCMT1 (10 µM) and SAM (200 µM) in 50 mM Tris-Cl buffer, pH 7.4 at 37 °C for 14 h. The incubation mixture was then filtered through a 10 kDa MWCO spin filter to remove the $His_6$-PCMT1.

For recombinant protein samples, a 100 µL reaction mixture containing $His_6$-GLUL (10 µM), $His_6$-PCMT1 (20 µM), and SAM (200 µM) in 50 mM Tris-Cl buffer, pH 7.4 was incubated at 37 °C for 14 h.

For samples prepared by cerebody enrichment, cerebody enrichment was performed as described above, with the following modifications: cell lysates were diluted to 9 mg/mL; 3.6 mg of lysate was used per sample; 200 µL of beads were used per sample; 100 µL of 150 ng/µL 3× FLAG peptide in TBS was used to elute.

To prepare samples for mass spectrometry, 100 µL of 10% SDS, 100 mM Tris-Cl, pH 7 buffer was added to each sample, after which 20 µL of 27.5% phosphoric acid was added and the samples were vortexed. Next, 1.32 mL of S-Trap binding buffer (100 mM triethylammonium bicarbonate in 90% methanol, pH 7.1) was added to each sample, then the samples were applied to S-Trap micro columns (Protifi) connected to a vacuum manifold. Each column was washed three times with 1 mL S-Trap binding buffer, after which the columns were centrifuged at 4000 × g for 1 min to remove residual buffer. Each sample was digested by adding 2 µg Glu-C resuspended in 20 µL 100 mM ammonium bicarbonate, pH 7 to the headspace of the column and incubating at 37 °C for 3 h. The digests were then eluted by adding 40 µL of 100 mM ammonium bicarbonate, pH 7 and centrifuging at 4000 × g for 1 min, then again with 40 µL of 0.2% formic acid, then once again with 40 µL of 0.2% formic acid in 50% acetonitrile. The eluates from each sample were pooled and concentrated to dryness in a vacufuge, then resuspended in 300 µL 0.1% trifluoroacetic acid. At this stage, 4 µL of the heavy standard mixture was spiked into each sample. The samples were then desalted using Pierce peptide desalting spin columns (Thermo) according to the manufacturer's protocol and concentrated to dryness in a vacufuge, then resuspended in 20 µL of 0.1% formic acid.

The resuspended samples (1 µL) were injected on an Orbitrap Eclipse Tribrid Mass Spectrometer coupled with a Vanquish Neo HPLC system (Thermo) at the Harvard Center for Mass Spectrometry. The peptide samples were first trapped on a trapping cartridge (300 µm × 5 mm PepMap™ Neo C18 Trap Cartridge, Thermo) prior to separation on a silica-chip-based micropillar column (µPAC, C18 pillar surface, 50 cm bed, Thermo). The column oven temperature was maintained at 35 °C. Peptides were eluted using a gradient at a flow rate of 300 nL/min over 90 min. The mobile phase A and weak wash liquid was water with 0.1% formic acid, the mobile phase B was acetonitrile with 0.1% formic acid, and the strong wash liquid was 80% acetonitrile with 0.1% formic acid. The mobile phase gradient consisted of a linear 69 min gradient from 1% to 33% mobile phase B, followed by a 13 min increase to 46% B, then a further 8 min plateau phase at 95% B. The autosampler temperature was 7 ºC. The Orbitrap Eclipse MS was operated in positive electrospray ionization mode with a voltage of 2.1 kV and the capillary temperature at 275 ºC. FAIMS compensation voltages was set at -45V with a cycle time of 3 s. The target selected ion monitoring (tSIM) was performed at a resolution of 120,000 with an isolation window of 1 m/z, RF lens of 30%, normalized AGC target at 200%, and maximum injection time of 300 ms. The z = 2 ions of four peptides of interest were monitored: TGDEPFQYKN (m/z = 599.7724), TGDEPFQYKcN (m/z = 590.7671), TGDEP*FQYKN (m/z = 602.7793), and TGDEP*FQYKcN (m/z = 593.7740). Dynamic exclusion was enabled with a mass tolerance of 10 ppm and exclusion duration of 1 s. Fragmentation of target precursors was performed by ten MS2 scans in orbitrap using higher-energy collisional dissociation (HCD) followed by one MS2 scan in ion trap using collision-induced dissociation (CID). HCD was set at a normalized collision energy (NCE) of 30% at a resolution of 60,000 with a normalized AGC target at $5\times10^4$ and a maximum ion injection time of 150 ms. CID was set at a collision energy of 35 with a normalized AGC target at $1\times10^4$ and a maximum ion injection time of 35 ms.

The raw MS1 chromatograms were extracted for each peptide of interest with a mass tolerance of ± 0.005 m/z. For each species, the peak area at the expected retention time was integrated and quantified using Xcalibur Qual Browser version 3.0.63. In each sample, the cN and N peptides were normalized to the respective heavy internal standards, then these normalized values were used to compare the cN and N peptide signals between samples.

X-ray crystallography

CRBN-DDB1 and PFQYKcN were mixed at a 1:3 molar ratio and concentrated to 10 mg/mL for crystallization at 25 °C by the hanging-drop vapor-diffusion method. 1 µL of protein sample was mixed with an equal volume of reservoir solution (8% v/v TacsimateTM pH 6.0, 20% w/v Polyethylene glycol 3,350). The largest crystal was harvested and flash-frozen in the crystallization buffer supplemented with 20% glycerol at –170 °C. The X-ray diffraction dataset was collected at the BL8.2.1 beamline at the Advanced Light Source in Berkeley and was integrated and scaled by XDS.[7] The complex structure was solved by molecular replacement using the program Phaser-MR of PHENIX[8] with the CRBN-DDB1 structural model (PDB: 5FQD) as search template. The complex structure model was generated by AutoBuild and refined and ligand-fitted using COOT[9] and PHENIX.[8] PyMOL (The PyMOL Molecular Graphics System, Version 2.0 Schrödinger, LLC) was used to generate figures.

Quantification of glutamine and glutamate levels in tissue samples

Glutamine and glutamate levels were measured using the Glutamine/Glutamate-Glo assay kit (Promega). Tissue samples (3-5 mg) were lysed by sonication (5 sec on, 15% amplitude) in 1.125 mL of an 8:1 (v:v) mixture of 50 mM Tris, pH 7.5 and 600 mM HCl, after which 125 µL of 600 mM Tris, pH 8.5 was added to each sample. Lysates were then cleared by centrifugation (20,000 × g, 4 °C, 10 min). A small aliquot from each sample was set aside and neutralized to pH 7 by addition of 600 mM Tris, pH 8.5, then this neutralized sample was used for total protein quantification by BCA protein assay. The remaining lysates were diluted to the same total protein concentration with an 8:1:1 (v:v:v) mixture of 50 mM Tris, pH 7.5, 600 mM HCl, and 600 mM Tris, pH 8.5. The lysates were then analyzed according to the manufacturer's protocol. Each tissue sample was analyzed in triplicate.

Quantification of ammonium levels in tissue samples

Ammonium levels were measured using the Ammonia Assay Kit (Rapid) (Megazyme). Tissue samples were lysed by sonication (5 sec on, 15% amplitude) in 150 µL PBS, after which the lysates were cleared by centrifugation (20,000 × g, 4 °C, 10 min). Total protein concentration in each sample was measured by BCA assay, then all lysates were diluted to the same total protein concentration. The samples were deproteinized by adding cold perchloric acid to a final concentration of 1 M, vortexing, and incubating on ice for 5 min. Samples were centrifuged (20,000 × g, 4 °C, 10 min) to remove the precipitated protein. Next, the samples were neutralized to pH 6.5-8.0 by addition of cold 2 M KOH then incubated on ice for 20 min to allow the perchloric acid to precipitate, which was then removed by centrifugation (20,000 × g, 4 °C, 10 min). The deproteinized and neutralized lysates were then analyzed according to the manufacturer's protocol. Each tissue sample was analyzed in duplicate.

Mouse studies

Adult (ca. 3-month-old) *Crbn*-null mice (CMV-Cre:*Crbn*$^{fl/fl}$ mice, herein referred to as Crbn$^{-/-}$ mice) and *Crbn*$^{fl/fl}$ littermates (herein referred to as WT mice) of both sexes (at approximately 1:1 ratio) were used throughout the study. All the animal procedures were performed in accordance with the guidelines and with the approval of the Animal Welfare Committee of Complutense University of Madrid and Comunidad de Madrid, and in accordance with the directives of the European Commission. The ARRIVE guidelines were followed as closely as possible. Mice were housed, handled, and assigned to the experimental groups as described.[10]

For the induction of excitotoxic seizures, kainic acid (KA, Sigma) was dissolved in isotonic saline, pH 7.4, and injected i.p. at 20 mg/kg body weight. Immediately after, mice were placed in clear plastic cages and monitored continuously for 135 min to assess seizure behavior. The highest convulsive score reached at each 5-min interval was given according to a modified Racine scale as described.[11,12] Mice were habituated to the experimenter and the experimental room prior to the test. The test was conducted during the early light phase under dim illumination (< 50 lx in the center of the corresponding cage) and video-recorded for subsequent analysis by a different trained observer, who remained blind towards the genotype of the animal.

#### III. Materials and instrumentation

General supplies

- (*R*, *S*)-Lenalidomide (BioVision 1862-25)
- MLN4924 (Selleck Chemicals S7109)
- TAK-243 (Selleck Chemicals S8341)
- MG132 (Selleck Chemicals S2619)
- Zeba 7 kDa desalting columns (0.5 mL and 5 mL, ThermoFisher 89882 and 89892)
- Dulbecco's Modified Eagle's Medium (DMEM) with 4 mM glutamine (ThermoFisher 11995065)
- DMEM, glutamine-free (ThermoFisher 10313021)
- Trypsin-EDTA (Fisher Scientific 25200114)
- Fetal bovine serum (FBS) (Peak Serum PS-FB2)
- Penicillin-streptomycin (100×) (Lonza 17-602E)
- Pierce IP lysis buffer (ThermoFisher 87788)
- T-PER Tissue Protein Extraction Reagent (ThermoFisher 78510)
- Protease/phosphatase inhibitor cocktail (100×) (Cell Signaling Technology 5872S)
- Protease inhibitor cocktail (Sigma-Aldrich 11873580001, 1 tablet dissolved in 2 mL water for a 25× stock solution)
- Thrombin CleanCleave Kit (Sigma RECOMT-1KT)

- Ni-NTA resin (G-Biosciences 786-940)
- Glutathione sepharose 4B resin (Cytiva 17075601)
- BCA solution (BCA Reagent A) (VWR 786-847)
- Copper Solution (BCA Reagent B) (VWR 76825-860)
- 12% Criterion TGX precast gels (Bio-Rad 5671044)
- iBlot 2 nitrocellulose transfer stack (Invitrogen IB23001; IB23002)
- Vivaspin spin concentrators (Cytiva)
- Opti-MEM I Reduced Serum Medium (ThermoFisher 31985070)
- TransIT-Pro Transfection Reagent (Mirus MIR 5760)

Cloning reagents

- Q5 site-directed mutagenesis kit (New England BioLabs E0552S)
- QuikChange Lightning Site-directed Mutagenesis Kit (Agilent 210518)
- HiFi DNA Assembly Kit (New England BioLabs E5520S)

Mass spectrometry

- Triethylammonium bicarbonate buffer (Sigma-Aldrich T7408-100ML)
- Pierce high pH reversed-phase peptide fractionation kit (ThermoFisher 84868)
- Lys-C (Promega VA1170)
- Trypsin (Promega VA5117)
- Glu-C (Promega V1651)
- TMTpro 16-plex (ThermoFisher A44520)
- Pierce Peptide Desalting Spin Column (ThermoFisher 89852)
- S-trap Micro Column (Protifi C02-micro-40)
- 96-well plate, 1.0 mL, round wells (Agilent, 5043-9305)

MTase Glo assay

- MTase Glo assay (Promega V7601)
- 96-well plate, white opaque (ThermoFisher 15042)

TR-FRET

- Proxiplate-384 Plus (VWR PERK6008280)

Electroporation

- Neon Transfection System 10 µL Kit (ThermoFisher MPK1096)
- Neon Transfection Tubes (ThermoFisher MPT100)
- DMEM with 4.5 g/L Glucose, without phenol red (Lonza, 12-917F)
- Trypsin-EDTA (0.5%), no phenol red (ThermoFisher, 15400054)

RT-qPCR

- RNeasy mini kit (Qiagen 74104)
- RNeasy lipid tissue mini kit (Qiagen 74804)
- Luna Universal One-step RT-qPCR kit (New England BioLabs E3005)
- PCR plate, 96-well (ThermoFisher AB0600)

Metabolite measurements

- Glutamine/Glutamate-Glo Assay (Promega J8021)
- Ammonia Assay Kit (Rapid) (Megazyme K-AMIAR)

Purchased synthetic peptides

- FQYKN (Genscript)
- EQMQN (Genscript)
- DIHAQ (Genscript)
- KILAQ (Genscript)
- DDKCQ (Genscript)
- YKN (Genscript)
- QYKN (Genscript)
- PFQYKN (Genscript)
- EPFQYKN (Genscript)
- DEPFQYKN (Genscript)
- GDEPFQYKN (Genscript)
- TGDEPFQYKN (Genscript)
- PFQYKA (Genscript)
- PFQYKQ (Genscript)
- PFQYAN (Genscript)
- PFQAKN (Genscript)

- PFAYKN (Genscript)
- PAQYKN (Genscript)
- AFQYKN (Genscript)
- PFQYFN (Genscript)
- PFQYPN (Genscript)
- PFQYYN (Genscript)
- PFQYLN (Genscript)
- PFQYEN (Genscript)
- PFQYQN (Genscript)
- PFQYIN (Genscript)
- PFQYSN (Genscript)
- PFQYGN (Genscript)
- PFQFKN (Genscript)
- PFQPKN (Genscript)
- PFQLKN (Genscript)
- PFQEKN (Genscript)
- PFQKKN (Genscript)
- PFQQKN (Genscript)
- PFQIKN (Genscript)
- PFQSKN (Genscript)
- PFQGKN (Genscript)
- TGDEP*FQYKN (P* = Pro-$^{13}C_5$,$^{15}N$) (Genscript)
- HHQKN (Genscript)
- HQQKN (Genscript)

Bacterial strains

- *E. coli* 5-alpha Competent (High Efficiency) (New England BioLabs C2987H)
- *E. coli* BL21 (DE3) (New England BioLabs C2527H)

Mammalian cell lines and tissues

HEK293T, SK-N-SH, and MEF cells were obtained from American Type Culture Collection (ATCC). HEK293FT cells stably expressing FLAG-CRBN (HEK-CRBN cells) were kindly provided by the Deshaies Lab (California Institute of Technology). Neuro-2a cells and mouse brain parts were kindly provided by the Guzmán Lab (Complutense U of Madrid).

Antibodies

| No. | Antibody Name | Host Species | Blocking Buffer | Supplier | Catalog# |
|---|---|---|---|---|---|
| 1 | $His_6$ | Mouse mAb | For TR-FRET | Abcam | ab18184 |
| 2 | HA-Tag | Rabbit mAb | 5% milk/TBST | Cell Signaling Technology | 14031S |
| 3 | Vinculin (VCL) | Mouse mAb | 5% milk/TBST | Bio-Rad | MCA465GA |
| 4 | GLUL | Rabbit pAb | 5% milk/TBST | Bethyl Laboratories | A305-323A |
| 5 | CRBN | Rabbit mAb | 5% BSA/TBST | Cell Signaling Technology | 71810 |
| 6 | PCMT1 | Mouse mAb | 5% milk/TBST | Santa Cruz Biotechnology | sc-100977 |
| 7 | PPA1 | Rabbit pAb | 5% milk/TBST | Proteintech | 14985-1-AP |
| 8 | Anti-mouse-HRP | Goat pAb | Secondary antibody | Rockland Immunochemicals | 610-1302 |
| 9 | Anti-rabbit-HRP | Goat pAb | Secondary antibody | Rockland Immunochemicals | 611-1302 |
| 10 | Anti-mouse-IRDye® 800CW | Goat pAb | Secondary antibody | LI-COR Biosciences | 925-32210 |

Plasmids

| No. | Plasmid name | Source |
|---|---|---|
| 1 | pET30b-$His_6$-PCMT1(WT) | Addgene 34852 |
| 2 | pET30b-$His_6$-PCMT1(S60A) | This work |
| 3 | pET19b-GST-GLUL(WT) | This work |
| 4 | pET19b-GST-GLUL(N373A) | This work |

| | | |
|---|---|---|
| 5 | pET19b-$His_6$-GFP | Previous work |
| 6 | pET19b-$His_6$-GFP-GLUL(WT) | This work |
| 7 | pET19b-$His_6$-GFP-GLUL(N373A) | This work |
| 8 | pET19b-$His_6$-GLUL(WT) | This work |
| 9 | pET19b-$His_6$-GLUL(N373A) | This work |
| 10 | pX459-sgCtrl | Genscript |
| 11 | pX459-sgPCMT1 | Genscript |
| 12 | pET28a-cerebody | Previous work |
| 13 | pAL-$His_6$-CRBN | This work |
| 14 | pAL-$His_6$-CRBN (N351A) | This work |
| 15 | pAL-$His_6$-CRBN (Y355A) | This work |
| 16 | pAL-$His_6$-CRBN (H357A) | This work |
| 17 | pAL-$His_6$-CRBN (W380A) | This work |
| 18 | pAL-$His_6$-CRBN (W386A) | This work |
| 19 | pAL-$His_6$-CRBN (H397A) | This work |
| 20 | pAL-$His_6$-CRBN (W400A) | This work |
| 21 | pAL-$His_6$-CRBN (V388I) | This work |
| 22 | pET-GST-GLUL (C-end 6-mer peptide) | This work |
| 23 | pACE-$His_6$-Venus-GLUL (full-length) | This work |

<u>DNA sequences</u>

| No. | Sequence name | Sequence (5' to 3') |
|---|---|---|
| 1 | sgCtrl | GCTTAGTTACGCGTGGACGAAGG |
| 2 | sgPCMT1 | GATTGTGGATTAGCTCCGAG |
| 3 | hGLUL-F | AAGAGTTGCCTGAGTGGAATTTC |
| 4 | hGLUL-R | AGCTTGTTAGGGTCCTTACGG |
| 5 | hActin-F | CATGTACGTTGCTATCCAGGC |
| 6 | hActin-R | CTCCTTAATGTCACGCACGAT |
| 7 | mGlul-F | TGAACAAAGGCATCAAGCAAATG |

| | | |
|---|---|---|
| 8 | mGlul-R | CAGTCCAGGGTACGGGTCTT |
| 9 | mPpa1-F | AGTACCGCGTCTTCCTCAAAA |
| 10 | mPpa1-R | GACCAGCGTGGAACCTCAAC |
| 11 | mCrbn-F | TACCTGGGAGCTGATATGGAGG |
| 12 | mCrbn-R | TTCCGCACCATGCTGACTTC |
| 13 | mPcmt1-F | GGAGGTGGTCTCACTCTTGG |
| 14 | mPcmt1-R | AGGTTGTGGATTAGCTCCGAG |
| 15 | mActin-F | GGCTGTATTCCCCTCCATCG |
| 16 | mActin-R | CCAGTTGGTAACAATGCCATGT |

Instrumentation

Protein quantification by bicinchoninic acid assay (BCA) and TR-FRET measurements were performed on multi-mode microplate reader SpectraMax iD5 (Molecular Devices LLC). Protein concentration and OD600 measurements were measured by Nanodrop One$^{C}$ Microvolume UV-Vis Spectrophotometer (ThermoFisher). Cell lysis was performed using Branson Ultrasonic Probe Sonicator (model 250) or Fisherbrand Sonic Dismembrator (model 120). Fluorescence and chemiluminescence imaging were performed using Azure Imager c600 or 600 (Azure Biosystems, Inc., Dublin, CA). Protein purification and analytical SEC was performed using an ÄKTA pure 25 equipped with F9-R fraction collector, C9n conductivity monitor, and computer running UNICORN v6.3.2.89 (GE Healthcare). Reverse-phased HPLC for peptide fractionation and protein purification was performed using an Agilent 1260 Infinity II system. Mass spectrometry for compounds and peptides were performed on a Waters ACQUITY UPLC system equipped with SQ Detector 2 mass spectrometer or a Bruker micrOTOF II LC-MS system. Proteomics data were obtained on Vanquish Neo HPLC system (ThermoFisher) connected in line to a Orbitrap Fusion Lumos Tribrid or Orbitrap Eclipse Tribrid Mass Spectrometer (both ThermoFisher) within the Mass Spectrometry and Proteomics Resource Laboratory at Harvard University. Intact protein mass spectra were collected using Bruker Impact II q-TOF mass spectrometer coupled to Agilent 1290 HPLC within the Mass Spectrometry and Proteomics Resource Laboratory at Harvard University. Western blotting transfer was performed using Invitrogen iBlot 2 dry blotting system. RT-qPCR was performed using an iQ5 Multicolor Real-Time PCR Detection System (Bio-Rad). Electroporation was performed using Neon electroporation system (ThermoFisher). Flow cytometry was conducted using FACSymphony A3 Lite analyzer (BD). Cell numbers and viability were measured using TC20 automated cell counter (Bio-Rad). Samples were dried using Vacufuge Plus (Eppendorf).

Software

Data was analyzed and visualized using Microsoft Excel (v16.44) and GraphPad Prism (v8.4.3). DNA and protein sequences were analyzed using Geneious (v11.0.3).

Proteomics data was analyzed using Xcalibur Qual Browser (v3.0.63) and Proteome Discoverer (v2.4.1.15). Flow cytometry populations were distinguished using BD FACSDiva (v8.0.1). Images were made using ImageJ (NIH, v1.52q), PyMOL (v2.0), Adobe Photoshop (v21.1.1) and Adobe Illustrator (v24.1).

#### IV. Synthetic procedures

General reagent information

All reactions were performed under air using the indicated method in general procedures. *N,N*-Dimethylformamide (DMF) (Beantown Chemical, catalog no. BT138690) was vigorously purged with argon for 1 h, followed by passage under argon pressure through two packed columns of neutral alumina (Pure Process Technology). Milli-Q (MQ) water was prepared using a Barnstead™ GenPure™ xCAD Plus Ultrapure Water Purification System (Thermo Fisher Scientific). DMSO-$d_6$ was purchased from Cambridge Isotope Laboratories. All other solvents and reagents were purchased from chemical suppliers (Sigma Aldrich, Chem-Impex International, Inc., TCI, Thermo Fisher Scientific, VWR Chemicals BDH®) and were used as received unless otherwise noted. Flash Column Chromatography was performed using silica gel purchased from Silicycle (SilicaFlash® F60, 40–63 µm) with the aid of a CombiFlash® NextGen 300+ Automated Flash Chromatography System (Teledyne ISCO). Organic solutions were concentrated in vacuo using a Buchi rotary evaporator.

General analytical information

New organic compounds (excluding peptides) were characterized by $^1$H NMR and low-resolution mass spectroscopy. NMR experiments were performed on a Bruker 400 MHz instrument at 24 °C. Chemical shifts are reported in parts per million (ppm) relative to residual solvent as an internal reference (DMSO: δ 2.50 ppm). The following abbreviations were used to explain multiplicities: s = singlet, d = doublet, t = triplet, q = quartet, dd = doublet of doublets, m = multiplet. Low-resolution mass spectra were obtained on a Waters ACQUITY UPLC system equipped with SQ Detector 2 mass spectrometer.

For purification of peptides, RP-HPLC analyses were performed on an Agilent 1260 Infinity II HPLC with UV detection (220 nm and 256 nm) using a XB-C18 column (3.6 µm, 4.6 × 250 mm) and an Agilent Prep-C18 column (5 µm, 30 ×100 mm). Linear gradients of MeCN (with 0.1 % TFA, buffer B) in water (with 0.1 % TFA, buffer A) were used for all systems to elute bound peptides. The flow rates were 1 mL/min (analytical) and 20 mL/min (preparative).The purified peptides were characterized by high-resolution mass spectra, which were recorded on a Bruker micrOTOF II LC-MS system.

General Procedure A: Synthesis of Fmoc-GGGF

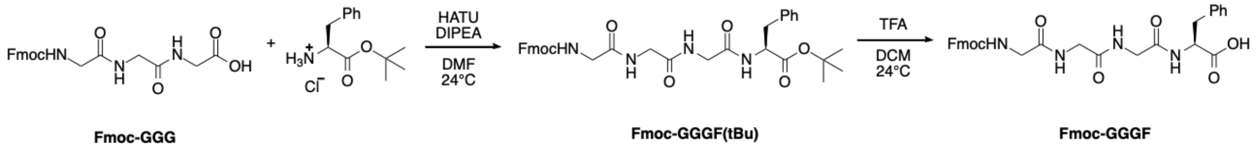


**Fmoc-GGG** (200.0 mg, 0.49 mmol, 1.00 equiv) and L-phenylalanine *tert*-butyl ester hydrochloride (250.7 mg, 0.97 mmol, 2.00 equiv) were dissolved in dry DMF (9.7 mL, 0.05 M). *N,N*-Diisopropylethyl amine (0.42 mL, 2.43 mmol, 5.00 equiv) and HATU (314.1 mg, 0.51 mmol, 1.05 equiv) were added in sequence to the stirred reaction mixture. After stirring at 24 °C for 24 h, the reaction mixture was concentrated with the aid of a rotary evaporator. The crude material was further purified by column chromatography (ISCO, 24 g column, 0–20% MeOH/$CH_2Cl_2$, 40 min gradient). The fractions containing the product were confirmed by TLC, concentrated, and triturated by diethyl ether (3x 2 mL) and then ethyl acetate (3x 2 mL) to afford **Fmoc-GGGF(tBu)** as a white solid (298.8 mg, 0.21 mmol, 44% yield).

**Fmoc-GGGF(tBu)** (60.0 mg, 97.6 μmol, 1.00 equiv) was dissolved in a solution of TFA (0.98 mL) and $CH_2Cl_2$ (0.98 mL). After 3 h, the reaction mixture was concentrated with the aid of a rotary evaporator and triturated by diethyl ether (3x 2 mL). The resulting white solid was dissolved with a mixture of methanol and DCM, concentrated, and dried under high vacuum to yield **Fmoc-GGGF** as a white solid (51.9 mg, 92.3 μmol, 95% yield), which was used directly in the following step without further purification.

**$^1$H NMR** (400 MHz, DMSO-*d*6) δ 12.75 (s, 1H), 8.22–7.99 (m, 3H), 7.89 (d, *J* = 7.5 Hz, 2H), 7.71 (d, *J* = 7.5 Hz, 2H), 7.55 (t, *J* = 6.0 Hz, 1H), 7.42 (t, *J* = 7.4 Hz, 2H), 7.39–7.12 (m, 7H), 4.51–4.33 (m, 1H), 4.33–4.11 (m, 3H), 3.83–3.57 (m, 6H), 3.04 (dd, *J* = 13.8, 5.1 Hz, 1H), 2.88 (dd, *J* = 13.8, 9.0 Hz, 1H).

General Procedure B: Synthesis of $H_2N$-Asn-OMe

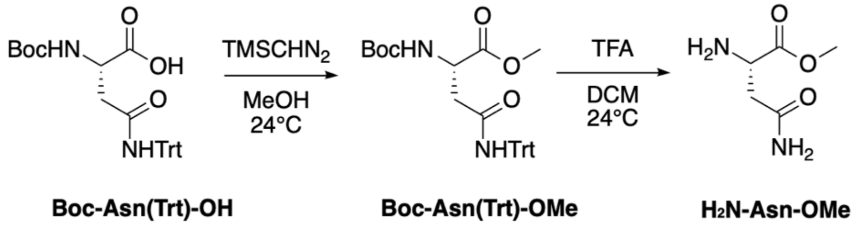


**Boc-Asn(Trt)-OH** (237.3 mg, 0.50 mmol, 1.00 equiv) was dissolved in a mixture of toluene (1.5 mL) and methanol (2.5 mL). (Trimethylsilyl)diazomethane in hexane (1.2 mL, 0.72 mmol, 1.44 equiv) was added dropwise to the stirred reaction until the yellow color persisted, and the reaction was further stirred at 24 °C for 30 min. The reaction mixture was concentrated with the aid of a rotary evaporator and dried under high vacuum to yield **Boc-Asn(Trt)-OMe** as a white solid (223.1 mg, 0.46 mmol, 91% yield).

**Boc-Asn(Trt)-OMe** (50.0 mg, 0.10 mmol, 1.00 equiv) was dissolved in a solution of TFA (1.02 mL) and $CH_2Cl_2$ (1.02 mL). After 3 h, the reaction mixture was concentrated with the aid of a rotary evaporator and triturated by diethyl ether (3x 2 mL). The resulting white solid was dissolved with methanol, concentrated, and dried under high vacuum to yield **$H_2$N-Asn-OMe** as a white solid (27.0 mg, 0.21 mmol, quantitative yield), which was used directly in the following step without further purification.

General Procedure C: Synthesis of $H_2$N-Asn-SMe

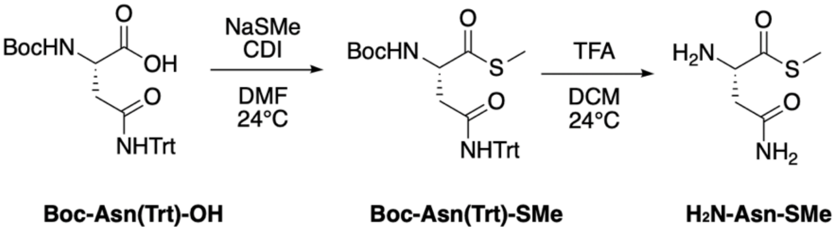


**Boc-Asn(Trt)-OH** (569.5 mg, 1.20 mmol, 2.10 equiv) and CDI (194.58 mg, 1.20 mmol, 2.10 equiv) were dissolved in dry DMF (2.9 mL, 0.2 M) and stirred for about 25 min. NaSMe (40 mg, 0.57 mmol, 1.00 equiv) was added portion-wise to the reaction mixture (3 times, wait 10 min after each addition), and the reaction was further stirred at 24 °C for 1 h. The reaction mixture was diluted with ethyl acetate and transferred to a separatory funnel. The organic layer was first washed with saturated aqueous $NaHCO_3$, extracted by ethyl acetate 3 times, and concentrated with the aid of a rotary evaporator. The resulting solid was again dissolved in ethyl acetate and washed with 1 M HCl and extracted 3 times. The organic layer was dried over $Na_2SO_4$, filtered, and concentrated with the aid of a rotary evaporator. The residue obtained was purified by column chromatography (ISCO, 4 g column, 0–10% MeOH/$CH_2Cl_2$, 30 min gradient) to afford **Boc-Asn(Trt)-SMe** as a white solid (288.0 mg, 0.37 mmol, 65% yield).

**Boc-Asn(Trt)-SMe** (30.0 mg, 59.4 μmol, 1.00 equiv) was dissolved in a solution of TFA (0.59 mL) and $CH_2Cl_2$ (0.59 mL). After 3 h, the reaction mixture was concentrated with the aid of a rotary evaporator and triturated by diethyl ether (3x 2 mL). The resulting white solid was dissolved with methanol, concentrated, and dried under high vacuum to yield **$H_2$N-Asn-SMe** as a white solid (17.4 mg, 95.6 μmol, quantitative yield), which was used directly in the following step without further purification.

General Procedure D: Synthesis of Fmoc-GGGFN(XMe)

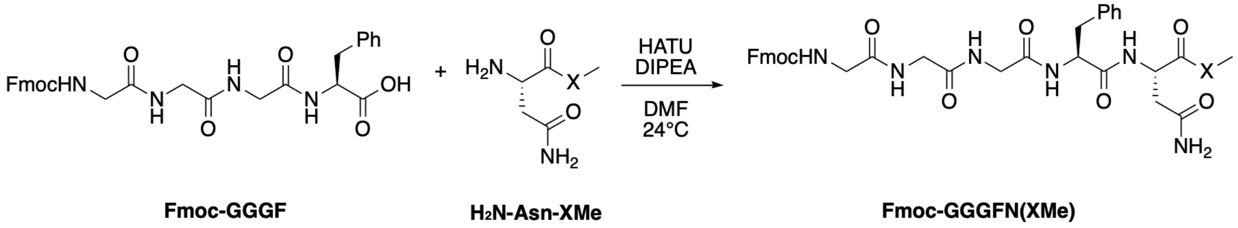

**Fmoc-GGGF** prepared from General Procedure A (1.00 equiv) and **$H_2$N-Asn-XMe** prepared from General Procedure B or C (1.26 equiv) were dissolved in dry DMF (1.0–1.6 mL). *N,N*-Diisopropylethyl amine (5.00 equiv) and HATU (1.05 equiv) were added in sequence to the stirred reaction mixture. After stirring at 24 °C for 24 h, the reaction mixture was concentrated with the aid of a rotary evaporator and triturated by diethyl ether (2x 2 mL) and then ethyl acetate (2x 2 mL). The resulting white solid was dissolved and flushed with DMF, concentrated, and further purified by preparative HPLC to afford **Fmoc-GGGFN(XMe)**.

**Fmoc-GGGFN(OMe)**

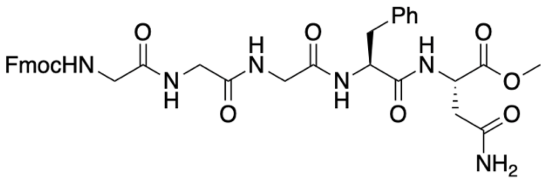


The title compound was prepared according to General Procedure D from **Fmoc-GGGF** (45.4 mg, 81.3 µmol, 1.00 equiv), **$H_2$N-Asn-OMe** (15.0 mg, 0.10 mmol, 1.26 equiv), N,N-diisopropylethyl amine (71 µL, 0.41 mmol , 5.00 equiv), and HATU (32.4 mg, 85.3 µmol, 1.05 equiv) in dry DMF (1.6 mL). After purification by preparative HPLC (Agilent 5 Prep-C18 OBD, 5 µm, dimensions 100 mm x 30.0 mm, 95–5% MQ water/ACN, 5 min gradient, 250 nm detection, 24 mL/min flow rate), the title compound, **Fmoc-GGGFN(OMe)**, was obtained as a white solid (55.8 mg, 43.0 µmol, 53% yield).

**$^1$H NMR** (400 MHz, DMSO-*d*6) δ 8.45 (d, *J* = 7.7 Hz, 1H), 8.22–7.97 (m, 3H), 7.89 (d, *J* = 7.5 Hz, 2H), 7.70 (d, *J* = 7.5 Hz, 2H), 7.54 (t, 1H), 7.45–7.14 (m, 10H), 6.93 (s, 1H), 4.74–4.44 (m, 2H), 4.38–4.04 (m, 3H), 3.81–3.48 (m, 9H), 3.00 (dd, *J* = 13.8, 4.4 Hz, 1H), 2.74 (dd, *J* = 13.7, 9.7 Hz, 1H), 2.58 (dd, *J* = 15.8, 6.1 Hz, 1H), 2.47–2.42 (m, 1H).

**Fmoc-GGGFN(SMe)**

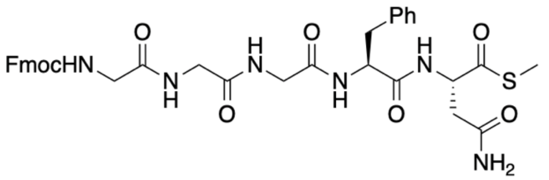


The title compound was prepared according to General Procedure D from **Fmoc-GGGF** (17.5 mg, 31.3 µmol, 1.00 equiv), **$H_2$N-Asn-OMe** (6.4 mg, 39.5 µmol, 1.26 equiv), N,N-diisopropylethyl amine (27 µL, 0.16 mmol , 5.00 equiv), and HATU (12.5 mg, 32.9 µmol, 1.05 equiv) in dry DMF (1.0 mL). After purification by preparative HPLC (Agilent 5 Prep-C18 OBD, 5 µm, dimensions 100 mm x 30.0 mm, 95–5% MQ water/ACN, 5 min gradient, 250 nm detection, 24 mL/min flow rate), the title compound, **Fmoc-GGGFN(SMe)**, was obtained as a white solid (4.8 mg, 6.8 µmol, 22% yield).

**$^1$H NMR** (400 MHz, DMSO-*d*6) δ 8.73 (d, *J* = 8.0 Hz, 1H), 8.26–7.96 (m, 3H), 7.89 (d, *J* = 7.5 Hz, 2H), 7.71 (d, *J* = 7.4 Hz, 2H), 7.54 (t, *J* = 6.1 Hz, 1H), 7.46–7.15 (m, 10H), 6.96 (s, 1H), 4.77 (q, *J* = 7.0 Hz, 1H), 4.69–4.46 (m, 1H), 4.34–4.10 (m, 3H), 3.83–3.47 (m, 6H), 3.14 (dd, *J* = 13.7, 3.9 Hz, 1H), 2.76 (dd, *J* = 13.8, 10.1 Hz, 1H), 2.60 (dd, *J* = 15.8, 5.6 Hz, 1H), 2.47–2.34 (m, 1H), 2.18 (s, 3H).

General Procedure E: Fmoc solid-phase peptide synthesis

Peptides were synthesized by Fmoc-SPPS on 2-chlorotrityl chloride (2-CTC) resin (loading 1.0–2.0 mmol/g, 0.25 mmol scale). 2-CTC resin was swelled in DMF for 1 h. The first Fmoc-amino acid (4 equiv) activated with DIPEA (8 equiv) in DMF was doubly coupled to the 2-CTC resin (loading 1.0–2.0 mmol/g, 0.25 mmol scale) for 30 min. Fmoc-deprotection was carried out with 20% piperidine in DMF (10 min × 2). Fmoc-amino acids (1 mmol in 5 mL of DMF, 4 equiv) were activated with HATU (1 mmol in 5 mL of DMF, 4 equiv) and DIPEA (2 mmol in 5 mL of DMF, 8 equiv) for 5 min and allowed to couple for 30 min with constant shaking. Boc-protected amino acids were employed as last amino acids for the synthesis of global protected peptides. The resulting resins were washed with DMF (× 3) and DCM (× 3), and dried under vacuum.

General Procedure F: Cleavage from the resin

The peptide was cleaved using Hexafluoroisopropanol (HFIP)/DCM (1:4 v/v) cocktail for 30 min. The cleavage mixture was filtered, and the resin was washed with DCM. The combined solutions were evaporated by $N_2$ bubbling to minimum volume, and the crude peptides were dissolved in MeCN:water (1:1) and further diluted to around 25% MeCN with water and lyophilized. The crude peptides were used for next step without purification.

General Procedure G: Synthesis of Target Peptides-cN

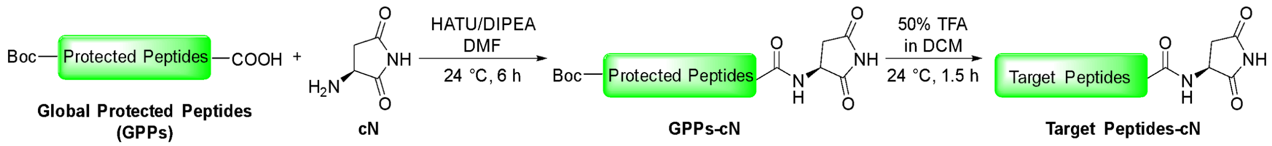


Crude **GPPs** (0.100 mmol, 1.00 equiv) and **cN** (0.30 mmol, 3.00 equiv) were dissolved in DMF (2.0 mL, 0.050 M). DIPEA (0.600 mmol, 6.00 equiv) and HATU (0.3 mmol, 3.00 equiv) were added sequentially to the stirred reaction mixture. After stirring at 24 °C for 6 h, DMF was removed with the aid of a rotary evaporator.

The crude material was dissolved in 2 mL of 50% TFA solution (in DCM, with 3% triisopropylsilane (TIPS), additional 3% ethanedithiol was used for the peptides containing Met). After stirring at 24 °C for 1.5 h, the obtained mixture was concentrated by $N_2$ bubbling to minimum volume, after which 1 mL of DMF was added. The mixture was purified by semi-preparative HPLC to yield **Target peptides-cN**.

### V. NMR Spectra and LC-MS Traces

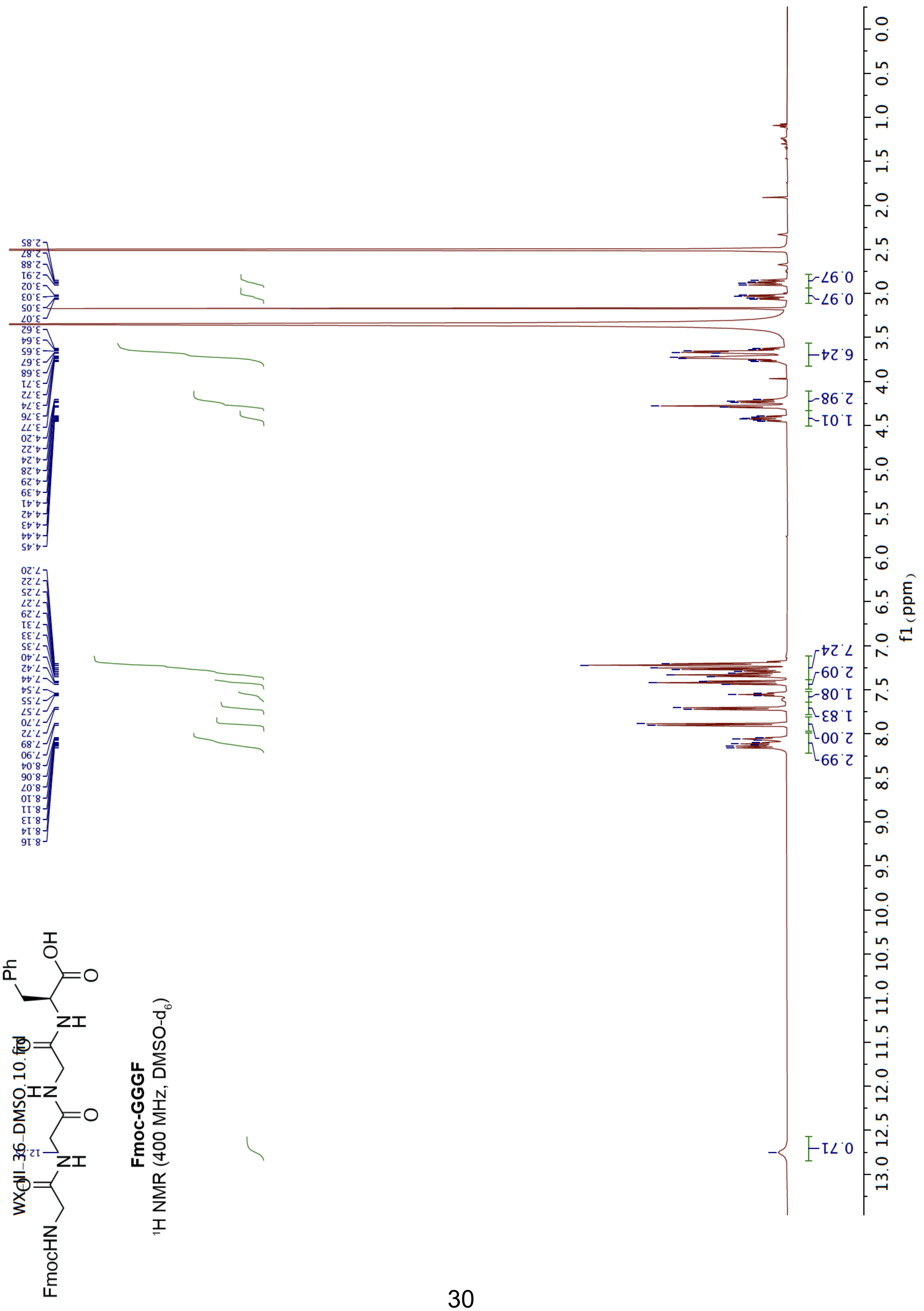

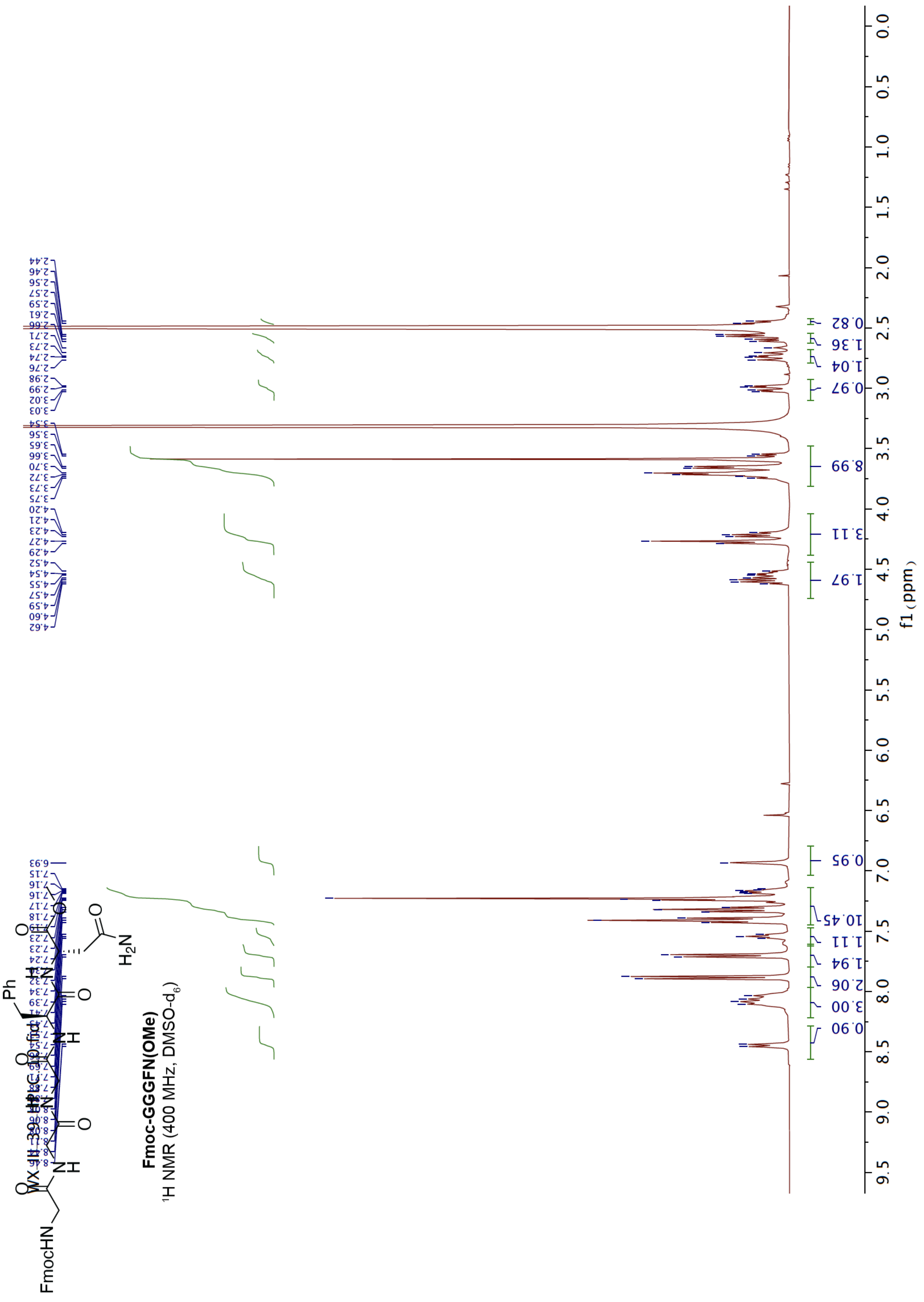

**Fmoc-GGGFN(OMe)**
$^1$H NMR (400 MHz, DMSO-$d_6$)

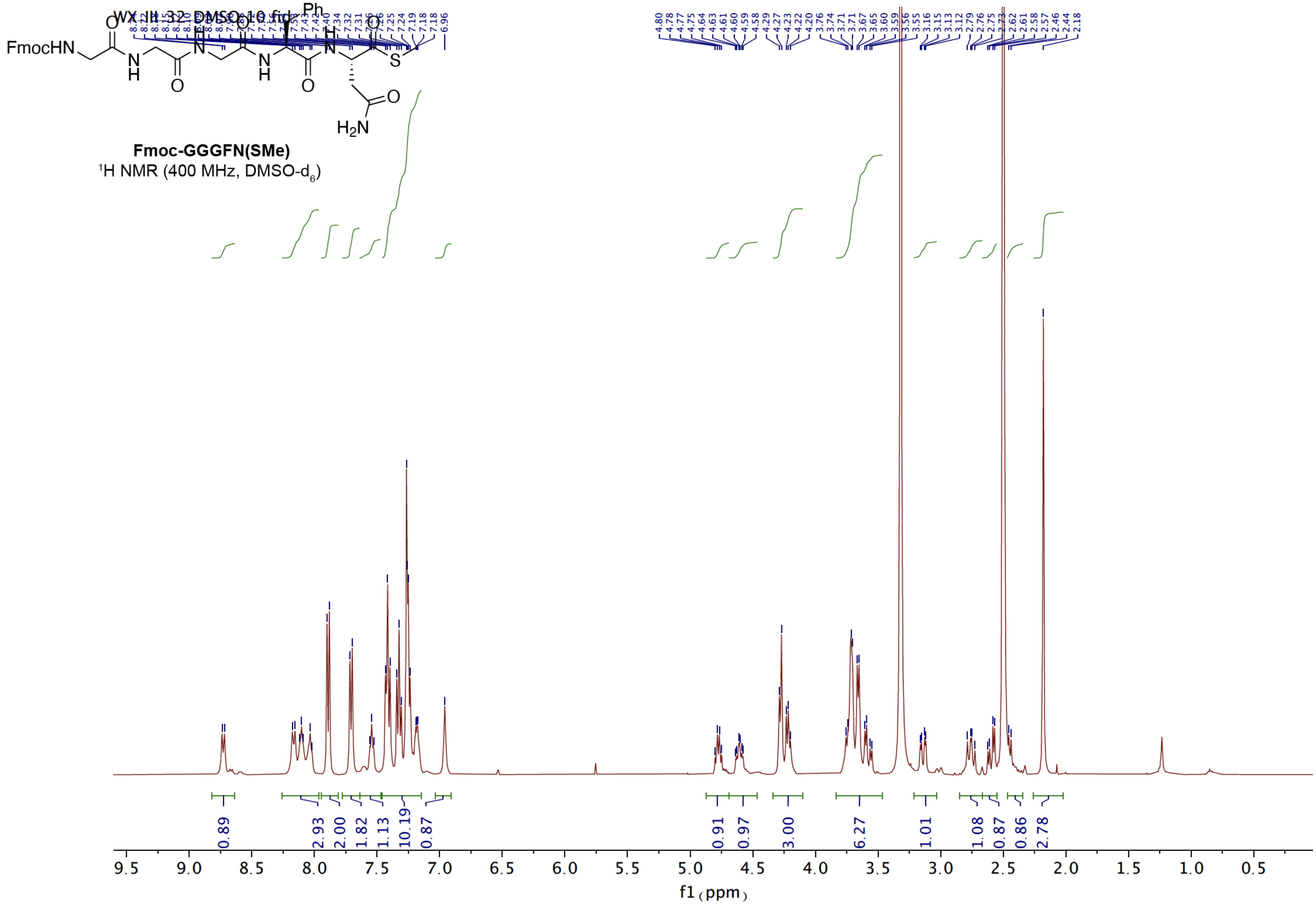
Fmoc-GGGFN(SMe)
$^1$H NMR (400 MHz, DMSO-$d_6$)
f1 (ppm)

**GGGVYPisoDHA.** HPLC-MS (BEH-C18 column: 1.7 µm, 130 Å, 2.1 × 150 mm at 220 nm) of GGGVYPisoDHA (obs. 872.3778; calc. for [M + H]+ 872.3897).

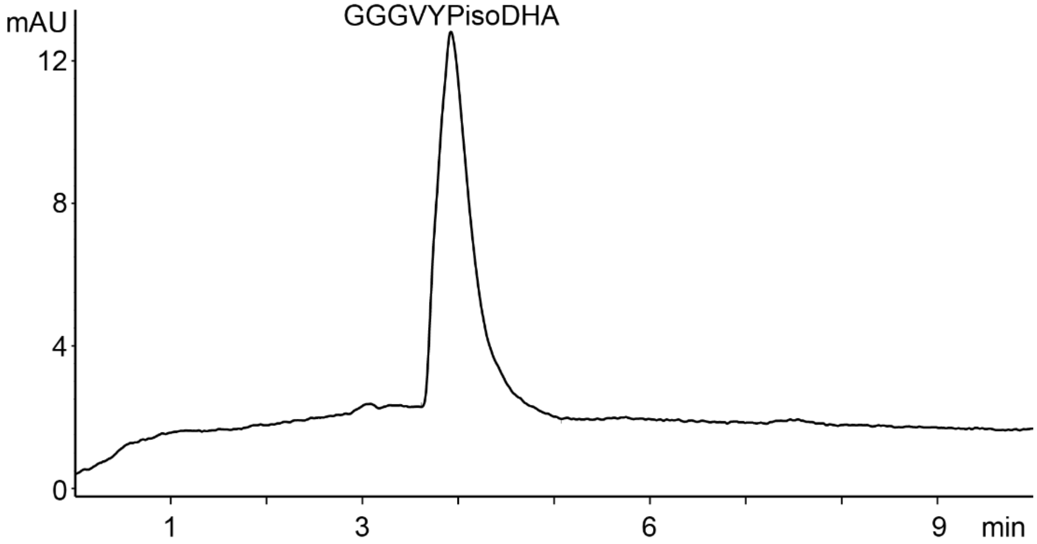


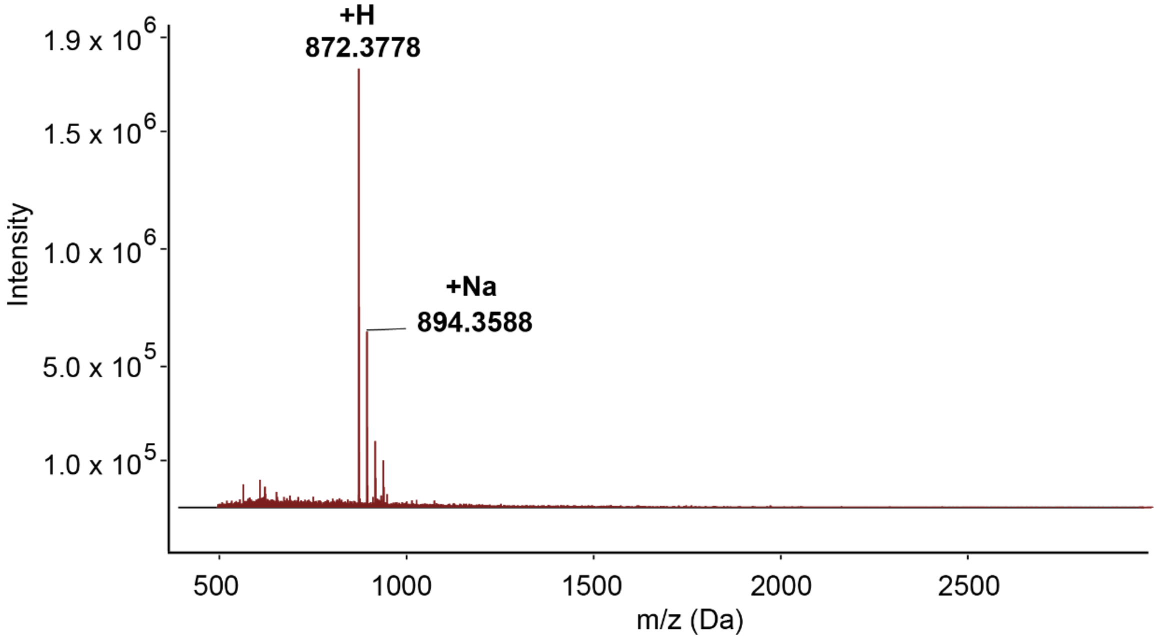

**PFQYKcN.** HPLC-MS (BEH-C18 column: 1.7 µm, 130 Å, 2.1 × 150 mm at 220 nm) of PFQYKcN (obs. 778.3684; calc. for [M + H]+ 778.3883).

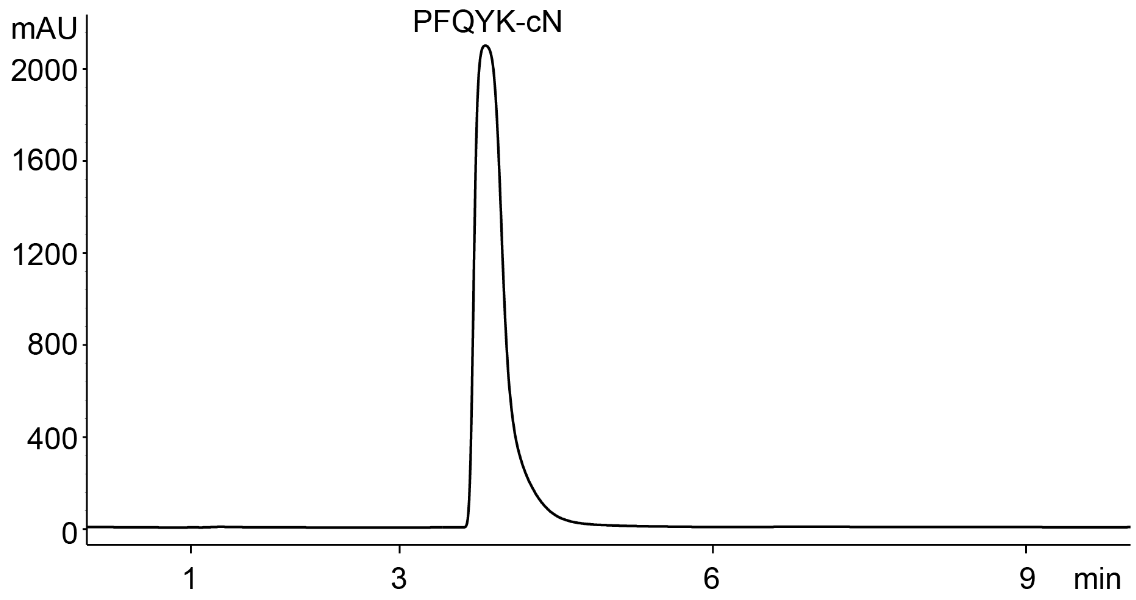


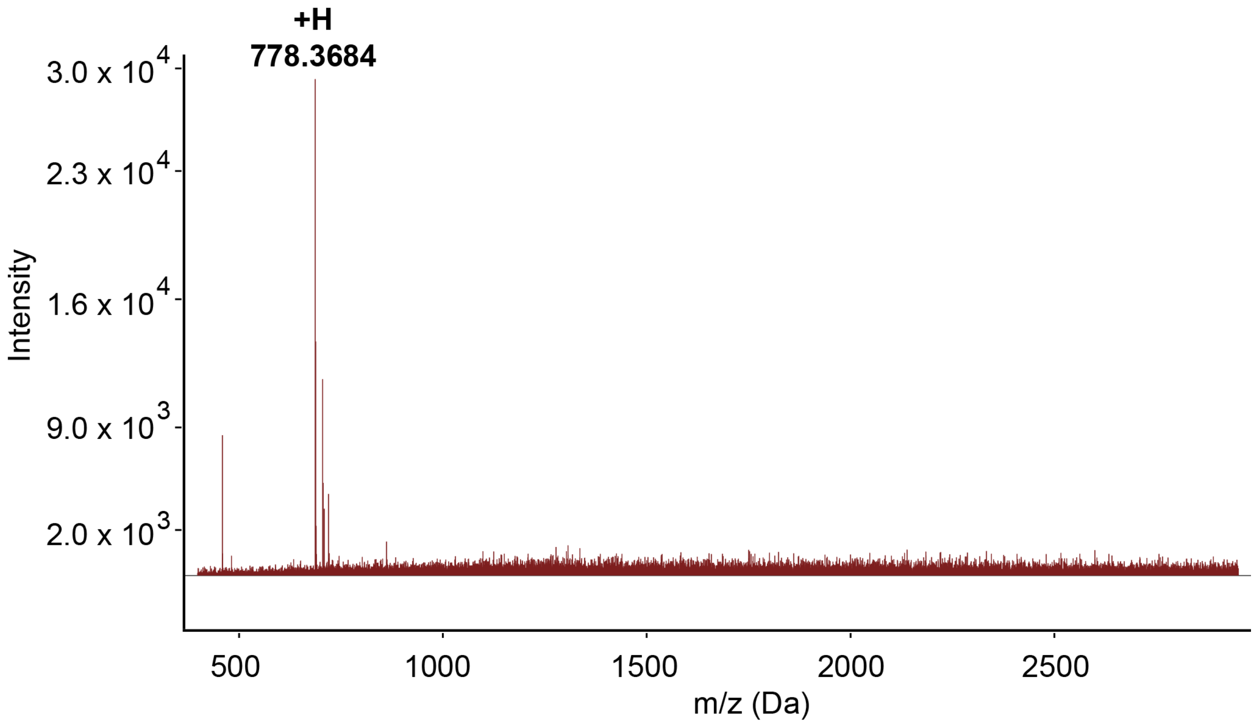

**PFQYKN(OMe)**. HPLC-MS (BEH-C18 column: 1.7 µm, 130 Å, 2.1 × 150 mm at 220 nm) of PFQYKN(OMe) (obs. 810.3755, 832.3611; calc. for $[M + H]^+$ 810.4145, $[M + Na]^+$ 832.3964).

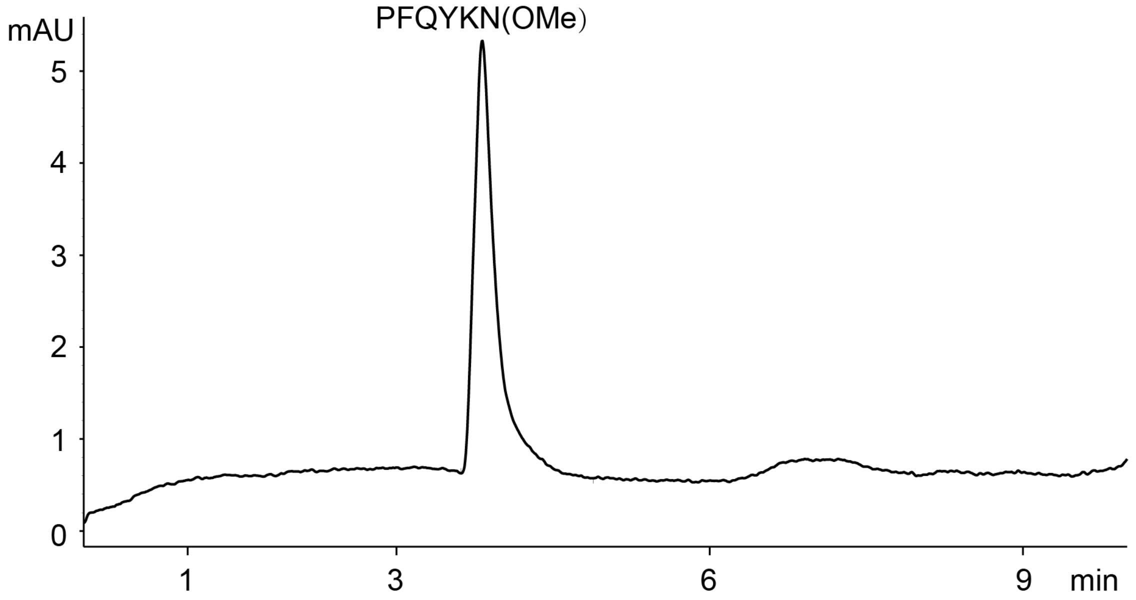


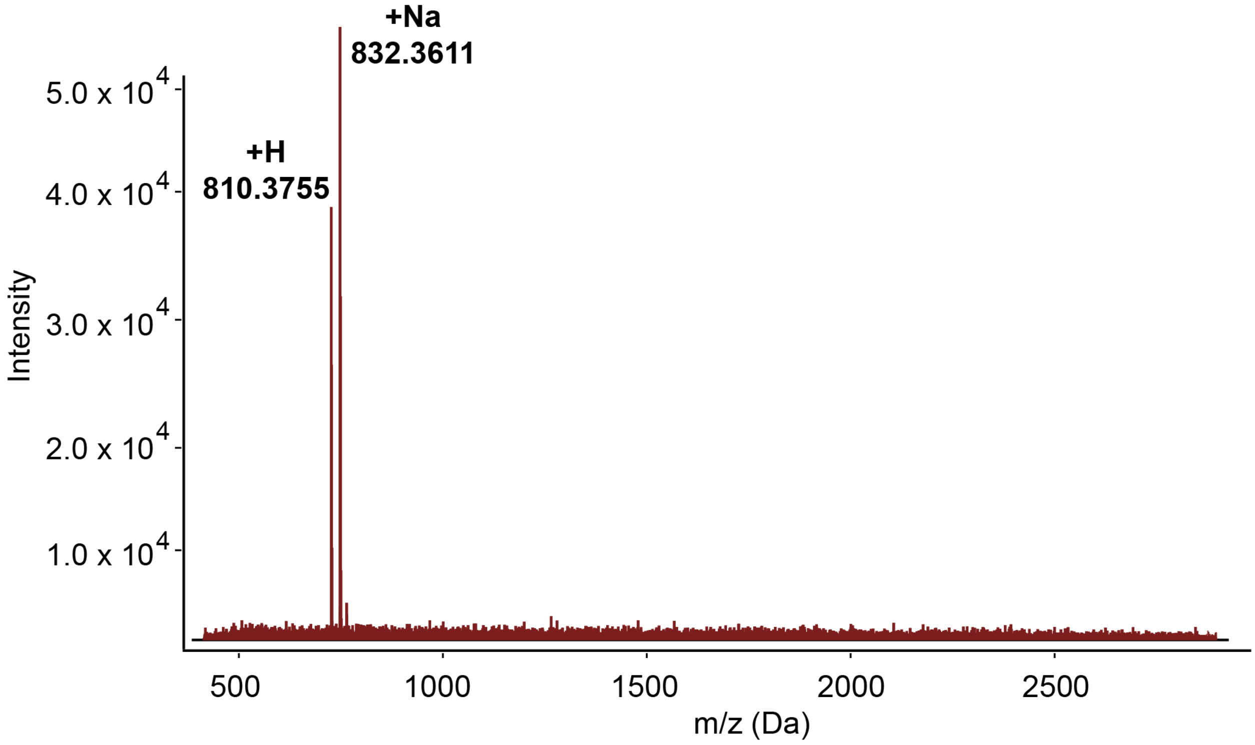
